# Spatial regulation of the glycocalyx component Podocalyxin is a switch for pro-metastatic function

**DOI:** 10.1101/2022.11.04.515043

**Authors:** Alvaro Román-Fernández, Mohammed A. Mansour, Fernanda G. Kugeratski, Jayanthi Anand, Emma Sandilands, Laura Galbraith, Kai Rakovic, Eva C. Freckmann, Erin M. Cumming, Ji Park, Konstantina Nikolatou, Sergio Lilla, Robin Shaw, David Strachan, Susan Mason, Rachana Patel, Lynn McGarry, Archana Katoch, Kirsteen J. Campbell, Colin Nixon, Crispin J. Miller, Hing Y. Leung, John Le Quesne, James C. Norman, Sara Zanivan, Karen Blyth, David M. Bryant

**Affiliations:** Institute of Cancer Sciences, University of Glasgow, Glasgow G61 1QH; The CRUK Beatson Institute, Glasgow G61 1BD, United Kingdom; Cancer Biology and Therapy Lab, Division of Human Sciences, School of Applied Sciences, London South Bank University, London SE1 0AA, UK; Biochemistry Division, Department of Chemistry, Faculty of Science, Tanta University, Tanta 31527, Egypt

## Abstract

The glycocalyx component and sialomucin Podocalyxin (PODXL) is required for normal tissue development by promoting apical membranes to form between cells, triggering lumen formation. Elevated PODXL expression is also associated with metastasis and poor clinical outcome in multiple tumour types. How PODXL presents this duality in effect remains unknown. We identify an unexpected function of PODXL as a decoy receptor for Galectin-3 (GAL3), whereby the PODXL-GAL3 interaction releases GAL3 repression of integrin-based invasion. Differential cortical targeting of PODXL, regulated by ubiquitination, is the molecular mechanism controlling alternate fates. Both PODXL high *versus* low surface levels occur in parallel subpopulations within cancer cells. Orthotopic intraprostatic xenograft of PODXL-manipulated cells or those with different surface levels of PODXL define that this axis controls metastasis *in vivo*. Clinically, interplay between PODXL-GAL3 stratifies prostate cancer patients with poor outcome. Our studies define the molecular mechanisms and context in which PODXL promotes invasion and metastasis.

## Introduction

The cellular glycocalyx is comprised of a network of glycoproteins and proteoglycans that coat all cells and that modulates cell interactions with the extracellular matrix (ECM) (*1*). An altered, bulky glycocalyx in cancer cells contributes to tumour progression and metastasis by mechanical regulation of cell-ECM interactions (*2-5*). A conundrum in cancer cell glycocalyx constituents influencing ECM interaction is that a number of these are mucins, such as MUC1 or Podocalyxin (PODXL), which in normal tissues are not localised to ECM-abutting domains but are instead apically localised and required for apical domain function (*6, 7*). This suggests a duality in mucin function, promoting normal tissue development at the apical domain or invasion and metastasis when alternately located to ECM-abutting membranes.

The duality of glycocalyx-associated mucins is particularly evident for the sialomucin PODXL. PODXL’s role in promoting apical lumen formation across tissues can be converted to inducing collective apical membrane-driven invasion by interfering with signals that control PODXL removal from ECM-abutting membranes (*7*). Such ‘inverted polarity’, apical proteins localised at the periphery of clusters of epithelial cells rather than at the lumen, is a defining feature of micropapillary cancers that are associated with poor clinical outcome (*8, 9*). Apical membrane-driven inverted polarity has been proposed as a mechanism for metastasis in colorectal carcinoma (*10*). Indeed, elevated PODXL levels are an independent prognostic indicator of disease aggressiveness in several cancers (*11-17*). Germline mutations in *PODXL* result in alternate phenotypes; some PODXL mutations are associated with focal segmental glomerulosclerosis and nephrotic syndrome affecting apical membrane function of PODXL in the kidney (*18, 19*), whereas other *PODXL* variants are associated with familial prostate cancer aggressiveness (*12*). This highlights that the functional contribution of mucins to the glycocalyx, whether it be to normal tissue function or disruption to this function and induction of metastatic features, must be regulated in a context-specific fashion, such as alternate subcellular targeting to the apical domain or to the ECM-abutting surface.

That PODXL is a transmembrane protein and that its elevated expression is associated with poor clinical outcome in tumours has led to suggestion of it as a targetable biomarker of metastasis (*20-23*). Given that the function of PODXL in normal tissues is also required, it is essential to identify the tumour-specific contexts regulating PODXL function. PODXL expression alone modestly stratifies outcome in some cancers but becomes a potent predictor of poor outcome when comparing cortical to intracellular levels of PODXL (*11, 23-28*). Moreover, tumour-restricted glycosylation patterns have allowed the development of tumour cell-selective therapeutic anti-PODXL antibodies (*29*). This emphasises that mucins such as PODXL require contextual modulation to form part of the tumour-associated, pro-metastatic bulky glycocalyx. Here, we determine the molecular mechanisms that control the switch of PODXL towards a pro-metastatic glycocalyx component. We identify an unexpected function of PODXL as a decoy receptor that relieves the invasion-inhibiting effect of the glycocalyx component Galectin-3 and demonstrate that the levels of this switch identify prostate cancer patients with high metastasis and poor outcome.

## RESULTS

### PODXL promotes invasive tunnel formation inside the ECM

We identified breast and prostate cancer cell lines with high PODXL expression, as PODXL has been associated with invasion and metastasis in these cancers. Non-tumorigenic prostate lines PREC-LH (*30*) (Fig. 1A) and RWPE1 cells (*31*) (Fig. 1B) minimally expressed *PODXL* mRNA. In contrast, highly metastatic prostate PC3 (*32*) and breast MDA-MB-231 (*33*) lines displayed highest *PODXL* mRNA with minimal change to gene copy number (dashed lined square, Fig. 1A; Supplementary Table 1). We therefore explored PODXL function in PC3 and MDA-MB-231 cells.

**Figure 1.**
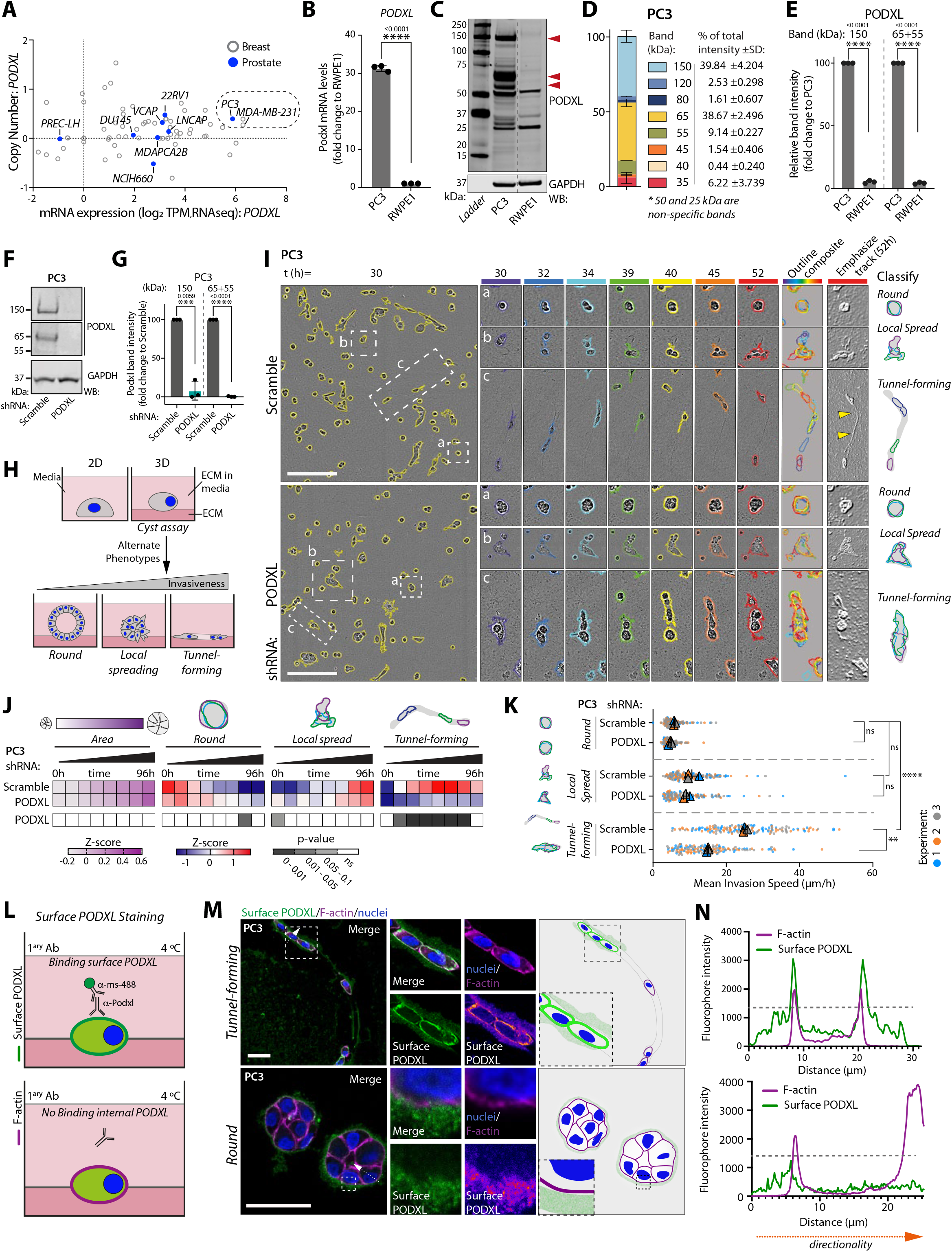
PODXL promotes formation of invasive tunnels in the ECM. (A) *PODXL* mRNA vs copy number. Prostate, blue; breast, grey. Data, CCLE. (B) *PODXL* mRNA, RWPE1 vs PC3. Normalised to RWPE1, n=3 independent isolations/reactions. (C) Western blot, RWPE1 and PC3 for PODXL and GAPDH (loading control). Arrowheads, major bands. Dashed line, non-adjacent lanes. (D-E) Relative contribution to total PODXL levels and relative PODXL levels in (C). Major bands, 150kDa and 55-65kDa. 50kDa and 25kDa bands, non-specific. Normalised to PC3, n=3 independent lysate preparations. (F-G) Western blot and quantitation, PC3 expressing Scramble or *PODXL* shRNA for PODXL and GAPDH (loading control). Normalised to Scramble, n=3 independent lysate preparations. (H) Schema, alternate 3D phenotypes. (I) Phase image, PC3 from (F). Cysts, yellow outline. Example of (a) round, (b) local-spreading, and (c) tunnel-forming objects shown. Outlines coloured by time in single frames and overlayed. Inverted image, ECM-tunnels. Schema, alternate 3D behaviours. (J) Heatmap, Area and phenotype classification of (I). Z-score-normalised values (blue to red). Heatmap, p-values (greyscale) compared to Scramble, Student’s t-test and Cochran-Mantel-Haenszel, Bonferroni adjusted. n=3 independent experiments, 2-3 technical replicates/condition, 254-518 cysts/condition/experiment. (K) Mean invasion speed (μm/h) of (I). n=3 independent experiments, 38-68 cysts/condition/experiment. (L) Schema, surface PODXL immunostE and IHC Staining aining. (M) Confocal images and schema, surface PODXL (green, or Fire LUT), F-actin (magenta) and nuclei (blue) in tunnel-forming or round 3D PC3. Dotted arrow, described in (N). (N) Intensity profiles for surface PODXL (green) and F-actin (magenta) from (M). Dotted line, threshold for background labelling outside cell boundary. Scale bars, (I) 300 or (M) 50 μm. p-values, unpaired Student’s 2-tailed t-test; ns, not significant, **P ≤ 0.005, ***P ≤ 0.0005, ****P ≤ 0.0001. Bar graphs, mean ± SD. Superplots, mean ± SD; circles, technical replicates; triangles, average/experiment.

The predicted molecular weight of PODXL is 58 kDa but likely due to glycosylation (*34, 35*) several bands of PODXL were detected in western blots in both cell lines, most prominently ∼150 kDa and 55-65 kDa (Fig. 1C-E, Fig. S1A). Depletion of PODXL via shRNA (Fig. 1F,G; Fig. S1A,B) or sgRNA-mediated knockout (Fig. S1C,D) largely abolished both PODXL bands.

We examined the phenotype of PODXL depletion. PODXL-depleted cells were impaired in collective cell invasion specifically when ECM was involved. In scratch-wound assays wherein cell monolayers were plated on, and overlayed with, ECM control cells from PC3 and MDA-MB-231 cell lines invaded via leader cells forming tunnels that were used by follower cells to invade (Fig. S1E-K, yellow arrowheads; Supplementary Movies 1-2), similar to previous observations (*36, 37*). PODXL-depleted cells failed to form leader cell-induced tunnels in both cell types (Fig. S1F,H), resulting in decreased invasion of the overall population (Fig. S1G, I), suggesting a common motility defect. In contrast to invasion defects in 3D, PODXL-depleted PC3 cells displayed no defect in 2D migration in scratch-wound migration assays without ECM addition (Fig. S1J-M). No defect was detected in PC3 proliferation in either in 2-dimensional (2D) culture or when embedded into ECM gels to form 3-dimensional (3D) acini (Fig. S1N,O). PODXL is therefore required for invasion into ECM.

To determine whether PODXL was essential for tunnel-inducing cells or movement of follower cells into tunnels we embedded single PC3 cells into ECM to induce 3D acinus formation (Figure 1H). We examined these using wide-field multi-day timelapse imaging, allowing detection of how clonal outgrowth of a single cell can progress to alternate 3D phenotypes (*38, 39*). By overlaying the outlines of acini over defined time intervals into a single image (outline composite), three phenotypes that occurred in parallel in the control PC3 cells (Scramble, non-targeting shRNA) could be observed: non-motile, round acini (Round); those that only invade locally (Local spreading); and those that move back and forward through a seeming conduit in the ECM (Tunnel-forming) that was reminiscent of leader cells at the front of invasion assays (Fig. 1I,J; Fig. S1F; Supplementary Movie 3). We emphasised the ECM in phase-contrast images (‘Emphasize track’; see methods), which revealed a tunnel in the ECM.

We developed a method for quantitative assessment of these phenotypes over time (Fig. S2A). While round and local-spreading phenotypes could potentially be distinguished by their shape-changes at a given static timepoint, tunnel-forming cells shared many of these shapes but instead moved along conduits in the ECM. By creating outline composites of cells imaged every hour for several days into 12h time intervals and converting composites into single objects, we could train a Gentle Fast Boosting machine learning model with high fidelity (92-98% accuracy) to user defined classification to detect the three aforementioned phenotypes. The application of this method to large-scale parallel live imaging of thousands of acini allowed robust statistical analysis of phenotypes, which could be presented in heatmaps that displayed how parameters of interest changed in 12h intervals along with statistical analysis across multiple replicates and experiments.

Quantitative assessment revealed that while PODXL-depleted PC3 cells could display modest elongation, they were devoid of the tunnel-forming phenotype, the local-spreading phenotype was unchanged, and switched to a round phenotype modestly upon PODXL knockdown (shRNA, Fig. 1I,J; Supplementary Movie 3) or significantly upon PODXL knockout (sgRNA; Fig S2B,C). Modest effects were observed on total area (Fig. 1I,J; S2B,C), corroborating a lack of effect of PODXL depletion on proliferation in 3D (Fig. S1N). When MDA-MB-231 cells are embedded into ECM, these initially become highly motile and elongated, also moving back and forwards through apparent ECM tracks before developing into a branched multicellular network (Fig. S2D; Supplementary Movie 4). Quantitative analysis revealed this as an increase in local-spreading and tunnel-forming activity over time (Fig. S2E). Upon PODXL depletion, MDA-MB-231 cells at early timepoint lost this initial motility and instead switched to a robust local-spreading phenotype (Fig. S2D,E; Supplementary Movie 4). The analysis of tunnel-forming objects in MDA-MB-231 cells upon PODXL depletion was complicated by the extreme elongation of non-motile cells in this condition, resulting in a lack of significant reduction in the tunnel-forming phenotype (Fig. S2D,E). Therefore, we directly examined the mean invasion speed of elongating PC3 and MDA-MB-231 cells in 3D culture, rather than using tunnel-forming classification as an indirect proxy for motility. In PC3 acini, PODXL depletion resulted in a decreased mean invasion speed specifically in the Tunnel-forming population of acini (Fig. 1K; Fig. S2F). A reduction in mean invasion speed was also observed in MDA-MB-231 cells upon PODXL depletion (Fig. S2G). This suggests that while PODXL depletion can have varying effects on acinus shape, across cell types there is a consistent reduction of motility in a 3D context. This is further supported by a common reduction of invasion in orthogonal ECM invasion assays in both PC3 and MDA-MB-231 cells (Fig. S1F-I). As this motility in PC3 cells results in formation of a tunnel, we therefore use tunnel formation in PC3 cells as method to identify movement over time from static images.

Expression of extracellularly GFP-tagged PODXL (isoform A; hereforth described as wild-type, WT) in PC3 cells stimulated the tunnel-forming phenotype to occur earlier at the expense of local spreading compared to GFP alone (GFP-PODXL; Fig. S3A-D). Intriguingly, the round phenotype was unaffected by PODXL depletion or overexpression despite stable viral manipulation of both conditions. Labelling of endogenous surface PODXL in non-permeabilized acini revealed ECM-adjacent labelling in tunnel-forming, but not round, acini (Fig. 1L-N). In GFP-PODXL-expressing acini, anti-GFP antibody staining to extracellular GFP revealed heterogeneity in surface GFP-PODXL presence, despite similar levels of total GFP-PODXL expression (Fig. S3E). This suggests that the specificity of PODXL effects on tunnel formation is not due to lack of PODXL expression in other phenotypes (round, local-spreading), but rather the existence of a specific context that facilitates cortical PODXL localisation to promote motility in the tunnel-forming population.

### Differential cortical localisation of PODXL controls invasiveness

We examined the context in which PODXL can promote invasive tunnel formation and whether this was related to cortical presentation of PODXL. Three rounds of serial sorting for endogenous surface PODXL levels allowed recovery of cell populations with surface PODXL at high or low levels (Fig. 2A-C). Strikingly, cells with altered surface levels of PODXL displayed different frequencies of local-spreading and tunnel-forming 3D phenotypes; low-surface PODXL cells lost the ability to form tunnels while high-surface PODXL cells were highly tunnel-forming (Fig. 2D,E; Supplementary Movie 5). Consequently, high-surface PODXL cells formed elongated tunnels that allowed rapid invasion of the cell population in ECM invasion assays (Fig. S4A, yellow arrowheads; Supplementary Movie 6), while wounded monolayers of low-surface PODXL cells lacked invasive chains and were poorly invasive (Fig. S4A-C). No proliferation differences were detected between high-surface and low-surface PODXL expressing cells in either 2D or 3D (Fig. S4D,E), similar to effects of PODXL depletion on parental cells (Fig. S1N,O).

**Figure 2.**
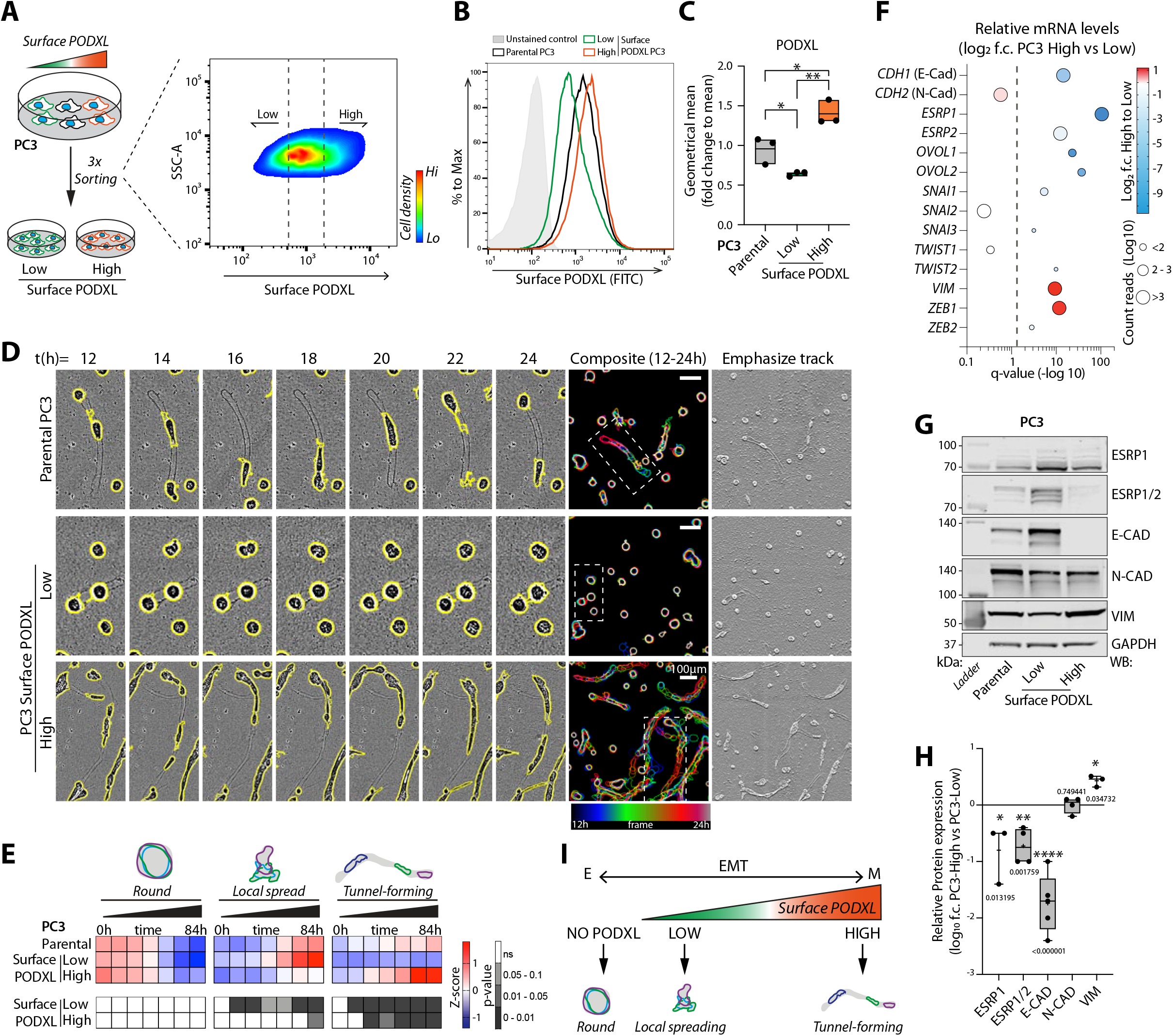
Tunnel-forming invasion is dependent on cortical PODXL. (A) Schema, isolation of PC3 subpopulations with Low or High surface PODXL levels. (B-C) Representative plot and geometrical mean from flow cytometry using anti-extracellular PODXL in cells from (A). Normalised to mean of PC3. Dots, experiments; midline, mean; boundaries, min and max values; n=3 independent experiments. (D-E) Phase images and quantitation of PC3 from (A). Cysts, yellow outlines. Outlines coloured by time in single frames and overlayed. Scale bar, 100 μm. Inverted image, ECM-tunnel (Emphasize track). Heatmap of relative levels of Round, Spread and Tunnel-forming PC3, Z-score normalised. n=4 independent experiments, 3-6 technical replicates/condition, 274-942 cysts/ condition/experiment. (F) Bubble plot, mRNA expression of EMT-related genes. X-axis, q-value (-log10), grey dashed line, significance (p=0.05); colour, PC3-high vs PC3-low mRNA expression fold change value (log2, blue to red); bubble size, average of normalised count reads of each transcript (Log10); n=4 independent RNA isolations/condition. (G) Western blot and quantitation, PC3 parental or sorted subpopulations for EMT-related markers (ESRP1/2, E-CAD, N-CAD, VIM) or GAPDH (loading control). High-surface versus low-surface PODXL levels; intensity fold change (log10). Box-and-whiskers: dots, replicates; +, mean; midline, median; boundaries, quartiles; n=3-5 independent experimental isolations of cell lysates. (I) Schema. Relationship between PODXL, EMT and phenotype. (C) p-values, unpaired Student’s 2-tailed t-test; (H) p values, multiple unpaired t-test, Welch’s correction. ns, not significant,*P ≤ 0.05, **P ≤ 0.005, ***P ≤ 0.0005, ****P ≤ 0.0001. Heatmap, p-values, Cochran-Mantel-Haenszel and Bonferroni adjusted, compared to control, greyscale as indicated.

The contrast in surface levels between high-surface versus low-surface PODXL cells was not reflected in major mRNA or total protein level changes of either PODXL or its known major binding partners (Ezrin, *SLC9A3R1*/NHERF1, *SLC9A3R2*/NHERF2); only a small reduction in *EZRIN* mRNA, but not protein, occurred (Fig. S5A-C) while total PODXL protein levels showed a modest difference between high-surface versus low-surface stable lines (1.68-fold change; Fig. S5B,C). Robust overexpression of GFP-PODXL (Fig. S4F,G) in low-surface stable lines did not increase tunnel formation (Fig. S4H), similar to a lack of effect in the round population in GFP-PODXL-overexpressing parental PC3 cells (Fig. S3A-D). This supports the notion of a cellular context that promotes surface PODXL and invasion, rather than a difference in PODXL complex expression between high-versus low-surface PODXL cells.

Analysis of RNA-seq of parental, high-surface and low-surface PODXL lines revealed significant enrichment of pathways associated with epithelial-mesenchymal transition (EMT) between sublines (Fig. S5D). Low-surface PODXL cells presented an epithelial profile with enrichment of epithelial transcripts (*CDH1, ESRP1/2*), whereas high-surface PODXL cells presented a mesenchymal profile (*VIM, ZEB1*; Fig. 2F), data confirmed at protein level (Fig. 2G,H). This trend was maintained in both 2D and 3D culture (Fig. S5E,F). This suggested that an EMT provides the context that promotes increased cortical levels of PODXL, independent of regulating the total levels of PODXL or major known PODXL binding partners. This raised the possibility that cortical PODXL might simply be a functionally irrelevant passenger event that nonetheless could be used to select for a mesenchymal mode of invasion. However, depletion of PODXL in high-surface PODXL cells attenuated the ability of these to form tunnels and the entire population of cells to invade (Fig. S5G-K). This confirms a functional role of PODXL in mesenchymal-type invasion.

Despite reports that PODXL expression is increased and required for full EMT effects (*40, 41*), we saw variable contribution of PODXL to EMT status. PODXL overexpression in low-surface PC3 cells increased epithelial characteristics (E-cadherin levels; Fig. S4F,G), while PODXL depletion in PC3 high-surface PODXL cells also increased E-cadherin levels (Fig. S5L,M). In contrast, in MDA-MB-231 cells, PODXL depletion increased epithelial, and decreased mesenchymal, markers (Fig. S5N,O) This reveals a complex contribution of PODXL the EMT status of cells, which differs in different contexts. In PC3 cells, invasion is related to increased cortical levels of PODXL, not only the EMT status of cells (Fig. 2I).

### Ubiquitination controls cortical levels of PODXL

As cell populations sorted for different PODXL surface levels possessed similar PODXL mRNA and total protein levels, we focused on ubiquitination as a potential mechanism to control PODXL abundance at the cell cortex (*42*). Endogenous PODXL ubiquitination was significantly altered between cells with different surface levels of PODXL; cells with low-surface PODXL levels showing strong ubiquitination of PODXL (Fig. 3A,B). Publicly available datasets indicated ubiquitination occurs on all four cytoplasmic lysine residues of PODXL (K431/525/526/547; phosphosite.org). Mutation of these four residues in GFP-PODXL (K431/525/526/547R; 4K>R, Fig. S6A) resulted in a strong attenuation of ubiquitination (Fig. 3C,D). Ubiquitination-deficient PODXL (GFP-PODXL 4K>R) cortical levels were enhanced (Fig. 3E,F; Fig. S6B,C) and invasive capacity increased (Fig. 3G,H) compared to wild-type PODXL (GFP-PODXL WT). This reveals that low levels or a lack of PODXL ubiquitination are associated with an EMT, which facilitates high cortical expression of PODXL. Notably, ubiquitination-deficient PODXL (GFP-PODXL 4K>R) was able to drive invasion in the absence of an EMT (Fig. S6D-G), suggesting that the ubiquitination status of PODXL is a major target of EMT processes, and PODXL is an amplifier of mesenchymal invasion, not a passenger molecule.

**Figure 3.**
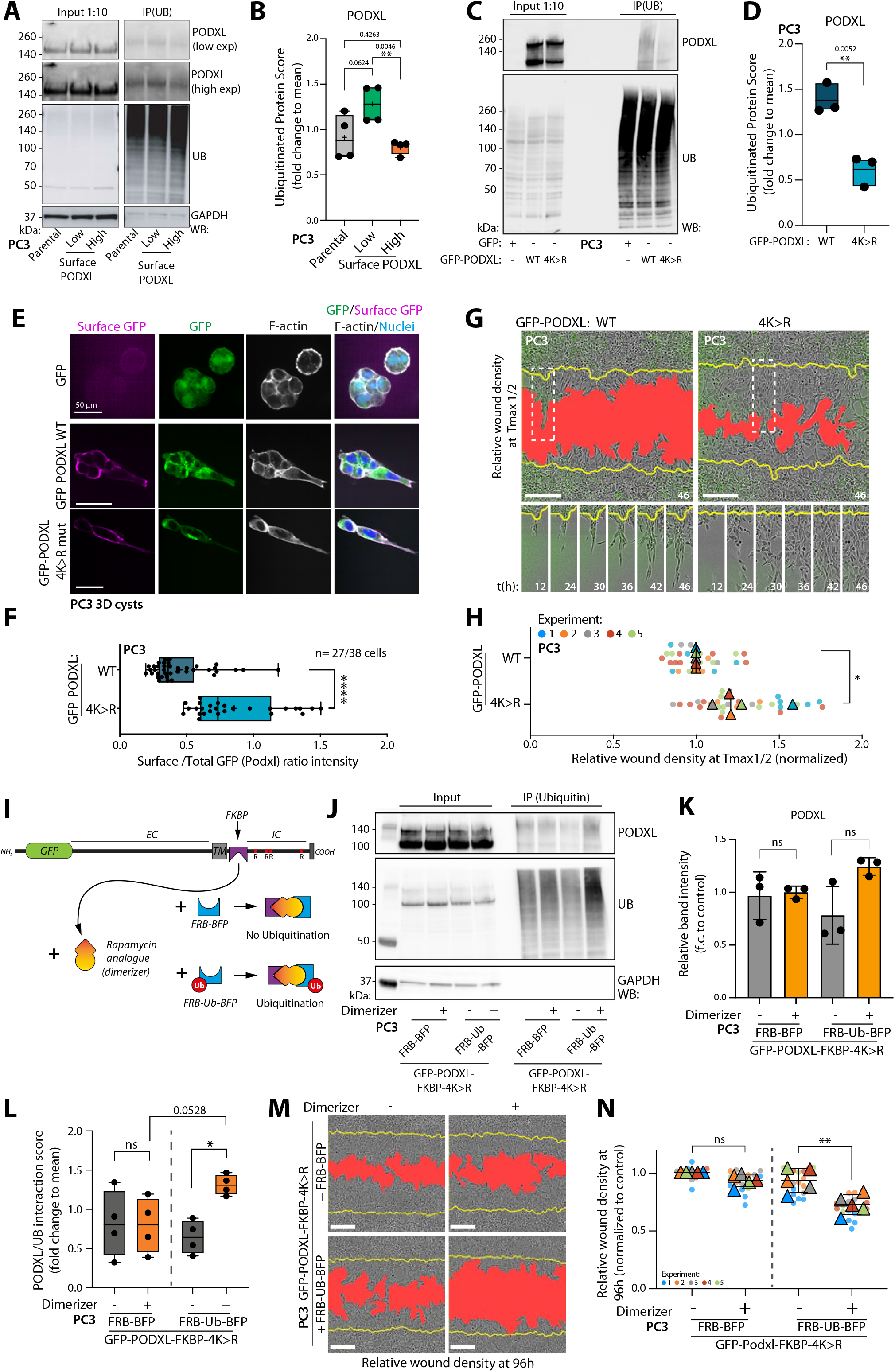
PODXL ubiquitination controls cortical levels. (A-D) Western blot and quantitation, in PC3 and (A-B) subpopulations sorted for PODXL surface levels or (C-D) expressing GFP or GFP-PODXL WT or 4K>R mutant for PODXL, Ubiquitin and GAPDH (loading control). Box-and-whiskers: dots, replicates; +, mean; midline, median; boundaries, quartiles; B, n=4 independent lysate preparations and protein isolations; D, n=3 independent experiments. (E) Immunofluorescence in PC3 cysts expressing GFP, GFP-PODXL WT or GFP-PODXL 4K>R using anti-GFP antibody. Surface GFP (magenta), total GFP (green), F-actin (grey) and nuclei (blue). (F) Quantitation of Surface:Total GFP (PODXL) intensity ratio from (E). Box-and-whiskers: dots, replicates; +, mean; midline, median; boundaries, quartiles; n=1, 2 technical replicates, 27-38 cysts analysed/condition. (G-H) Phase images and quantitation. Yellow line, initial wound; red, wound at T_max_^1/2^ invasion. Circles, technical replicates (n=4-8/experiment); triangles, average/independent experiment (n=5); colour, experiment number. Mean ± SD. (I) Cartoon, inducible ubiquitination of PODXL. (J-L) Western blotting and quantitation, GFP-PODXL (FKBP+4K>R) with or without dimerization and co-expression of FRB-BFP or FRB-Ubiquitin-BFP for PODXL, Ubiquitin and GAPDH (loading control). Quantitation of total levels of PODXL, normalized to average, n=4 independent experiments. (M-N) 3D invasion of (J), phase images and quantitation as described in (G). Circles, technical replicates (n=2-5/experiment); triangles, average/independent experiment (n=5); colour, experiment number. Mean ± SD. Scale bars, (E) 50 or (G,M) 300 μm. p-values, unpaired Student’s 2-tailed t-test, ns, not significant, *P ≤ 0.05, **P ≤ 0.005, ***P ≤ 0.0005, ****P ≤ 0.0001.

To determine, conversely, if ubiquitination drives low surface levels of PODXL we created an experimentally inducible PODXL ubiquitination system based on rapalogue-induced FKBP-FRB oligomerization (Fig. 3I) (*43*). We inserted an FKBP dimerization domain into the cytoplasmic domain of non-ubiquitinatable 4K>R mutant PODXL immediately after the transmembrane domain followed by the cytoplasmic tail. We co-expressed a TagBFP-tagged FRB dimerization domain fused to ubiquitin (FRB-Ub-BFP) or a dimerization domain lacking ubiquitin as a control (FRB-BFP). Only the combination of FRB-Ub-BFP and dimeriser robustly induced PODXL ubiquitination (Fig. 3J-L), decreased surface level of PODXL (Fig. S6H-I), and reduced the invasive ability of the cell population (Fig. 3M,N). These manipulations altered PODXL localization without change to total GFP-PODXL expression (Fig. 3J,K). These data indicate that differential ubiquitination of PODXL is the key context provided by an EMT controlling whether PODXL participates in invasive activities.

### GAL3 interacts with the extracellular domain of N-glycosylated PODXL

That ECM-adjacent location of PODXL was essential for its function prompted us to investigate cortical PODXL interactors. We used MS-based proteomics to compare the interactome of PC3 cells overexpressing either GFP, GFP-PODXL, or GFP-PODXL sorted for high surface levels, isolated by GFP Trap precipitation (Fig. 4A). We used stable isotope labelling with amino acids in cell culture (SILAC) for accurate protein quantification. We cross-referenced this with the total proteome of GFP-PODXL high surface cells and with RNA-seq comparing PC3 subpopulations sorted for endogenous PODXL at high-surface versus low-surface levels (Fig. 4A).

**Figure 4.**
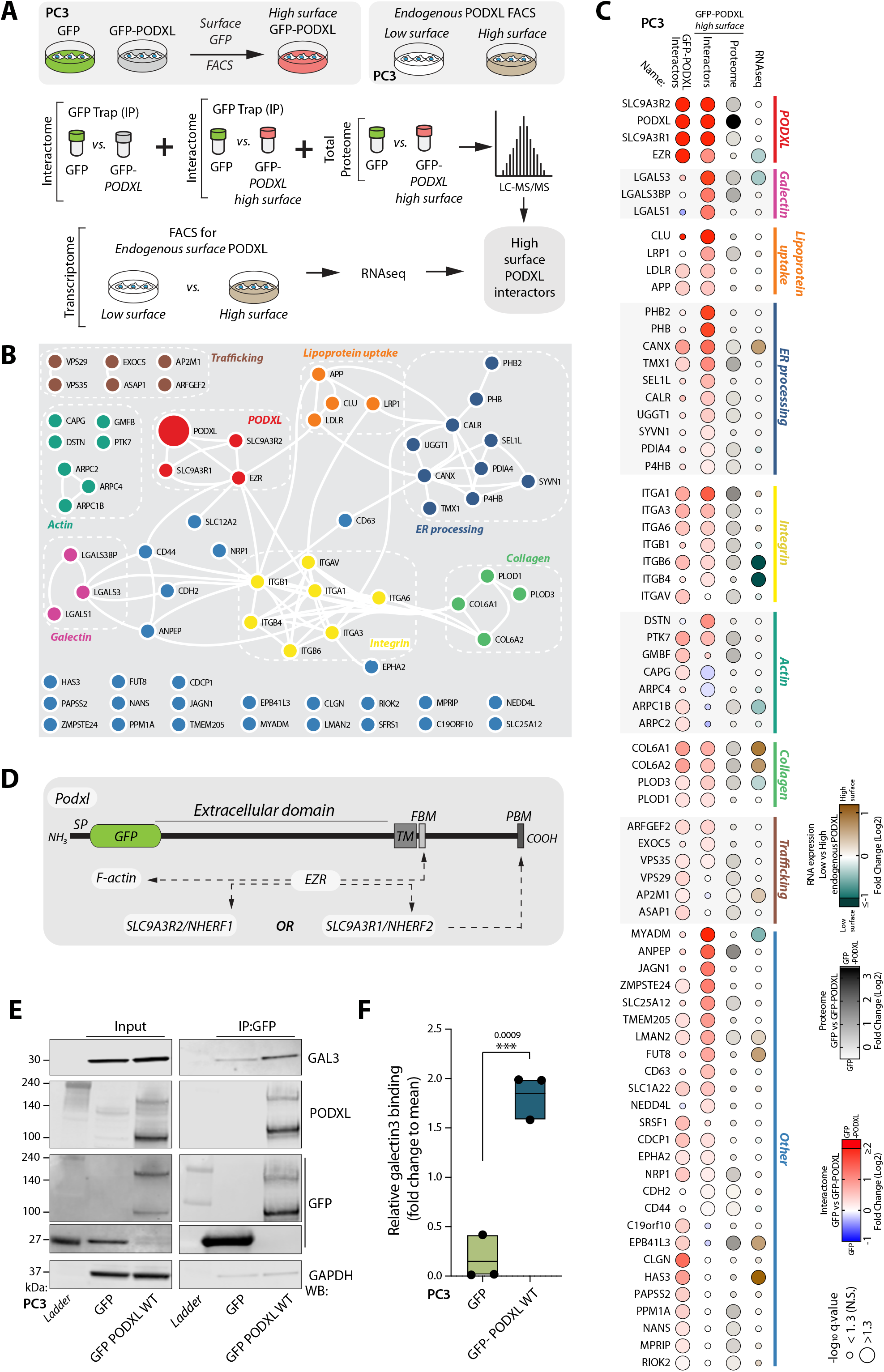
Multi-omic interrogation reveals GAL3 as a cortical PODXL interactor. (A) Schema, multi-omic (interactome, proteome, transcriptome) profiling to identify high-surface PODXL interactors. (B) Network diagram of subgroup interactions among identified PODXL binding partners. Networks, STRING analysis; layout, Cytoscape analysis. (C) Bubble-plot, PODXL binding partners sorted by subgroup then interaction score with high-surface PODXL. For proteomics, colour represents fold change (Log_2_) interaction with PODXL compared to GFP alone; Interactome, blue-to-red; Proteome, greyscale. For RNA-seq, colour represents fold change (Log_2_) endogenous high-surface compared to low-surface PODXL expression. Bubble size, significance (-log_10_). (D) Cartoon, GFP-PODXL domains and known PODXL interactors. SP, signal peptide; TM, transmembrane domain; FBM, FERM-binding-motif; PBM, PDZ-binding-motif. (E) GFP-Trap immunoprecipitation of GFP or GFP-PODXL WT from PC3 cells. Western blotting of input and immunoprecipitates for GAL3, PODXL, GFP and GAPDH (loading control for GFP). (F) Quantitation of relative GAL3 to precipitated GFP vs GFP-PODXL WT in PC3 cells. Floating bar graph: dots, experiments; midline, mean; n=3 independent experimental culturing, lysate preparation and protein isolation. p-values (unpaired Student’s 2-tailed t-test), ns, not significant,*P ≤ 0.05, **P ≤ 0.005, ***P ≤ 0.0005, ****P ≤ 0.0001.

We identified multiple interactors of PODXL, many of which could be organised into functional complexes using STRING network analysis (Fig. 4B,C). In addition to the reported core PODXL complex (PODXL, SLC9A3R1, SLC9A3R1, EZR) (*7, 44-46*) (Fig. 4D), this included networks for glycocalyx-associated components, including Galectins, Integrins, Collagen, but also Lipoprotein uptake, ER processing, Actin regulation, Trafficking and additional proteins not easily assigned to a complex (Fig. 4B,C). The core PODXL complex was the strongest interaction group, but notably these interactions were unchanged between GFP-PODXL versus GFP-PODXL sorted for high expression (Fig. 4C, top clustered group). This excludes changes in the core complex underpinning differential cortical localisation.

We noted robust induction of association of glycocalyx-associated Galectins with high-surface GFP-PODXL, while these were low in GFP-PODXL alone (Fig. 4B,C). LGALS1/GAL1 and LGALS3/GAL3 are beta-galactoside-binding lectins and components of the tumour-associated bulky glycocalyx that regulate function and structure of glycoproteins (*47*). GAL1 has been indicated to promote pro-metastatic function of the glycocalyx (*3*). Whereas GAL3 has been reported as promoter of several kinds of cancer, data from prostate cancer is controversial(*48*); GAL3 is higher in benign prostatic hyperplasia and lower in adenocarcinoma compared to control patients (*48, 49*). Of this Galectin interaction group, GAL3 was the strongest interactor of high-surface GFP-PODXL (Fig. 4C). We confirmed the association of PODXL with GAL3 in independent immunoprecipitations (Fig. 4E,F, Fig. S7A-J), including that non-ubiquitinatable PODXL (GFP-PODXL 4K>R), which has higher surface presentation, showed increased association with GAL3 compared to WT PODXL (Fig. S7D,E).

GAL3 exists in both extracellular and intracellular pools. We mapped the interaction site of GAL3 on PODXL (Fig. S7A-G). GAL3 association with PODXL was maintained with both isoforms of PODXL (PODXLa, PODXLb), upon mutation of binding sites for Ezrin (Ferm-binding motif; FBM*) or NHERF1/2 (DPDZ motif) either alone or together, or upon fusion of Ezrin in place of the PDZ motif (DPDZ + Ezr). This suggests that GAL3 association is independent to core complex association. Indeed, deletion of the entire PODXL cytoplasmic domain (GFP-PODXL DIC) still resulted in robust association of GAL3 with PODXL (Fig. S7F,G), revealing that the association is extracellular. We were unable to recover stable expressers of PODXL with the extracellular domain deleted (GFP-PODXL DEC). In contrast to the lack of effect of the mutants above, mutation of N-linked glycosylation sites in PODXL (5N>Q mutant; (*50*)) abolished GAL3 interaction (Fig. S7A-C). Reciprocally, mutation of the GAL3 carbohydrate recognition binding domain (CRD) to uncouple association of N-acetyllactosamine carbohydrate glycoprotein motifs (R186S; Fig. S5H) (*51*) abolished the association of GAL3 with PODXL (Fig. S7I,J). Although we cannot rule out indirect association due to potential effects of these mutations on other interaction partners, these data suggest that GAL3 associates with the N-glycosylated PODXL extracellular domain via the GAL3 CRD.

### GAL3 controls invasion by regulating surface levels of PODXL

PODXL manipulations that increased surface levels (i.e. GFP-PODXL sorted for high levels, 4K>R) also increased tunnel-forming activity and invasion (Fig. 3G,H; Fig. S7K-M), as well as increased GAL3 association (Fig. 4C; Fig. S7D-G). This initially suggested that GAL3 might be required for cortical function of PODXL. However, this contrasted with *LGALS3* (GAL3) mRNA expression being highest in the non-invasive PC3-low subpopulation and lowered in the invasive high-surface cells (Fig. S7N). Extracellular levels of GAL3 protein were also robustly increased in low-surface PODXL cells (Fig. S7O, P). This suggests that extracellular GAL3 might repress PODXL function. We investigated whether GAL3 was a positive or negative invasion regulator.

GAL3 depletion induced extreme tunnel-forming activity (Fig. 5A-D). Addition of recombinant extracellular GAL3 (reGAL3) to the ECM strongly suppressed tunnel formation in high-surface PODXL cells (Fig. 5E,F). As expected from the heterogenous invasion context of parental cells, overexpression of GFP-PODXL alone failed to induce tunnel formation across most 3D acini (Fig. 5G-J). However, combining this with GAL3 depletion resulted in most 3D structures becoming tunnel-forming. Restoration of GAL3 WT expression (RNAi-resistant TagRFP-T-GAL3), but not non-PODXL-binding GAL3 mutant (R186S), repressed tunnel-forming activity at late timepoints (Fig. 5G-J; note lack of significance in tunnel formation from 60-96h). This was related to PODXL surface levels as GAL3 depletion increased surface PODXL levels in parental PC3 cells and low-surface PODXL cells, but not high-surface PODXL cells that already expressed lowered levels of GAL3 (Fig. 5K,L, Fig. S7N, Fig. S8A,B). Similarly, stimulation with reGAL3 was sufficient to decrease surface levels of PODXL both in parental and high-surface PODXL expressing cells (Fig. S8C-E). This suggests that GAL3-PODXL is a mutually antagonistic ligand-receptor pair, wherein GAL3 triggers internalisation of PODXL from the surface, but in doing so PODXL reduces extracellular GAL3 levels. Accordingly, depletion of PODXL resulted in an increase specifically of the extracellular pool of endogenous GAL3 (Fig. S8F,G). In cells depleted for endogenous GAL3, addition of reGAL3 resulted in its rapid internalisation, but not when PODXL was co-depleted (Fig. S8H-J). Therefore, GAL3-PODXL is a ligand-receptor pair that triggers co-internalisation of both molecules.

**Figure 5.**
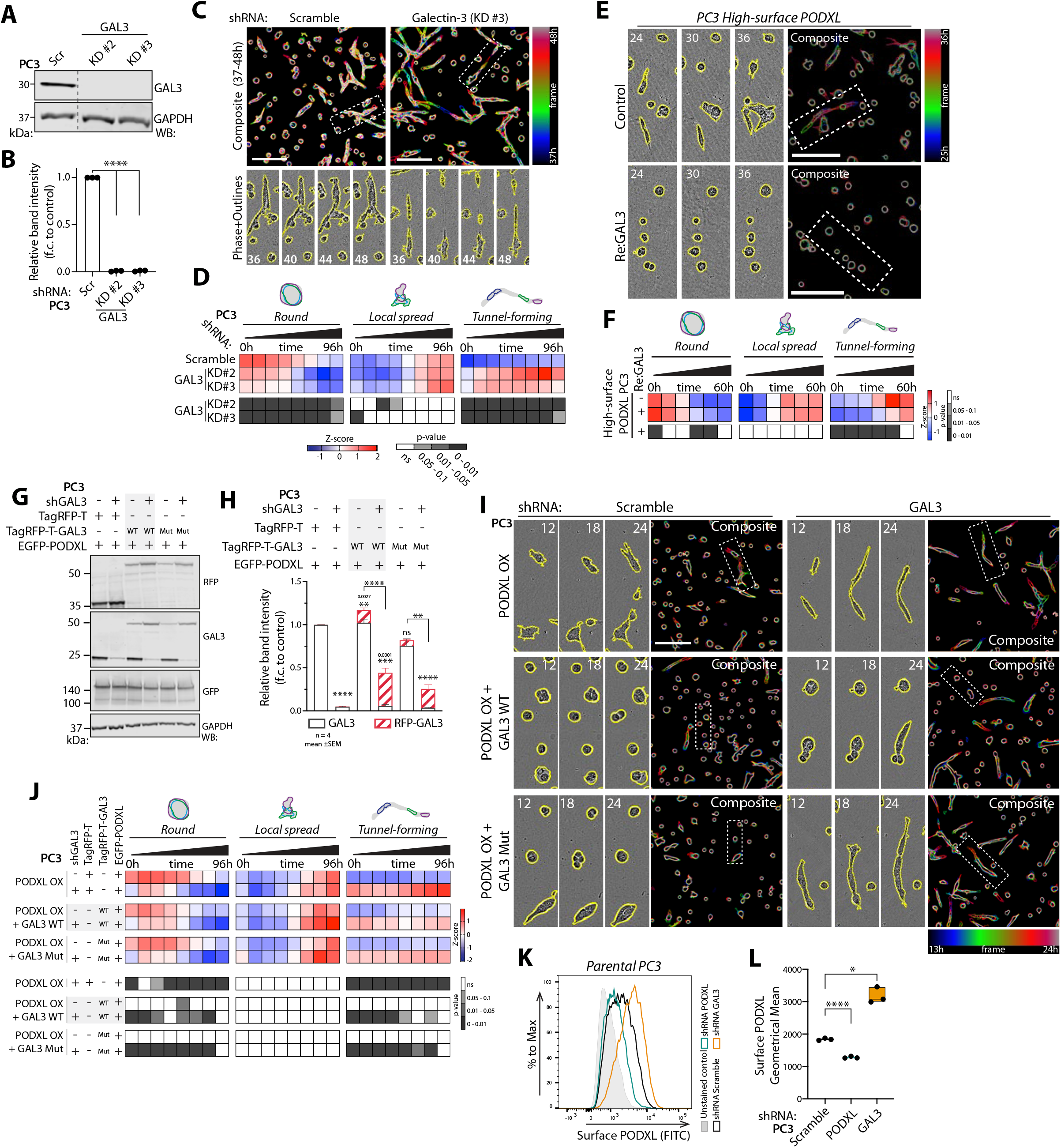
GAL3-PODXL are a mutually antagonistic ligand-receptor pair. (A-B) Western blot and quantitation, PC3-high expressing Scramble or *GAL3* shRNAs for GAL3 or GAPDH (loading control). Lines, not adjacent in gel. Normalised to Scramble, mean ± SD, n=3 independent experiments. (C-D) Phase images and quantitation of (A). Cysts, yellow outlines. Outlines coloured by time in single frames and overlayed. Heatmap, proportions of Round, Spread, Tunnel-forming, Z-score normalised. n=3 independent experiments, 4 technical replicates/condition, 1063-1789 cysts/condition/experiment. (E-F) Phase images and quantitation of PC3-high cysts (control) with recombinant GAL3 (Re:GAL3). Cysts, yellow outlines. Outlines coloured by time in single frames and overlayed. Heatmap, proportions of Round, Spread, Tunnel-forming, Z-score normalised. n=3 independent experiments, 3 technical replicates/condition, 297-577 cysts/condition/experiment. (G-H) Western blot and quantitation, PC3 overexpressing GFP-PODXL WT, co-expressing either TagRFP-T, TagRFP-T-GAL3 (WT or R186S mutant), and Scramble or *GAL3* shRNA for RFP, GAL3, GFP or GAPDH (loading control for RFP). Combined band intensity of GAL3 relative to control. Mean ± SD, n=4 independent experiments. (I-J) Phase images and quantitation of cysts from (G). Cysts, yellow outlines. Outlines coloured by time in single frames and overlayed. Heatmap, proportions of Round, Spread, Tunnel-forming, Z-score normalised. n=2 independent experiments, 3 technical replicates/condition, 287-551 cysts/condition/experiment. (K-L) Representative plot and geometrical mean of surface PODXL levels in Scramble, *PODXL* or *GAL3* shRNA cells. Bar chart: dots, experiments; midline, mean; boundaries, min and max values; n=3 independent experiments. Heatmap p-values, Cochran-Mantel-Haenszel test, Bonferroni adjusted, comparing proportion to control, greyscale as indicated. Scale bars, 300 μm. p-values, unpaired Student’s 2-tailed t-test; ns, not significant, *P ≤ 0.05, **P ≤ 0.005, ***P ≤ 0.0005, ****P ≤ 0.0001.

### GAL3-PODXL binding prevents GAL3 repression of integrin-dependent invasion

Experimental manipulations that removed the ability of PODXL to be ubiquitinated (4K>R; Fig. 3D) enhanced association of GAL3 with PODXL (Fig. S7D-G), yet these cells were uncoupled from the inhibitory effect of GAL3 on invasion (Fig. 3G,H; S7K-M). Conversely, expression of a non-GAL3-binding PODXL mutant (N-linked glycosylation-deficient; 5N>Q) uncoupled GAL3 association (Fig. S7A-C) but was unable to restore invasion and tunnel-forming activity to PODXL-depleted cells, as could be seen when knockdown cells were rescued with WT PODXL (Fig. S7Q). This contrasts with depletion of GAL3 strongly inducing tunnel-forming activity (Fig. 5A-D). This suggests that GAL3 is inhibitory to invasion, and this inhibition is relieved by binding to PODXL. Therefore, PODXL may be a decoy receptor to relieve a repressive extracellular function of GAL3; uncoupling GAL3 from PODXL does not relieve the repression GAL3 confers on other invasion-regulating pathways.

Our PODXL-interactome analysis revealed association of PODXL with cell-ECM interactors (Integrins, Collagen) (Fig. 4B-C), in particular the enhanced association of b1-integrin with high-surface level PODXL. A previous report indicated that PODXL repressed surface levels of ⍰1-integrin in ovarian cancer cells (*23*). GAL3 associates with 1-integrin (*52*) and controls its surface distribution and clathrin-independent 1-integrin internalization (*53, 54*). We speculated that GAL3-PODXL co-internalisation might act to relieve GAL3-dependent integrin internalization, therefore promoting integrin function in invasion.

We used flow cytometry to measure steady state surface levels of total and active integrins. Despite equivalent total ⍰1-integrin levels between parental and sublines (Fig. S9A), low-surface PODXL cells display a strong reduction in the ratio of active:total ⍰1-integrin steady state surface levels, whereas high-surface PODXL cells have robust levels of cortical active:total ⍰1-integrin ratio (Fig. S9B-F). Accordingly, overexpression of GFP-PODXL WT or 4KR mutant increased the active: total surface ratio of ⍰1-integrin (Fig. S9G-L). PODXL depletion decreased the surface levels of active ⍰1-integrin (p=0.0845; Fig. S9M-Q). In contrast, GAL3 depletion did not affect the active:total surface ⍰1-integrin ratio, but rather increased the absolute levels of ⍰1-integrin on the cell surface and consequently the absolute levels of active surface ⍰1-integrin (Fig. S9M-P). This reveals that surface PODXL is associated with ⍰1-integrin activation on the cell surface, while GAL3 controls total ⍰1-integrin cell surface levels.

We used a quantitative biochemical approach (capture-ELISA) to directly measure whether altered surface levels of integrin may be related to its internalization (*55*). Whereas GFP-PODXL WT internalisation was not altered in GAL3-depleted versus control cells at the timepoints measured, GAL3-depleted cells presented a significant decrease in a5⍰1-integrin internalization (Fig. 6A-D). Ubiquitin-deficient PODXL (GFP-PODXL 4K>R), which has increased association with GAL3 (Fig. S7D,E), exacerbated this effect, displaying slowed internalisation of PODXL (Fig. 6E,F) and robust inhibition of a5⍰1-integrin internalization (Fig. 6G,H). This suggests that GAL3-PODXL interaction may control invasion by regulating integrin surface levels and association with the ECM. Accordingly, cells sorted for endogenous high-surface PODXL levels, or expressing GFP-PODXL (WT or 4K>R) showed enhanced association with Matrigel and Collagen-I, but not Fibronectin or Laminin-1 (Fig. S10A-D). Moreover, while cells depleted for PODXL and expressing control GFP alone formed round acini surrounded by Collagen-IV, rescue with RNAi-resistant GFP-PODXL WT or 4K>R expression resulted in cells forming tunnels lined by Collagen-IV (Fig. S10E; Supplementary movie 7). As this Collagen-IV antibody detects mouse Collagen-IV provided by the Matrigel, this reveals that one function of ECM-adjacent PODXL is to promote the organisation of the ECM to form collagen-lined invasion tunnels.

**Figure 6.**
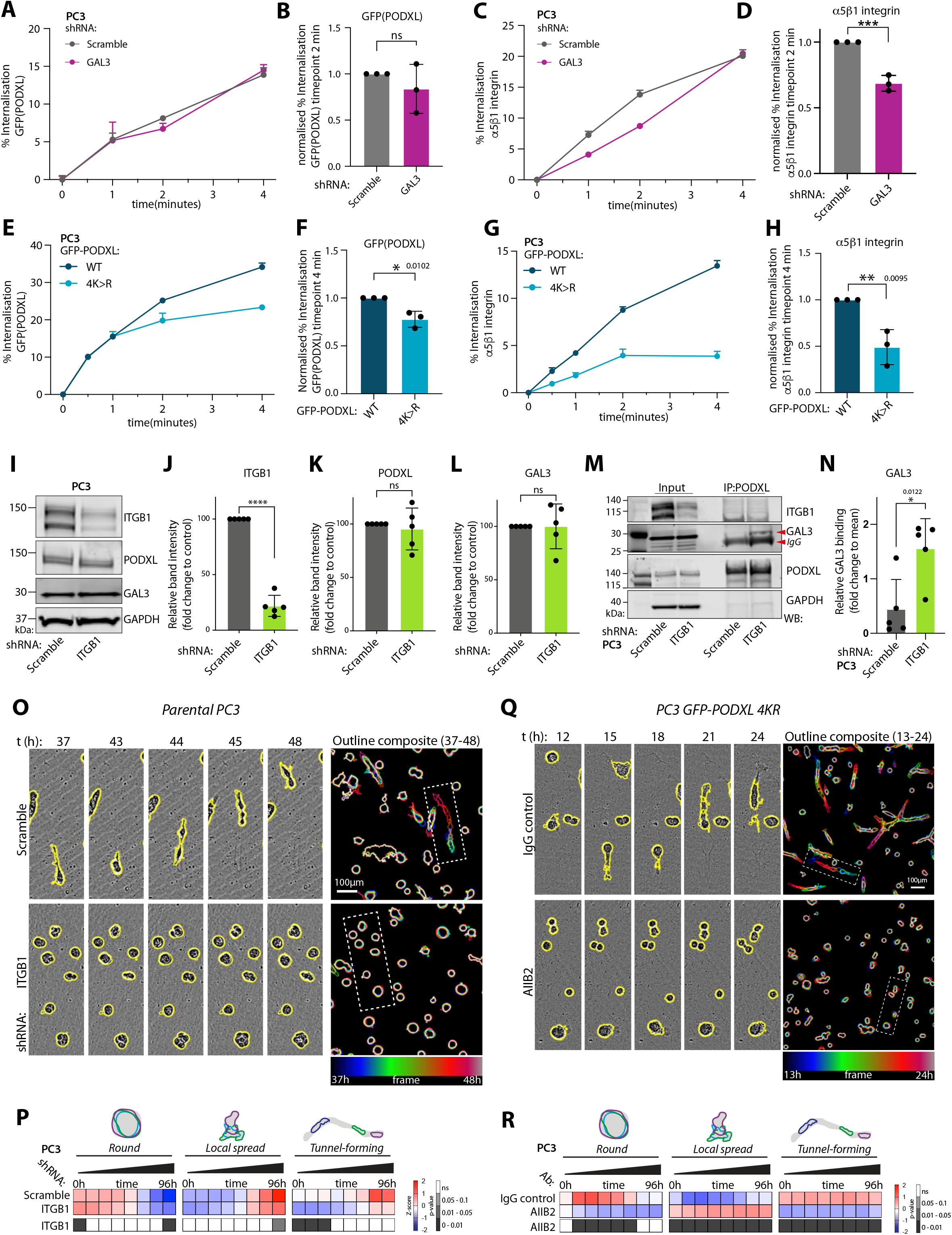
PODXL is a decoy receptor for GAL3 to release GAL3 repression of integrin-dependent invasion. (A-H) Receptor internalisation by capture-ELISA from GFP-PODXL WT-expressing PC3 cells co-expressing Scramble or GAL3 shRNA (A-D) or in cells with GFP-PODXL WT versus 4K>R mutant expression (E-H). Representative graphs, percentage of GFP-PODXL (A), a5 1-integrin (C), GFP-PODXL WT versus 4K>R (E, G) internalisation at indicated timepoints. Values, mean ± SEM, n=3 independent experiments. Graphs showing relative endocytosis at timepoint 2 minutes (B, D) or 4 minutes (F, H) normalised to control; 3 independent experiments combined. (I-L) Western blotting and quantitation, PC3 expressing Scramble or *ITGB1* shRNA for ITGB1, PODXL, GAL3 and GAPDH (loading control). Quantitation across replicates. Mean ± SD, n=5 independent experiments. (M-N) Co-immunoprecipitation, western blotting and quantitation of endogenous PODXL and GAL3 from PC3 in (I) for ITGB1, GAL3, PODXL and GAPDH (loading control). Quantitation across replicates. Mean ± SD, n=5 independent experiments. (O-R) Phase images and quantitation from cells expressing (O, P) Scramble or *GAL3* shRNAs or (Q, R) GFP-PODXL 4K>R mutant (IgG control) with 1-integrin blockade (AIIB2, blocking antibody). Cysts, yellow outlines. Outlines coloured by time in single frames and overlayed. Heatmap, proportions of Round, Spread, Tunnel-forming, Z-score normalised. (P) n=4 independent experiments, 3 technical replicates/condition, 444-702 cysts/condition/experiment. (R) n=3 independent experiments, 2 technical replicates/condition, 137-354 cysts/condition/experiment. Bar graphs, p-values, unpaired Student’s 2-tailed t-test; ns, not significant, *P ≤ 0.05, **P ≤ 0.005, ***P ≤ 0.0005, ****P ≤ 0.0001. Heatmap, p-values, Cochran-Mantel-Haenszel test Bonferroni adjusted, to compare area and proportion of each classification to control, greyscale as indicated. Scale bar 100 μm.

We tested whether ⍰1-integrin was a key effector of surface PODXL-driven matrix alteration to form ECM tunnels. ShRNA-mediated stable depletion of ⍰1-integrin (ITGB1) did not affect expression of PODXL or GAL3 (Fig. 6I-L) but rather enhancedd the GAL3-PODXL interaction (Fig. 6M,N). Depletion of endogenous ⍰1-integrin or inhibition with a function-blocking antibody (AIIB2) abolished tunnel-forming ability in acini (Fig. 6O,P; Fig. S10F,G). Crucially, the high-level tunnel-forming activity of non-ubiquitinatable PODXL (GFP-PODXL 4K>R) was completely abolished by treatment with ⍰1-integrin function-blocking antibody (AIIB2; Fig. 6Q,R). Collectively, these data demonstrate that i) GAL3-PODXL are a ligand-receptor pair whose, ii) co-internalisation acts as a decoy for GAL3, thereby iii) relieving GAL3-directed integrin internalisation and iv) resulting in integrin-dependent remodelling of Collagen in the ECM to form invasive tunnels.

### PODXL cortical localization promotes metastasis *in vivo*

To assess the *in vivo* function of PODXL in tumorigenesis, we performed orthotopic intraprostatic xenografts of PC3 cells manipulated for PODXL expression or localisation into CD1-nude male mice (Fig. S11A). At 8-week endpoint this allows for assessment of primary tumour (PT) formation and multi-organ metastasis (*38*). PODXL-depleted cells showed no defect in engraftment (PT; Fig 7A). Prior to sacrifice at 8-weeks post-transplantation, ultrasound imaging revealed smaller prostate tumours in the PODXL-depleted condition compared to control shRNA-expressing cells (both, 7 mice per condition; Fig. 7B). Macroscopic observation confirmed tumours in additional mice, resulting in 10 control and 9 PODXL-depleted primary tumours, again confirming smaller tumours via decreased prostate weight in the PODXL-depleted condition (Fig. 7C). In mice with a primary tumour, PODXL-depleted cells showed a 2.4-fold reduction in the number of organs that contained macrometastases (detected in 8 out of 10 mice, control; 3 out of 9 mice, PODXL-depleted; Fig 7D, p value=0.0397 using chi-square analysis on raw values). In the low number of mice that still managed to form metastases from PODXL-depleted cells (n=3, PODXL-depleted), these showed no apparent difference in the number of organs with macrometastases per mouse or the tissue tropism of metastasis (Fig. 7E,F). This suggests a major role *in vivo* of PODXL is in tumour formation and metastasis, rather than tissue tropism.

**Figure 7.**
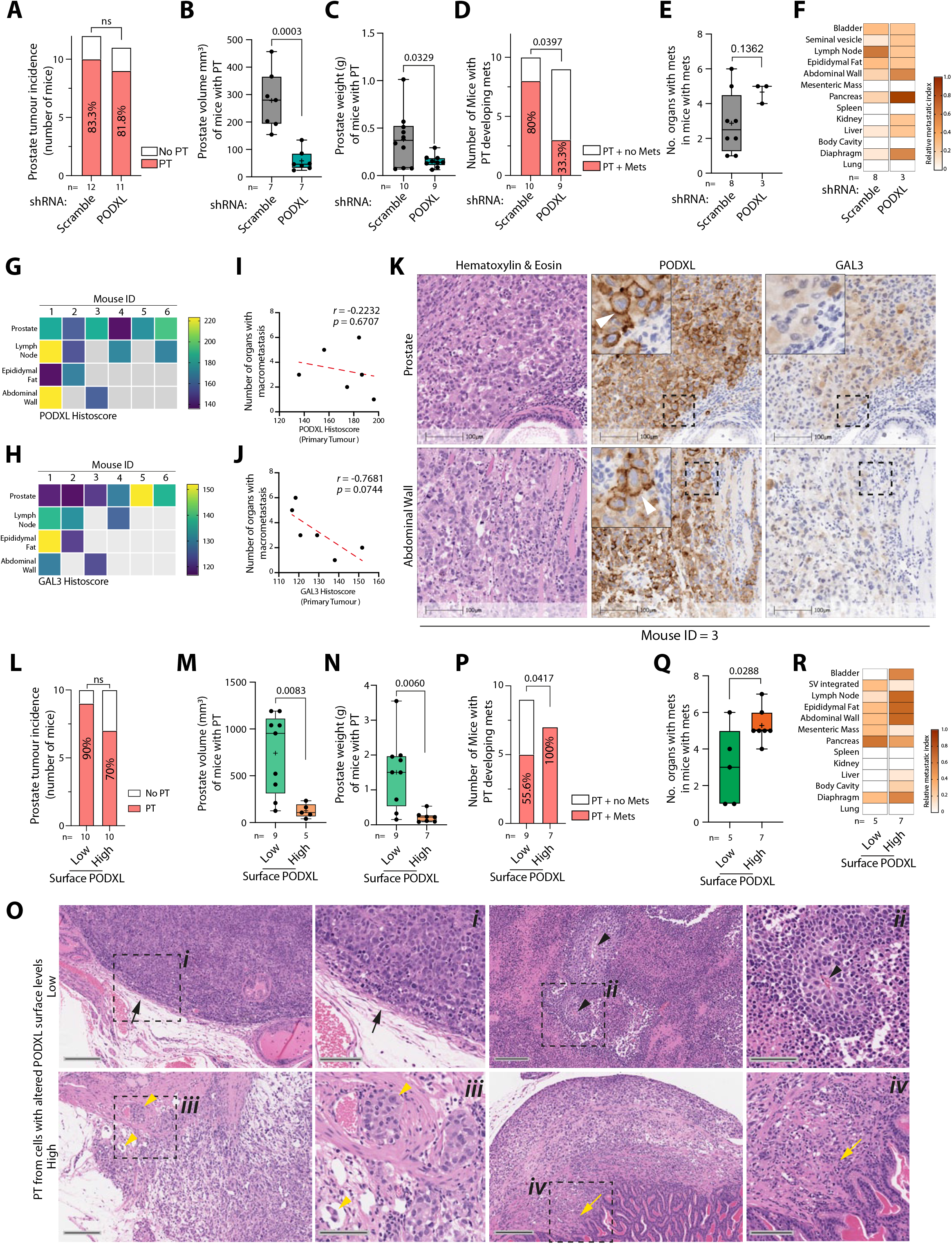
PODXL cortical levels regulates metastasis *in vivo*. (A-R) Primary tumour (PT) and metastasis in mice injected with PC3 (A-K) expressing Scramble or *PODXL* shRNA, n=12 mice/condition or (L-R) sorted for low or high-surface PODXL, n=10 mice/condition. (A, L) PT incidence. In mice with PT, (B, M) prostate volume by ultrasound, (C, N) prostate weight, (D, P) number of mice with metastases, (E, Q) number of metastasis-containing organs/mouse, in mice with metastasis, (F, R) metastasis site tropism. (G-H) Heatmap, weighted histoscore for PODXL or GAL3 across six control mice (Scramble), for PT (Prostate) or metastatic sites. (I-J) Pearson Correlation of Histoscore from PT of six control mice (Scramble) with number of organs in bearing macrometastasis. (K) Serial sections from Mouse ID=3 of the PT (prostate) and metastasis (Abdominal Wall), stained for H&E, PODXL or GAL3. Scale bar, 100μm. Arrowhead, cortical PODXL. (O) Images at intermediate magnification (H&E, scale bar 200μm) with high magnification inserts (H&E, scale bar 100μm). Low-surface PODXL tumours (i-ii), black arrows, solid growth with rounded contours, black arrowheads, geographic necrosis with sparse islands of viable tumour around capillaries. High surface PODXL tumours (iii-iv), yellow arrowheads, looser architecture, infiltrative growth with frequent lymphovascular space invasion, yellow arrow, infiltrative focus. Box-and-whiskers plots, dots, values from individual mice; +, mean; midline, median; boundaries, quartiles; error bars, min to max values; n=3-10 mice that developed PT. p-values (chi-square test on raw counts for contingency tables or unpaired Student’s 2-tailed t-test for box-and-whiskers plots, ns, not significant,*P ≤ 0.05, **P ≤ 0.005, ***P ≤ 0.0005, ****P ≤ 0.0001.

Using a weighted histoscore for intensity that accounts for regional differences within a tumour, both PODXL and GAL3 showed no clear pattern or directionality of change in total labelling between primary prostate tumours and metastases (Fig. 7G,H; n=6 shRNA scramble control mice with primary tumours and metastasis). However, total GAL3, but not PODXL, levels in the prostate tumour showed an inverse association with the number of organs displaying macrometastases (r=-0.7681, p=0.0744, n=6 mice; Fig. 7I,J). Mirroring the heterogeneity in surface levels of PODXL in PC3 cells *in vitro*, control PC3 prostate tumours showed variable PODXL localisation, with regions of clear cortical PODXL labelling (Fig. 7K, arrowheads; Fig. S11B) as well as regions of positive staining lacking cortical PODXL. In serial sections of these tumours, GAL3 labelling also displayed regional differences in labelling, which was not mutually exclusive to regions of high surface PODXL. In contrast, examination of metastases to sites common to all mice with metastases (Lymph Node, Epididymal Fat, Abdominal Wall) revealed prominent cortical PODXL localisation (arrowheads) in the majority of metastases (Fig. 7K, Fig. S11B, arrowheads), and low but not absent levels of GAL3 in the majority of metastases. Collectively, this suggests an association of high-surface localisation of PODXL with metastasis.

To test this association, we performed intraprostatic transplantation of PC3 cells sorted for alternate surface PODXL levels (low-surface vs high-surface) (Fig. S11A). These two conditions similarly showed no difference in engraftment (Fig. 7L). Ultrasound imaging pre-sacrifice showed high-surface PODXL cells presented strongly reduced PT size compared to low-surface PODXL cells (Low surface, n=9; high-surface, n=5; Fig. 7M). Macroscopic observation and weighing prostates after sacrifice allowed detection of additional small PTs in low surface PODXL cells, indicating lower PT weight in high-surface PODXL cell tumours (low-surface, n=9; high-surface, n=7; Fig. 7N). There were differences in tumour morphology between high and low surface PODXL conditions (Fig. 7O). High-surface PODXL tumours displayed a less cohesive growth pattern with some preservation of acinar arrangement. These were much less circumscribed with infiltrative edges and a propensity to involve adjacent tissues, such as adipose tissue or seminal vesicle (4/5 cases) and lymphovascular spaces (3/5 cases). The low surface PODXL group were solid in their growth pattern, though circumscribed with a rounded contour and infiltration of adjacent tissues and lymphovascular spaces was only observed in 1/6 of the cases. Much of the low-surface PODXL tumour mass was necrotic with occasional islands of viable tumour seen adjacent to central capillaries.

Mirroring the infiltrative edges observed in high-surface PODXL primary tumours, this condition presented with a 100% metastasis rate, compared to a ∼55% metastasis rate from low-surface PODXL cells (Fig. 7P; high-surface PODXL, 7/7 of mice; low surface PODXL, 5/9; p value=0.0417 using chi-square analysis). Moreover, high-surface PODXL cells displayed a ∼1.8-fold increase in the number of organs presenting with macrometastasis per mouse with primary tumour (Fig. 7Q), without altered tissue tropism (Fig. 7R). These data confirm an *in vivo* contribution of PODXL to metastasis with high-surface PODXL levels representing a context that drives frequent metastasis.

### PODXL-GAL3 expression stratifies poor clinical outcome

In clinical samples, a high cortex-to-cytoplasm ratio for PODXL localisation is a superior predictor of outcome than PODXL expression alone (*11, 23-28*). Our studies mirror this, with only PODXL at the cell surface promoting invasion *in vitro* and high-level metastasis *in vivo*; we identify GAL3 as a key regulator of surface PODXL levels. We examined whether combining GAL3 expression with PODXL from tumours would allow stratification of poor outcome patients.

*PODXL* mRNA levels were largely unchanged between normal prostate and primary tumour tissue, but were strongly elevated in metastatic samples and associated with disease recurrence across multiple patient cohorts (Fig. 8A,B, Fig. S12A-D). Comparison of patients based on quartiles (Q1-4) of expression revealed that highest *PODXL* expression (*PODXL*^HI^; Q4) identified patients with faster disease progression, with effect sitting below or at significance cut-off depending on cohort (Fig. 8C; Fig. S12E,F). *LGALS3* mRNA displayed the inverted pattern wherein expression was strongly decreased in metastases and associated with disease recurrence across all cohorts, but variably altered between normal tissue and primary tumour (Fig. 8D,E, Fig. S12G-L). Lowest *LGALS3* expression (*LGALS3*^LO^; Q1) identified patients with accelerated disease progression in two of three cohorts (Fig. 8F; Fig. S12M,N).

**Figure 8.**
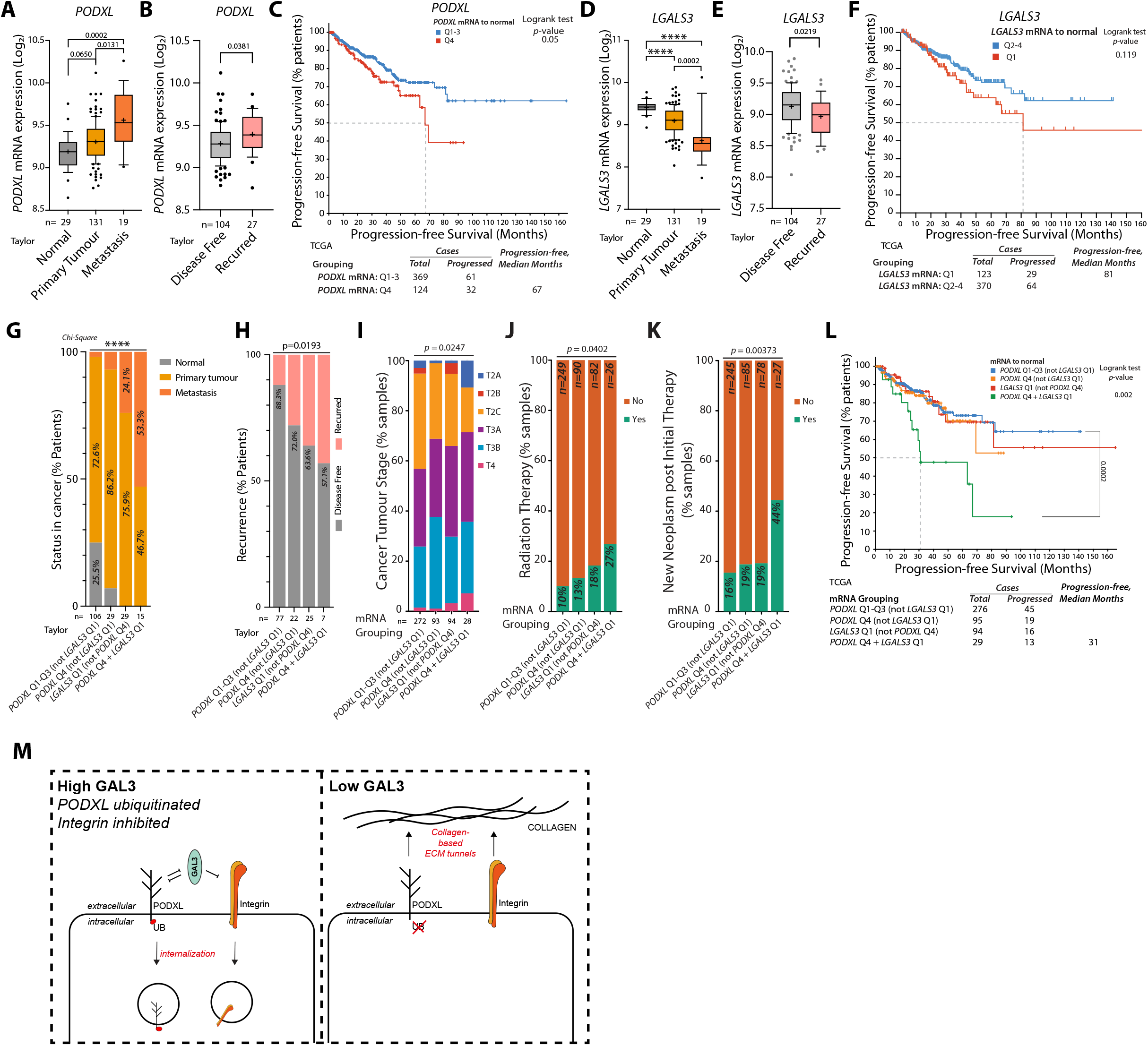
PODXL-GAL3 expression levels identify metastatic, treatment-resistant, poor outcome tumours. (A-B) *PODXL* mRNA expression in (A) normal prostate, primary tumour and metastasis or (B) primary tumours in Disease-free or recurred patients. (C) Progression-free survival (% patients, over months) in patient cohorts of high (quartile 4, Q4) versus not high (quartiles 1-3, Q1-3) *PODXL* mRNA expression. (D-E) *LGALS3* mRNA expression in (D) normal prostate, primary tumour and metastasis or (E) primary tumours in disease-free or recurred patients. (F) Progression-free survival (% patients, over months) in patient cohorts of low (quartile 1, Q1) versus not low (quartiles 2-4, Q2-4) *LGALS3* mRNA expression. (G-L) Clinical parameters in patient cohorts for combinations of high PODXL (Q4) and low *LGALS3* (Q1). (G) Sample type (Normal, Primary Tumour, Metastasis; % patients). (H) Disease-free versus recurred status (% patients). (I) Tumour stage (% samples). (J). Radiation Therapy status (% samples). (K) New neoplasm post initial therapy status (% samples). (L) Progression-free survival (% patients, over months). (M) Model, PODXL-GAL3 interplay controlling invasion. Box-and-whisker graphs (A, B, D, E): 10-90 percentile; +, mean; dots, outliers; midline, median; boundaries, quartiles. Patient numbers presented in figure. Statistical analyses, Kruskal–Wallis Test (A, D), Welch’s t-test (B, E), Log-rank t-test (C, F, L) or chi-square test (G, H, J, K, L). p values: ns, not significant,*P ≤ 0.05, **P ≤ 0.005, ***P ≤ 0.0005, ****P ≤ 0.0001. Data, Taylor (GSE21032) or TCGA (PRAD cohort).

We examined whether grouping patients based on combination of *PODXL*^HI^ and *LGALS3*^LO^ expression would identify prostate cancer patients with different clinical parameters. We divided patients into four groups: a) *PODXL* not high, b) *PODXL*^HI^ but not *LGALS3*^LO^, c) *LGALS3*^LO^ but not *PODXL*^HI^, and d) combined *PODXL*^HI^ and *LGALS3*^LO^(Fig. 8G-K). This resulted in a stepwise decrease in clinical favourability in outcome for patients, with the *PODXL*^HI^-*LGALS3*^LO^ grouping identifying patients with poorest outlook. The *PODXL*^HI^-*LGALS3*^LO^ combination was not found in normal tissue samples but: *i)* was present most frequently in metastases (53% of samples; Fig. 8G), *ii)* was found most frequently in tumours from patients with recurring disease (42.9% patients; compare to 17.7% recurrence when *PODXL* was not high and *LGALS3* was not low; Fig. 8H), *iii)* was associated with higher tumour stage (Fig. 8I), and *iv)* occurred in patients that had received radiation therapy (Fig. 8J). Furthermore, *PODXL*^HI^-*LGALS3*^LO^ patients were 2.3-fold more likely to develop a new neoplasm after initial therapy (44% of patients; compare to 16-19% in other groups; Fig 8K). The combination of *PODXL*^HI^-*LGALS3*^LO^ selected for patients with significantly reduced progression-free survival (Fig. 8L), improving on stratification based on PODXL or LGALS3 expression alone (Fig 8C, F). These data indicate that the collective interplay between PODXL-LGALS3 could be utilised as a potential biomarker of poor outcome, highly metastatic disease in prostate cancer patients.

## DISCUSSION

The bulky tumour-associated glycocalyx can promote tumour growth and metastasis by regulating cell-ECM signalling, yet some of these components are apical membrane-located molecules in normal, apical-basal polarised epithelia, raising the question of how they function at the cell-ECM interface. Using Podocalyxin as a model for the duality of these proteins, our data indicate that an additional context is required to control localisation of these components to the ECM interface.

Although PODXL has been implicated in the aggressiveness of multiple cancer types (*11-17*), the cellular mechanisms by which PODXL confers this function have remained largely unclear. Interaction of PODXL with a number of cytoplasmic proteins, such as Ezrin/EBP50 (*56*), Dynamin-2 (*57*), gelsolin (*58*), cortactin (*59*), and ARHGEF7/Rac1 (*13*) suggested a function of PODXL in regulating cytoskeletal dynamics. Key work from the Roskelley and McNagny labs indicated that in breast cancer cells, PODXL drives an Ezrin-dependent collective migration and tumour budding phenotype (*60*). This Ezrin-dependence was further corroborated in breast cancer cells when studying the role of PODXL in tumour cell extravasation, which curiously reported the PODXL extracellular domain as dispensable for extravasation (*41*). This is in contrast to blocking antibodies to the PODXL extracellular domain showing a decrease in tumour burden and metastasis (*22*). In our system, we find that the entire cytoplasmic domain of PODXL is dispensable for invasive activity. This suggests that there are key differences in how PODXL is contributing to tumourigenesis in different cell types. In prostate cancer cells, at least, we identify that at the surface, the major function of PODXL resides in its extracellular domain, acting as a rheostat on the integrin-repressing activity of the glycocalyx component GAL3.

Our findings allow us to present a model of how this GAL3-PODXL complex controls integrin-based invasion (Fig. 8M). Expression of PODXL alone is insufficient to stimulate invasion and instead requires PODXL retention mechanisms at the ECM-interacting membrane domain. Indeed, even in polarised epithelia where PODXL is normally apically localised, a pool of PODXL is delivered to the basolateral domain before retrieval in a phosphorylation-dependent fashion(*7, 61*). Experimental mimicking of these phosphorylation events in PODXL induces PODXL localisation to the ECM-abutting membrane and drives invasion We here identify that ubiquitination is antagonistic to such retention. In the glycocalyx, GAL3 normally acts to repress integrin-driven invasion by triggering integrin internalisation. Retention of PODXL at ECM-abutting membrane relieves this integrin inhibition where PODXL acts as a decoy receptor for GAL3, controlling extracellular GAL3 uptake. Concomittantly, GAL3 acts to trigger PODXL internalisation. The balance of PODXL-GAL3 modifies the glycocalyx to tune the invasion capacity of cells. A PODXL^HI^-GAL3^LO^ combination therefore allows for high-level integrin-based invasion, which we demonstrate occurs along collagen-lined invasion conduit tunnels. We report that differential ubiquitination of PODXL is a key event during EMT to control invasion.

The identity of the ubiquitin ligase and proteases that regulate PODXL ubiquitin directly remain unclear. The E3 ligase Nedd4-2 has been reported to control PODXL complex stability by regulating the ubiquitination of PODXL complex component Ezrin, but not PODXL itself (*62*). Counterintuitive to this model, we detected Nedd4-2 association only with high-surface PODXL interactome. Similarly, we do not yet know how ubiquitination affects PODXL fate. The Rab35 GTPase has been proposed to regulate tethering of PODXL in intracellular vesicles to the cell surface to promote surface PODXL retention (*63-65*), but whether ubiquitination modulates Rab35 association is unknown. Once internalised, a pool of PODXL can be delivered to exosomes and the correct level of PODXL in exosomes is essential for the ability of these extracellular vesicles to confer invasive behaviours onto recipient cells (*65*). The release of PODXL in exosomes *in vivo* can also prepare the pre-metastatic niche for cell arrival by controlling ECM organisation (*65*). Notably, this PODXL-regulated phenotype transfer also involves modulation of integrin trafficking and consequent remodelling of extracellular matrix, mirroring our findings. Ubiquitination of PODXL may therefore be a mechanism to fine-tune cell autonomous invasive activity versus paracrine signalling to the stroma via exosomes, regulating a permissive ECM organisation for invasion and metastasis. Whether PODXL also plays a role in matrix secretion remains unknown.

A key finding from our studies is that although all cells expressed PODXL, only some of these were able to undergo invasion. We identify that this context depends on the surface delivery of PODXL for its ability to modulate GAL3. The identification of GAL3-PODXL as a mutually repressive ligand-receptor pair, the balance of whose interplay impacts on integrin signalling, was unexpected as GAL1 association with the glycocalyx promotes ECM-integrin signalling in glioblastoma (*3*) and PODXL represses surface levels of 1-integrin in ovarian cancer cells (*23*). Moreover, GAL3 has been reported as a positive regulator of tumourigenesis in other contexts (*66*). Our findings revealed, contrastingly and in prostate cancer at least, that GAL3 is an inhibitor of integrin-dependent invasion with PODXL acting as a decoy to release GAL3-mediated repression of integrins. Why GAL3 may have opposing functionality in different tumour types is not known but may relate to whether PODXL presentation on the surface is required for tumourigenesis in all contexts. Indeed, targeting of PODXL to extracellular vesicles is one mechanism to influence metastasis in a non-cell autonomous fashion (*65*). Unravelling these key differences will be important to understanding how inhibition of PODXL could be used for clinical intervention.

Synder et al (*22*) demonstrated the first proof-of-concept that systemic treatment with PODXL-blocking antibodies is both tolerated and can reduce both primary tumour and metastasis of MDA-MB-231 in murine xenografts. The requirement for surface PODXL presentation for metastasis further supports PODXL an excellent target for therapeutic intervention with such antibodies. The identification of an interplay with GAL3 controlling whether PODXL contributed to invasion is clinically important. This provides a molecular mechanism for why high surface-to-cytoplasm ratio of PODXL in tumours is a superior indicator of disease aggressiveness than PODXL levels alone (*11, 23-28*), and we now add that examination of transcript levels of the GAL3-PODXL axis identifies prostate cancer patients with advanced, recurrent, metastatic disease. Accordingly, restricting PODXL participation in the cell-ECM glycocalyx by promoting either PODXL ubiquitination or treatment with anti-PODXL antibodies (*21, 22, 67, 68*) may present attractive mechanisms to attenuate metastasis.

## Materials and Methods

### Plasmids and cell lines

We used pLKO.1 puromycin lentiviral vector for stable knock-down (KD) or pLENTI-Crispr v2 (*69*) lentiviral vector for stable knock-out (KO) of target proteins. Target sequences are listed in Supplementary Table 2. GFP control and GFP-tagged PODXL constructs (Isoform A WT, ΔPBM, FBM*, ΔPBM+FBM*, ΔPBM+Ezrin, 5N>Q, 4K>R, FKBP-4K>R, ΔIC, ΔEC and Isoform B WT) were generated by subcloning using fragment synthesis (GeneArt, Thermo Fisher Scientific) and/or direct mutagenesis (Q5® Site-Directed Mutagenesis Kit, NEB) and cloned into the retroviral expression vector pQCXIZ by either restriction digestion and ligation, or by In-Fusion cloning (Takara). FRB-BFP control and FRB-Ubiquitin-BFP constructs were generated by fragment synthesis and cloned into the retroviral expression vector pQCXIH. TagRFP-T control and TagRFP-T-LGALS3 constructs (WT and R186S mutant, both *LGALS3* RNAi-resistant) were inserted into a modified version of the lentiviral expression vector pLEX303. In all instances, GFP-PODXL WT and mutants are based on the PODXLa splice variant (NM_001018111.3), except for where indicated that GFP-PODXLb (NM_005397.4) is used.

Stable cell lines were made by stable lenti- or retro-viral transductions. For lentivirus, we co-transfected HEK293-FT cells with plasmids of interest with VSVG (pMD2.G; Addgene plasmid 12259) and psPAX2 (Addgene plasmid 12260) packaging vectors into HEK293-FT cells using Lipofectamine 2000 (Thermo Fisher Scientific). Viral supernatants at day 2 and 3 after transfection were collected, centrifugated at 3,500xg, filtered using PES 0.45 μm syringe filters (Starlab) to remove cell debris, and concentrated using Lenti-X concentrator (Clontech) following the manufacturer’s protocol. Recipient cells were transduced with lentivirus for 72 h before either selection with suitable antibiotic and/or FACS sorting for fluorescent protein using BD Fusion Sorter, where appropriate. For retroviral transfections, 293-GPG packaging cells were transfected with retroviral plasmids were transfected into 293-GPG packaging cells using Lipofectamine 2000, viral supernatants were collected daily at days 5-7 after transfection and separated from cell debris as described above. Recipient cells were incubated with viral supernatants for 24h at 32 ºC with 10 μg/mL Polybrene (Millipore), then the medium was changed and cells were incubated for 48h at 37 ºC, before selection in appropriate antibiotic-containing medium and/or cell sorting for fluorescent proteins. Antibiotic concentrations used were: 5 μg/mL blasticidin (InvivoGen), 200 μg/mL hygromycin (Merck), 2 μg/mL puromycin, 300 μg/mL G418 or 100 μg/mL Zeocin (last three from Thermo Fisher Scientific).

### Cell culture

Cell lines were cultured as follows: PC3 (ATCC) cells in RPMI-1640 (#31870) completed with 10% Fetal Bovine Serum (FBS), 2 mM L-glutamine and 1% penicillin/streptomycin. RWPE-1 cells (ATCC) in Keratinocyte Serum Free Medium (K-SFM) with 50 μg/mL Bovine Pituitary Extract (BPE) and 5 ng/mL Epidermal Growth Factor (EGF). MDA-MB-231 (ATCC) in Dulbecco’s modified Eagle’s medium (DMEM, #21969) supplemented with 10% FBS, 2 mM L-glutamine and 1% penicillin/streptomycin. HEK293-FT (Thermo Fisher Scientific) in DMEM with 10% FBS, 2 mM L-glutamine and 0.1 mM Non-Essential Amino Acids (NEAA). 293-GPG (K. Mostov, UCSF) (*70*) in DMEM with 10% heat-inactivated FBS, 2 mM L-Glutamine, 25 mM HEPES, 1 μg/mL tetracycline (Sigma T-7660), 2 μg/mL puromycin (Sigma P-7255) and 0.3 mg/mL G418. Unless specified annotated, all reagents were from Thermo Fisher Scientific. For SILAC-based analysis of Podocalyxin interactome and total proteome, cells were metabolically labelled using SILAC DMEM (Life Technologies) supplemented with 10% 10 kDa dialyzed FBS (PAA), 1% glutamine, and 1% penicillin/streptomycin. SILAC medium used for the “medium” labeled cells contained ^13^C_6_ l-arginine (84 mg/liter) and D4 l-lysine (175 mg/liter), and for the “heavy” labeled cells contained ^13^C_6_ ^15^N_4_ l-arginine (84 mg/liter), and ^13^C_6_ ^15^N^2^ l-lysine (175 mg/liter) (Cambridge Isotope Laboratories). Experiments were conducted when incorporation rate of labelled amino acids was superior to 95%. All cells were routinely checked for mycoplasma contamination and authenticated using short tandem repeat (STR) profiling.

### 3-Dimensional (3D) culture

3D culture of PC3, RWPE1, and MDA-MB-231 was performed as described in previous studies (*38*). Briefly, 96-well ImageLock plates (Essen Biosciences) or 96-well black/clear bottom plates (Greiner, SLS) were precoated with 10 μL of Matrigel per well (BD Biosciences) for 15 min at 37 ºC. 100 μL of single cell suspensions at low density (1.5 × 10^4 cells/mL) in appropriate medium containing Matrigel at 2% (PC3, RWPE-1) or 4% (MDA-MB-231) were seeded on top of pre-coated wells. For 3D immunoblotting assays, 6-well plates were precoated with 180 μL of Matrigel (BD Biosciences) per well for 20 min at 37 ºC. 1.6 mL of single cell suspensions at low-medium density (12.5 × 10^4 cells/mL) in appropriate medium containing 2% Matrigel were seeded on top of pre-coated wells and harvested after 2-3 days. For treatment experiments drugs, blocking antibodies or the appropriate controls were added from the time of plating, unless otherwise specified. Drugs were added as follows: dimerization experiments, 200 μM A/C heterodimerizer (Takara) or ethanol control; for recombinant Galectin-3 (ReGal3) (R. Jacob, Philipps-Universität Marburg), 1.5 μM ReGal3 was added to cells; for 1-integrin inhibition, 8 μg/mL of AIIB2 blocking antibody (DSHB) was added. Unless otherwise specified, 3-4 replicate wells were included per experiment as technical replicates, and experiments were repeated at least in 3 independent occasions.

### Live 3D cyst imaging and analysis

3D cultures were prepared on 96-well ImageLock plates (Essen Biosciences) as described in ‘3D culture’ section. Plates were then imaged at 10x magnification hourly over four days using an IncuCyte® ZOOM or IncuCyte® S3 (Essen BioScience) with Incucyte ZOOM Live Cell Analysis System Software 2018A or 2021A, respectively. Outlines of imaged objects were detected and generated using Incucyte software, then we applied a custom macro in ImageJ v1.51n to generate composite overlays of objects from 12 consecutive hour time frames, rainbow colour-coded by time. CellProfiler v3.1.9 (*71*) was used to obtain morphological features from resultant objects, and a Fast Gentle Boosting classifier was used to train and identify different behaviours (Round, Spread, Tunnel-forming) in CellProfiler Analyst v2.2.1 (*72*). We then used custom pipelines using KNIME Data Analytics Platform v4.0.2 to combine data from multiple experiments, normalise to controls, calculate the Z-score and perform statistical analysis using the Cochran-Mantel-Haenszel test, and Bonferroni adjusted to compare proportion of each classification to control. Heatmaps of normalised data and p-values were created using GraphPad Prism 9. Emphasized-tracks images were performed with ImageJ v1.51n by using “Remove Outliers, radius=25 threshold=2 function”, “Invert LUT”, “Remove NaNs” and applying shadow funcions.

To analyse speed of invasion of cells within the different classifications (Round, Spread and Tunnel-forming), CellProfiler v3.1.9 was used to identify and mask outlines from objects within each behaviour, then ImageJ v1.51n was used to measure the movement of objects that appeared consistently in at least 12 consecutively outlined images, tracking the nucleus position.

### Immunofluorescence in 2D cell/3D cysts, imaging and analysis

3D cultures were prepared on 96-well black/clear bottom plates (Greiner, SLS) as described in ‘3D cyst culture’ section. For 2D, cells were plated on same plates 48h prior to fixation. In both cases, cells were gently washed in PBS prior to fixation in 4% paraformaldehyde (Thermo Fisher Scientific) for 10 min, then washed twice in PBS. Samples were blocked in PFS (0.7% fish skin gelatin/0.025% saponin/PBS) for 1h at room temperature with gentle shaking, followed by addition of Alexa Fluor primary antibodies diluted 1:100 (unless otherwise specified) in PFS overnight (O.N.) at 4ºC. Three washes in PBS were followed by addition of Alexa Fluor secondary antibodies (1:200), Alexa Fluor Phalloidin (1:200) or Hoechst (1:1,000) (all Thermo Fisher Scientific) diluted in PFS for 1h at room temperature, and 3 washes in PBS. For staining of surface proteins, primary antibody diluted in fresh complete medium was added to live cells for 30 min at 4ºC to label the surface proteins and to prevent internalization. Then, cysts were gently washed in PBS, fixed and blocked as described above before appropriate secondary antibodies or dyes were added for 1h at room temperature, followed by 3 washes in PBS. Plates were imaged using either a Zeiss 880 Laser Scanning Microscope with Airyscan or an Opera Phenix™ high content analysis system (PerkinElmer).

For surface/total PODXL quantification, images were processed using Harmony imaging analysis software (PerkinElmer, Harmony 4.9). Briefly, cells expressing GFP-tagged variants of PODXL were stained with Phalloidin to mark the cortex, Hoechst 34580 to mark the nuclei and surface-GFP to detect intensity of cortical PODXL. A combination of these stainings was used to create masks that allow generation of cell regions (cortical, cytoplasmic, nuclear), followed by calculation of mean intensity in each region. For quantification of endogenous surface PODXL, fluorophore signal intensity of Surface PODXL and F-actin were detected using ImageJ v1.51n.

### Wound healing migration and invasion assay

ImageLock 96-well plates (Essen Biosciences) were used for wound healing assays. For invasion, plates were precoated with 20 μL of 1% Matrigel (Corning) diluted in medium O.N. at 37ºC. Cells were resuspended in complete medium at a density of 7×10^5 cells/mL and 100 μL were added per well. After 4h of incubation at 37 ºC, cells forming a monolayer were wounded using a wound-making tool (Essen Biosciences), washed twice with PBS to remove debris and overlaid with either 100 μL of medium (for migration assays) or 50 μL of 25% Matrigel in complete medium followed by addition of 100 μL of complete media after incubation at 37 ºC for 1h to allow Matrigel polymerization (for invasion assays). Wound closure was imaged every hour for 4 days using IncuCyte® ZOOM or IncuCyte® S3 (Essen BioScience). Analysis of relative wound density (RWD) was performed using Incucyte software. Results were normalized to control using Microsoft Excel 16.53 and presented as RWD at each time point (using R v3.6.2) or, unless otherwise specified, at the time point at which the average RWD of control samples reached 50% of confluence (T_max_^1/2^) using GraphPad Prism 9.

### Cell viability assays

2D and 3D cell viability assays were performed using CellTiter-Glo® 3D Cell Viability Assay (Promega) as per the manufacturer’s instructions. Briefly, cells were resuspended at low density (1.5×10^4 cells/mL) either in complete medium for 2D assays or complete media containing 2% of Matrigel for 3D assays. 100 μL of cell suspension was added per well on black/clear bottom 96-well plates (Greiner) either uncoated or precoated with 10 μL of Matrigel for 2D and 3D assays, respectively. 100 μL of equilibrated CellTiter-Glo reagent was added to each well at days 0, 1, 2, 3 or 5 after plating, mixed and prepared for luminescence detection using a Tecan Safire 2 and SparkControl v2.3 software. Statistical analysis was performed using Microsoft Excel 16.57 and GraphPad Prism 9.

### Flow Cytometry and Fluorescence Activated Cell Sorting

For cortical protein localization, cells were washed twice in iced cold PBS, detached using 2mM EDTA in PBS for 15 minutes and resuspended in PBS + 2% FBS. Cells were counted and 3×10^5 cells per sample were blocked for 30 minutes at 4ºC with primary antibodies (5 μg/million cells, unless otherwise specified; Supplementary Table 3), then washed and stained with a corresponding secondary conjugated antibody (1:200, Alexa Fluor, Thermo Fisher Scientific) for 30 minutes at 4ºC. After washing, samples were processed with a BD FORTESSA Z6102 (BD FACSDIVA software, v8.0.1), then analysed with FlowJo software (version 10.1r5) and GraphPad Prism 9. For sorting of cell lines stably expressing fluorescent-tagged proteins, cells were trypsin detached, resuspended in PBS+ 2% FBS, strained through 40 μm EASTstrainer (Greiner) and processed using a BD FUSION SORTER (BD FACSDIVA software, v8.0.1).

### Immunoblotting

When samples were obtained from cell lysates harvested on plastic (2D), cells at ∼80% confluency were washed with cold PBS. Ice-cold Lysis buffer (50mM Tris-HCl, pH 7.4, 150mM NaCl, 0.5mM MgCl2, 0.2mM EGTA, and 1% Triton X-100) with cOmplete protease inhibitor cocktail and PhosSTOP tablets (Roche) was added for 15 minutes. Cells were scraped and passed through a 27 ^1/2^-gauge needle before centrifugation at 14,000 x g at 4 ºC for 15 min, then supernatant was collected. MicroBCA Protein Assay Reagent kit (Pierce) was used for protein quantification. To obtain cell lysates of 3D samples, a similar procedure was followed with the particularity that after the centrifugation step only the top phase of the supernatant was collected, not collecting the semi-opaque layer between pellet and supernatant, rich in ECM components. To calculate protein concentration, 10 μL of each sample were immunoblotted and probed for GAPDH.

To identify proteins from extracellular media, cells were seeded in 100mm plastic plates and permitted growth until they were in exponential phase (∼80% confluent). Complete media was removed followed by three washes in PBS to remove most traces of FBS, and incubated on 7mL of serum-free media for 24 h. Working at 4 ºC onwards, media was collected and serially centrifuged at 300g, 2000g and 10000g for 10, 10 and 30 min respectively, collecting the supernatant after each step. Supernatant pH was adjusted to a final pH of 5 by adding 10% of trifluoroacetic acid (TFA). 70 μL of Strataclean beads (Agilent technologies) were added (10 μL beads/mL media) followed by an incubation step for 1h in a rotor wheel. Beads bound to supernatant proteins were collected by centrifugation at 872g for 1 min, supernatant was discarded and 35 μL of loading buffer was added to the beads, then boiled at 95ºC for 5 minutes. In all instances, SDS-PAGE protein separation in Bolt™ 4-12% Bis-Tris Plus Gels (Thermo Fisher Scientific) was followed by protein transference to PVDF membranes using an iBlot 2 transfer system (Thermo Fisher Scientific). Membranes were incubated for 45 min in Rockland blocking buffer (Rockland) followed by primary antibody (list in Supplementary Table 3) binding overnight at 4ºC (1:1000 unless stated otherwise). After three washes in TBST (Tris-buffered saline with 0.1% Tween®20 detergent), appropriate secondary antibodies were added for 1h to membranes, then washed again in TBST and imaged using an Odyssey Imaging System (LI-COR Biosciences) or ChemiDoc Imager (BioRad). Band quantification was performed using Image Studio Software 6.0 (LI-COR Biosciences) or Image Lab 6.1 (BioRad), respectively. Graphs were generated on GraphPad Prism 9. GAPDH was used as a loading control for each immunoblot and a representative image for each sample set is shown where appropriate. Uncropped membranes are presented in Supplementary Figure 13.

### Co-Immunoprecipitation (Co-IP), GFP-Trap and Ubiquitin-Trap (UB-Trap)

Cell lysates were obtained as described above, using lysis buffer recommended by manufacturers (For Co-IP and GFP-Trap, 10 mM Tris/Cl pH 7.5, 150 mM NaCl and 0.5% NP-40 with 1x cOmplete protease inhibitor cocktail and PhosSTOP tablets (Roche); for UB-Trap, BlastR™ Lysis Buffer (# BLST01, Cytoskeleton) with 1x Protease inhibitor cocktail (PIC02, Cytoskeleton) and 1x De-Ubiquitination/ SUMOylation inhibitor (N-ethylmaleimide and TPEN, NEM09BB, Cytoskeleton)). To identify relevant PODXL protein-protein interactions by Co-IP, 1 mg of cell lysate was immunoprecipitated with 2 μg of anti-PODXL antibody (3D3, Thermo Fisher Scientific) overnight at 4ºC with rotation. Protein complexes were identified by adding anti-mouse IgG agarose (and mouse IgG agarose as control, #A6531 and #A0919 respectively, both Sigma) for 1 hour at 4ºC in rotation, prior to 3 washes in lysis buffer. Immuno-complexes were unbound from beads by resuspension in loading buffer with sample reducing agent and heated in a heat block for 10 minutes at 100 ºC. Protein separation and transfer to PVDF membranes is as described above. For lysates of cells expressing GFP-tagged proteins, immunoprecipitation was performed using GFP-Trap Kit (Chromotek) as per the manufacturer’s instructions. For ubiquitinated-proteins detection, ubiquitination enrichment immunoprecipitation was performed using Signal-Seeker Ubiquitination Detection kit (Cytoskeleton) as per the manufacturer’s instructions. For FKBP-FRB dimerization system experiments, cells were incubated in media containing 200 μM of A/C Heterodimerizer (Takara) or ethanol control for 4h prior to lysis. In all cases, subsequent immunoblotting was performed as described in immunoblotting method.

### RT-qPCR

For detection of mRNA transcriptional levels of *PODXL* amongst cell lines, RT-qPCR was used. RNA was extracted using the RNeasy kit (Qiagen), followed by reverse transcription by using High-Capacity cDNA Reverse Transcription kit (Thermo Fisher Scientific). RT-qPCR was performed using Taqman Master Mix and Taqman primers: PODXL: Hs01574644_m1; GAPDH: Hs02758991_g1 (Thermo Fisher Scientific) per the manufacturer’s instructions. Expression was normaled to GAPDH levels between cells. Three independent samplings of RNA from a single biological condition was used, with one technical replicate per experimental sampling.

### RNA-sequencing

RNA-sequencing was performed as described in previous studies(*73*). Briefly, RNA was extracted as described in the RT-qPCR section, including a DNase digestion step using RNase-Free DNase Set (Qiagen). Quality control of all RNA samples was performed using a 2200 Tapestation and High-sensitivity RNA screentape (Agilent) to measure the quality of the purified RNA, and only with RIN values >7.9 were processed, with 1ug of total RNA used as initial input. Libraries were then prepared using manufacturers standard procedures (Illumina TruSeq stranded mRNA). Libraries were then sequenced on an Illumina NextSeq 500 instrument, on a High-Output 75 cycle run with paired-end 36bp read length.

For data analysis, quality checks and trimming on the raw RNA-seq data files were done using FastQC version 0.11.8, FastP version 0.20 and FastQ Screen version 0.13.0. RNA-seq paired-end reads were aligned to the GRCm38 version of the human genome and annotation using HiSat2 version 2.1.0. Expression levels were determined and statistically analysed by a combination of HTSeq version 0.9.1, the R environment version 3.6.1, utilizing packages from the Bioconductor data analysis suite and differential gene expression analysis based on the negative binomial distribution using the DESeq2 package version 1.22.2. Pathway Analysis was preformed using MetaCore from Clarivate Analytics (https://portal.genego.com/).

### Mass spectrometry (MS): Sample preparation

For evaluation of SILAC labelling, 20 mg of total lysates from cells grown in SILAC medium were used for in-solution protein digestion. Proteins were reduced using 1 mM dithiothreitol (Sigma-Aldrich) for 1 h, RT and alkylated using 5.5 mM of iodoacetamide (Sigma-Aldrich) for 45 min, RT protected from light. A pre-digestion step was performed with Lys-C (Alpha Laboratories) for 3 h, RT. Then, peptides were diluted 4-fold with ABC buffer (Ammonium bicarbonate 50 mM pH8) and digested with Trypsin MS grade (Promega) overnight, RT. The next day, samples were acidified to pH <4.0 using 1% TFA (Sigma-Aldrich). Peptides were desalted and concentrated using C18 Stage Tips.

For the interactome analyses of PODXL, SILAC mixtures were loaded onto 4-12% SDS polyacrylamide gels for electrophoretic separation of proteins. Gel was stained and each gel lane was divided into 8-10 slices. Proteins were digested into peptides using in-gel digestion method (*74*).

For the total proteome analysis of PODXL, 200 mg of protein (mixture containing 100 mg of protein derived from each SILAC condition) was lysed in SDS buffer (4% SDS, 100 mM dithiothreitol, 100 mM Tris HCl pH 7.6). Proteins were digested into peptides using the filter-aided sample preparation (FASP) method (*75*). Subsequently, 50 mg of peptides was fractionated using strong anion exchange (SAX) chromatography on Stage Tips.

### MS data acquisition

Purified tryptic peptides were injected on an EASY-nLC system coupled online to an LTQ-Orbitrap Pro Elite, through a nanoelectrospray ion source (Thermo Fisher Scientific). An active background ion reduction device (ABIRD, SmartSource Solutions, LLC) was used to decrease air contaminants. Peptides were separated on a 20 cm fused silica emitter (New Objective) packed in-house with reverse-phase Reprosil Pur Basic 1.9 mm (Dr. Maisch GmbH). The emitter was heated to 35° C using a column oven (Sonation). The peptides were eluted with a two-steps gradient at a flow rate of 200 nl/min, with buffer A (0.1% formic acid) and buffer B (80% acetonitrile, 0.1% formic acid). The gradient length for in-gel digestion bands was of 150 min, and for the SAX fractions was of 255 min. MS data were acquired in a positive ion mode using data-dependent acquisition with XCalibur software (Thermo Fisher Scientific). The mass range acquired for the full MS scan was 300 to 1650 m/z, with a resolution of 120,000 at 400 m/z. Only multiply charged ions from two to five charges were selected for fragmentation. For PODXL IP samples, higher-energy collision dissociation fragmentation was triggered for the 10 most intense ions, using a maximum injection time of 150 ms or a target value of 40,000 ions, at a resolution of 15,000. For PODXL proteome samples, collision-induced dissociation fragmentation was triggered for the 10 most intense ions, using a maximum injection time of 25 ms or a target value of 5,000 ions. A dynamic exclusion window of 60 s was used for ions that had already been selected for MS/MS.

### MS data analysis

The MS .raw files were processed with the MaxQuant software version 1.5.0.36. The Andromeda search engine performed the search against the human UniProt database (release 2013, 88847 entries). The multiplicity was set accordingly to the SILAC labels of each experiment, and remainder parameters included: minimal peptide length of 7 amino acids; variable modifications acetyl (protein N-term) and oxidation (M); fixed modification carbamidomethyl (C); specificity for trypsin cleavage; and maximum two missed cleavages allowed. For the search of the parent mass and fragment ions, an initial mass deviation of 4.5 and 20 ppm, respectively, was required. The re-quantify parameter was enabled, and match between runs was used between similar fractions. The false discovery rate at the protein and peptide level were set to 1%. Unique peptides with at least two ratio counts were used. The MaxQuant output table “ProteinGroups.txt” was uploaded in Perseus version 1.5.2.11 for downstream analysis. Datasets were filtered to remove contaminants, reverse peptides that match a decoy database and proteins only identified by site. Proteins with at least one unique peptide were used for analyses. Normalized SILAC ratios were log_2_-transformed and intensities were log_10_-transformed. To organize PODXL-interacting proteins into functional cluster we used Cytoscape v3.9.0 with integrated STRING network analysis (0.7 confidence score).

### Internalization assay by capture ELISA

Protein internalization was measured as described in (*55*). Briefly, cells were surface labelled with 0.13 mg/mL NHS-SS-biotin (Pierce) in PBS at 4 ºC for 30 min. Internalization was re-established by placing cells in complete media in the presence of 0.6 mM of primaquine to disrupt vesicle recycling for the indicated times. Cell were placed on ice and biotin removed from the cell surface by MesNa treatment. Following that, cells were lysed and the quantity of biotinylated receptors determined using capture-ELISA. Antibodies used: α5β1 (VC5, BDPharmingen, 555651), TfnR (CD-71, DBPharmigen, 555534), GFP (Abcam, ab1218).

### Internalization assay by immunofluorescence

Recombinant Galectin-3 (ReGal3) internalization was measured by immunofluorescence. Briefly, 1.5 μM ReGal3 was bound to cell surface of PC3 cells in 2D culture for 15 min at 4ºC. Internalization was re-established by placing cells at 37 ºC for 0, 2, 10 minutes. Cells were washed twice in PBS, then fixed in 4% PFA (Thermo Fisher Scientific) for 10 min, and washed in PBS. ReGal3 was detected by immunofluorescence using a primary antibody anti-Gal3 following the procedure described in the Immunofluorescence in 2D cell/3D cysts and imaging section.

### Cell Attachment

96-well black/clear bottom plates (Greiner) were pre-coated with 20 μL of different extracellular matrices as follows (1:100 Laminin (ref), 1:100 or 10 μg/mL fibronectin (#F1141; Sigma-Aldrich), 1:200 or 10 μg/mL of Collagen-I (354236, Corning) or 50% Matrigel (BD Biosciences) in PBS) and incubated O.N. at 37 ºC. Cells were detached by mild methods (2mM EDTA in PBS for 15 minutes) and counted. 1,500 cells in 100 μL were added per pre-coated well, and incubated at 37 ºC for 30, 60, 120 or 240 min, followed by a wash in pre-warmed PBS, fixation and staining, as previously described in immunofluorescence section.

### Animal studies

Animal experiments were performed in compliance with ethical regulations of UK Home Office Project Licence (P5EE22AEE), approved and sanctioned by welfare and ethical review board of University of Glasgow under the Animal Scientific Procedures Act 1986 and the EU directive 2010. CD-1 nude (CD1-*Foxn1*^*nu*^) male mice (6-weels of age) were obtained from Charles River (UK) and acclimatised for at least 7 days. Mice were kept in a barriered facility at 19-22 ºC and 45-65% humidity in 12 hour light/darkness cycles with access to food and water *ad libitum* and environmental enrichment. A cohort of 12 mice (PC3 shRNA Scramble or PODXL) or 10 mice (PC3-High or PC3-Low) were used per experiment. Surgery (under anaesthesia and with analgesia) was performed to implant 2 × 10^6 PC3 cells within the anterior prostate lobe of each mouse, followed by close monitoring of tumour development (including by palpation) and health conditions, until endpoint of the experiment, 8-weeks post-implantation. Primary tumours were imaged and measured (volume) using VevoLAB ultrasound equipment and VevoLAB 3.1.1 software. Metastases were analysed by gross observation in bladder, seminal vesicle, lymph nodes, epididymal fat, abdominal walls, mesenteric mass, pancreas, spleen, kidney, liver, body cavity, diaphragm and lungs. Statistical analysis was performed using Chi-squared test (with Fisher’s exact test) in GraphPad Prism 9.

### H&E and IHC Staining

All Hematoxylin & Eosin (H&E) and Immunohistochemistry (IHC) staining was performed on 4μm formalin fixed paraffin embedded sections (FFPE) which had previously been maintained at 60°C for 2 hours.

FFPE sections for Galectin-3 (Gal3) (87985, Cell Signaling) IHC staining were dewaxed using an Agilent pre-treatment module. Sections were then heated to 97°C for 20 minutes in a high pH target retrieval solution (TRS) (K8004, Agilent) for heat-induced antigen retrieval (HIER), washed in flex wash buffer (K8007, Agilent) and loaded onto an Agilent autostainer link48. Sections were blocked with peroxidase (S2023, Agilent) for 5 minutes, washed with flex wash buffer and Gal3 applied (1/600) for 35 minutes. Sections were washed with flex wash buffer and rabbit envisions secondary antibody (K4003, Agilent) applied for 30 minutes. After another wash in flex wash buffer Liquid DAB (K3468, Agilent) was added for 10 minutes. The sections were washed in water and counterstained with hematoxylin z (RBA-4201-00A, Cell Path).

FFPE sections for Podxl (ab15038, Abcam) were stained on a Leica Bond Rx autostainer undergoing on-board dewaxing (AR9222, Leica) and epitope retrieval using ER2 solution (AR9640, Leica) for 20 minutes at 95°C. Sections were washed with Leica wash buffer (AR9590, Leica) before peroxidase block was performed using an Intense R kit (DS9263, Leica). FFPE sections were rinsed with wash buffer and Podxl antibody applied for 30 minutes (Abcam, 1/1000). Sections were rinsed with wash buffer and rabbit envision secondary antibody applied for 30 minutes. After a final wash in wash buffer sections were visualised using DAB and counterstained with Haematoxylin from Intense R kit.

H&E staining was performed on a Leica autostainer (ST5020). Sections were dewaxed, graded alcohols applied, and then stained with Haem Z (RBA-4201-00A, CellPath) for 13 mins. Sections were washed in water, differentiated in 1% acid alcohol, washed, and the nuclei blu’d in Scotts tap water substitute. After washing with tap water sections were placed in Putt’s Eosin for 3 minutes. To complete the H&E and IHC staining sections were washed in tap water, dehydrated through graded ethanol, and placed in xylene. The stained sections were coverslipped from xylene using DPX mountant (CellPath, UK). Stained FFPE sections were scanned at x 20 magnification using a Lecia Aperio AT2 slide scanner and analysed using Halo software (Indica Labs).

Following immunohistochemistry, tumour areas were manually drawn and analyzed by area quantification using Halo software. Expressions were scored using Histoscore (= S (1 x % area of weak positive staining) + (2 x % area of moderate positive staining) + (3 x % area of strong positive staining). All statistical analyses and graphs were made using GraphPad Prism 9.

### Analysis of cancer cell lines from CCLE

Data from a panel of breast and prostate cancer cell lines was obtained from the CCLE (Cancer Cell Line Encyclopedia, Broad Institute(*76*)) and analysed using GraphPad Prism 9.

### Analysis of patients

Patient data were obtained from cBioportal.org and CANCERTOOL (*77*) and analysed using cBioportal.org tools and GraphPad Prism 9. Datasets include Glinsky (*78*), Grasso (GSE35988), Lapointe (GSE3933), Taylor (GSE211032), TCGA (obtained from cBioportal.org), Tomlins (GSE6099) and Varambally (GSE3325).

## Acknowledgments

We thank the Core Services and Advanced Technologies at the Cancer Research UK Beatson Institute with thanks to the Biological Services Unit, Histology Services, Nikki R. Paul in Beatson Advanced Imaging Resource and Molecular Technologies.

## Funding

This work was supported by the following grants:

D.M.B. NIH K99CA163535, CRUK (C596/A19481), E.S., AR.-F., L.M., A.H., D.S. CRUK (C596/A17196 and A31287), M.M., Royal Society (NF161630), C.M. and R.S. CRUK (A29801) and R.P. and H.Y.L. CRUK (A22904). S.M., J.A., K.J.C, K.B. CRUK (A29799 and A17196), J.N. CRUK (A17196 and A31287), F.K and S.Z. CRUK (A29800).

## Author contributions

D.B. and A.R.F conceived the project. Under supervision of D.B, wet lab experiments were performed by A.R.F, M.M, J.P, E.S., E.C. K.N. and J.N. Under supervision of D.B. and C.M., E.F. and R.S. performed computational analyses. F.K. and S.L. performed mass spectrometry analysis under supervision of S.Z. *In vivo* murine experiments were performed by R.P., S.M., L.G. and J.A. under supervision of K.B, K.J.C, and H.L. Tissue sectioning and labelling was performed by C.N. Histopathology was assessed by K.R. and J.L.Q. Assistance from L.M. and D.S. was given for developing imaging techniques. A.R.F. composed the figures under supervision of D.B. The manuscript was written by A.R.F., E.S., and D.B. with input from all authors. All authors discussed the results and commented on the manuscript.

## Corresponding author

Correspondence to David M. Bryant.

## Data availability

The raw mass spectrometry files and the MaxQuant search results files have been deposited as partial submission to the ProteomeXchange Consortium via the PRIDE partner repository with the dataset identifier PXD032911. The RNA-seq data has been deposited in NCBI BioProject database with Accession: PRJNA897704. All data needed to evaluate the conclusions in the paper are present in the paper and/or the Supplementary Materials

