## Supplementary Text and Figures for "Spatial regulation of the glycocalyx component Podocalyxin is a switch for pro-metastatic function"

**Figure S1. Identifying the phenotype of PODXL depletion.**

(A) Representative western blot for PODXL and GAPDH as loading control in MDA-MB-231 cells stably expressing control (Scramble) or *PODXL* shRNA.
(B) Quantitation at two major band size regions (150kda, 55+65kDa) of (A) from n=3 independent experiments (lysate preparations). Mean ± SD.
(C) Representative western blot for PODXL and GAPDH as loading control in PC3 cells stably expressing control (Scramble) or *PODXL* sgRNA.
(D) Quantitation at two major band size regions (150kda, 55+65kDa) of (C) from n=3 independent experiments (lysate preparations). Mean ± SD.
(E-M) Wound-healing 3D invasion of PC3 (F,G, J) and MDA-MB-231 cells (H,I); or 2D migration of PC3 cells (K-M) assays examined by hourly time-lapse imaging. (E) Schema. Representative phase-contrast images show invasion over 72h (F) or migration over 30h (K). Yellow line, initial wound position; red line, remaining wound region at $T_{max}^{1/2}$ closure (2D, 30h; 3D, 72h). Arrowheads, invasive tunnels. White square, region of interest. Elapsed time, hours (h). Representative experiment, over time mean ± SD, n=3 independent experiments (G, M). Quantitation across experiments (G, I, M). Circles, technical replicates (n=4-8/experiment); triangles, average per independent experiment (n=3-7); colour, experiment number.
(N-O) Cell viability assays in PC3 cells expressing scramble or *PODXL* shRNA in 2D (N) or 3D (O) cell cultures at indicated times after plating. Mean ± SD, n=3 independent experiments.

p-values calculated using unpaired Student's 2-tailed t-test; ns, not significant,*P ≤ 0.05, **P ≤ 0.005, ***P ≤ 0.0005, ****P ≤ 0.0001. Data in superplots (H, I, M) mean ± SD. Circle, technical replicate. Triangle, mean per experiment. Colour, experiment number. Scale bars, 300 µm.

**Figure S2. PODXL depletion attenuates motility in the ECM.**

(A) Schema, method for classification and quantitation of user-define heterogeneous phenotypes. 1. Hourly time-lapse imaging across multiple days of 3D culture of PC3 cysts in 96-well plates by Incucyte imaging. 2. Cysts are automatically detected and converted to digital objects. 3. Object outlines from hourly imaging are pseudo-coloured in a rainbow lookup table according to time and overlaid onto a single image in iterative 12h blocks. 4. Size and shape features are extracted from resulting concatenated objects. A Fast Gentle Boosting classifier is used to define three phenotypes (Round, Local-spreading, Tunnel-forming. 5. Confusion matrix between user-classification and predicted label is used to ensure high-accuracy training (95-98% accurate). 6. Frequency of each phenotype over time, and statistical analysis across technical replicates and independent experiments, is presented in heatmaps.
(B-E) PODXL loss of function and phenotype in PC3 (B,C; sgRNA knockout) and MDA-MB-231 cells (D,E,G; shRNA knockdown), compared to appropriate scramble control. (A,C) Phase-contrast images from time-lapse series. Object outlines, yellow. Time, hours (h). Outlines (PC3, 12h-36h; MDA-MB-231, 25-36h) colourcoded by time and overlaid into single image (Composite). Inverted image at indicated timepoint to emphasize ECM-tunnel in PC3 or MDA-MB-231 cells (Emphasize track). (C,E) Heatmaps, relative levels of user-defined behaviours in Scramble or *PODXL* KO/KD (Round, Spread, Tunnel-forming), Z-score normalised values. For PC3, n=3 independent

experiments, 2-3 technical replicates per condition, totalling 486-825 initial cysts per condition per experiment. For MDA-MB-231, n= 3 independent experiments, 2-3 technical replicates per condition, totalling 283-539 initial cysts per condition per experiment.
(F-G) Mean invasion speed in µm/h within the tunnel-forming behaviour for (F) PODXL knockout (sgRNA) in PC3 or (G) PODXL knockdown (shRNA) in MDA-MB-231 cells, compared to appropriate scramble control. Mean ± SD. For PC3, n=3 independent experiments, 40-48 cysts tracked per experiment (127-135 total independent cysts analysed). For MDA-MB-231, n=3 independent experiments, 46-63 cysts tracked/experiment (161-188 total independent cysts analysed).

p-values stated (G,H) (unpaired Student's 2-tailed t-test), Data in superplots (E,F), mean ± SD. Circle, technical replicate; triangle, mean per experiment. Colour, experiment number. Data in heatmaps (C,E), user-defined classification frequency as a mean of Z-score-normalised values (blue-to-red). Heatmap p-values, Student's t-test and Cochran-Mantel-Haenszel test, and Bonferroni adjusted, compared to control, greyscale as indicated. (B,D) Scale bar, 50 µm.

**Figure S3. Localisation and contribution to 3D behaviour of GFP-PODXL.**
(A) Cartoon, GFP-PODXL domains. SP, signal peptide; TM, transmembrane domain.
(B) Western blotting of cells stably expressing GFP or GFP-PODXL for PODXL or GAPDH as loading control.
(C) 3D phenotypes of GFP-PODXL- or GFP-expressing cells. Phase-contrast images from time-lapse series. Object outlines, yellow. Time, hours (h). Outlines (25-36h) colourcoded by time and overlaid into single image (Composite). Dashed rectangle, area of interest.
(D) Heatmap, relative levels of user-defined behaviours in GFP-PODXL- or GFP-expressing cells (Round, Spread, Tunnel-forming), Z-score normalised values. n=5 independent experiments, 4 technical replicates per condition, totalling 490-1,188 initial cysts per condition per experiment.
(E) Anti-GFP antibody immunofluorescence in 3D cysts expressing either GFP or GFP-PODXL. Surface GFP (magenta), total GFP (green), F-actin (grey) and nuclei (blue).

Data in heatmap (D), user-defined classification frequency as a mean of Z-score-normalised values (blue-to-red). Heatmap p-values, Student's t-test and Cochran-Mantel-Haenszel test, and Bonferroni adjusted, compared to control, greyscale as indicated. Scale bar, 100 µm.

**Figure S4. High cortical PODXL-driven invasion in an EMT.**
(A-C) Wound-healing 3D invasion in parental PC3 cells or subpopulations sorted for High or Low endogenous surface PODXL levels examined by hourly time-lapse imaging. (A) Representative phase-contrast images, invasion over 48h. Yellow line, initial wound position; red line, remaining wound region at $T_{max}^{1/2}$ closure. Arrowheads, invasive tunnels. White square, region of interest. Elapsed time, hours (h). (B) Representative experiment, over time. (C) Quantitation across experiments. Circles, technical replicates (n=4-8/experiment); triangles, average per independent experiment (n=4); colour, experiment number. Mean ± SD, n=4 independent experiments.

(D-E) Cell viability assays in PC3 cells sorted for Low or High endogenous PODXL surface levels in 2D (A) or 3D (B) cell cultures at indicated times after plating. Mean ± SD, n=3 independent experiments.
(F) Western blotting of PODXL, EMT-related markers (E-CAD, ESRP1/2, N-CAD, VIM, ZEB1) or GAPDH in total cell lysates from PC3 sorted for low surface PODXL and expressing either GFP or GFP-PODXL WT.
(G) Quantification from (F) across replicates. Levels in GFP-PODXL WT versus GFP expressing cells; intensity fold change. Mean ± SE, n=3 independent experimental isolations of cell lysates.
(H) Heatmap, relative levels of user-defined behaviours in low-surface PODXL cells expressing GFP or GFP-PODXL WT (Round, Spread, Tunnel-forming), Z-score normalised values. p-values were calculated by Cochran-Mantel-Haenszel test, and Bonferroni adjusted, to compare proportion of each classification to control, greyscale as indicated. n= 3 independent replicates, 3 technical replicates per condition, totalling 469-587 initial cysts per condition per experiment.

Scale bars, 300 µm (A). p-values (C-E,G) (unpaired Student's 2-tailed t-test), ns, not significant,*P ≤ 0.05, **P ≤ 0.005, ***P ≤ 0.0005, ****P ≤ 0.0001.

**Figure S5. Characterization of alterations between high-surface and low-surface PODXL PC3 cells.**
(A) Bubble plot, mRNA for PODXL-complex components from RNA-seq comparing low- and high-surface PODXL expressing PC3 cells. X-axis, q-value (-log10), grey dashed line, significance (p=0.05); colourscale High-surface vs Low-surface mRNA expression fold change (log2, blue to red); bubble size, normalised count reads (log10), n=4 independent RNA isolation experiments.
(B) PC3 parental or sorted subpopulations, western blotted for PODXL, NHERF1, EZRIN, and GAPDH as loading control for NHERF1.
(C) Quantitation from B, band intensity fold change (log10) between High-surface and Low-surface. Box-and-whiskers: dots, replicates; +,mean; midline, median; boundaries, quartiles; n=5-9 independent lysate preparation experiments.
(D) Enriched pathway maps determined from RNA-seq of PC3 subpopulations sorted for endogenous surface levels of PODXL. Enrichment was carried out using KEGG Pathway and MetaCore analyses. Y-axis denotes, top ten enriched pathways; X-axis, rich factor (percentage of targeted genes in each pathway); bubble colourmap, q-value; bubble size, total number of identified genes.
(E) Total 2D or 3D cell lysates from PC3 parental or sorted subpopulations. Western blotting of EMT-related markers (ESRP1/2, E-CAD, N-CAD, VIM, ZEB1) or GAPDH.
(F) Quantification from (E) across replicates. High-surface versus low-surface PODXL levels; intensity fold change (log10). Box-and-whiskers: dots, replicates; +, mean; midline, median; boundaries, quartiles; n=3-7 independent experimental isolations of cell lysates.
(G) Western blotting for PODXL or GAPDH as loading control in high-surface PODXL cells stably expressing scramble or *PODXL* shRNA.
(H) 3D phenotypes of high-surface PODXL cells expressing scramble or *PODXL* shRNA. Phase-contrast images from time-lapse series. Object outlines, yellow. Time, hours (h). Outlines (49-

60h) colourcoded by time and overlaid into single image (Composite). Dashed rectangle, area of interest.
(I) Heatmap, relative levels of user-defined behaviours in high-surface PODXL cells expressing scramble or *PODXL* shRNA (Round, Spread, Tunnel-forming), Z-score normalised values. p-values were calculated by Cochran-Mantel-Haenszel test, and Bonferroni adjusted, to compare proportion of each classification to control, greyscale as indicated. n=3 independent experiments, 4 technical replicates per condition, totalling 1,215-1,633 initial cysts per condition per experiment.
(J) Wound-healing 3D invasion in high-surface PODXL cells expressing scramble or *PODXL* shRNA examined by hourly time-lapse imaging. Representative phase-contrast images at 72h. Yellow line, initial wound position; red line, remaining wound region at $T_{max}^{1/2}$ closure. Arrowheads, invasive tunnels. White square, region of interest. Elapsed time, hours (h).
(K) Quantitation across experiments. Circles, technical replicates; triangles, average per independent experiment; colour, experiment number. Mean ± SD, n=4 independent experiments, with 4-8 technical replicates per experiment.
(L-M) Western blotting for EMT-related markers (E-CAD, VIM, ZEB1) or GAPDH as loading control in high-surface PODXL PC3 cells stably expressing scramble or *PODXL* shRNA. Quantification across replicates, mean ± SD, n=2 independent experimental isolations of cell lysates.
(N-O) Western blotting for EMT-related markers (E-CAD, VIM, ZEB1) or GAPDH as loading control in MDA-MB-231 cells stably expressing scramble or *PODXL* shRNA. Quantification across replicates, mean ± SE, n=3 independent experimental isolations of cell lysates.

Scale bars, 300 µm (H,J). p-values (F,K,O) (unpaired Student's 2-tailed t-test), ns, not significant,*P ≤ 0.05, **P ≤ 0.005, ***P ≤ 0.0005, ****P ≤ 0.0001.

**Figure S6.** U**biquitination controls PODXL cortical levels.**
(A) Cartoon, GFP-PODXL WT and ubiquitination mutant (K431/525/526/547R; 4K>R). SP, signal peptide; TM, transmembrane domain.
(B-C) Cell surface GFP levels in PC3 cells expressing either GFP or GFP-PODXL WT or 4K>R by flow cytometry. (B) Representative plot. (C) Quantitation, fluorescence intensity geometrical mean normalised to mean. Floating-bars plot: dots, experiments; midline, mean; n=3 independent experiments.
(D-G) Examination of EMT profile after expression of GFP or GFP-PODXL WT or 4K>R mutant. (D) Representative western blot for E-CAD, N-CAD, VIM, GFP and GAPDH as loading control for GFP. (E-G) Quantitation of markers across n=3 independent experimental isolations of cell lysates.
(H-I) PC3 cells cultured on plastic ('2D') co-expressing GFP-PODXL-FKBP 4K>R mutant with either FRB-BFP or FRB-Ubiquitin-BFP without (-) or with (+) dimeriser for 4 hours. (H) Representative staining in non-permeabilised cells with anti-GFP (magenta), fluorescent protein GFP (green) and TagBFP (blue). Scale bar, 25 µm. (I) Quantitation of Surface:Total GFP intensity ratio. Box-and-whiskers: 10-90 percentile; +, mean; dots, outliers; midline, median; boundaries, quartiles; n=124-714 independent cysts imaged/condition, 4 independent wells per experiment (n=1).

p-values (unpaired Student's 2-tailed t-test), ns, not significant,*P ≤ 0.05, **P ≤ 0.005, ***P ≤ 0.0005, ****P ≤ 0.0001

**Figure S7. Mapping the association of GAL3 with PODXL.**
(A-G) GFP-Trap immunoprecipitation performed on cells expressing GFP or GFP-PODXL WT and mutants.
(A) Cartoon, GFP-PODXL domains and mutants. SP, signal peptide; TM, transmembrane domain; FBM, FERM-binding-motif; PBM, PDZ-binding-motif. N, N-glycosylated asparagine residues. K, ubiquitinated lysine residues. WT, wild-type; ΔPBM, PDZ-binding motif mutant; FBM*, FERM-binding motif mutant; 5N>Q, N-glycosylation mutant; 4K>R, ubiquitination mutant; ΔIC, intracellular domain deletion; ΔEC, extracellular domain deletion. (B,D,F) Representative western blots for GAL3, GFP and GAPDH (as loading control for GFP) antibodies in indicated conditions. (C,E,G) Quantitation of endogenous GAL3 with GFP-PODXL mutant, fold change to mean. Floating-bars plots: dots, experiments; midline, mean; boundaries, min and max values; n=3 independent experiments.
(H) Cartoon, TagRFP-T-GAL3 WT and R186S (N-acetyllactosamine binding-deficient mutant). CRD, carbohydrate-recognition domain. R, arginine; S, serine.
(I) GFP-Trap immunoprecipitation performed from cells co-expressing either GFP or GFP-PODXL WT and either TagRFP-T, TagRFP-T-GAL3 WT or R186S mutant. Western blot for GAL3, GFP and GAPDH (as loading control) antibodies.
(J) Quantitation, GFP-PODXL interaction with TagRFP-T-GAL3 WT versus R186S. Floating-bars plot, line at median, n=3 independent experiments.
(K-M) Phenotype of PC3 cells expressing GFP-PODXL WT or intracellular domain deletion (ΔIC) mutant in cyst assay (K) or invasion assay (L-M). (K) Heatmap, relative levels of user-defined behaviours (Round, Spread, Tunnel-forming), Z-score normalised values. p-values were calculated by Cochran-Mantel-Haenszel test, and Bonferroni adjusted, to compare proportion of each classification to control, greyscale as indicated. n=3 independent experiments, 4 technical replicates per condition, totalling 415-569 initial cysts per condition per experiment. (L) Representative phase-contrast images at 66h. Yellow line, initial wound position; red line, remaining wound region at $T_{max}^{1/2}$ closure. Elapsed time, hours (h). (M) Quantitation across experiments. Circles, technical replicates; triangles, average per independent experiment; colour, experiment number. Mean ± SD, n=3 independent experiments with 3 technical replicates per experiment.
(N) *LGALS3* mRNA expression in parental PC3 and High-surface and Low-surface PODXL-sorted subpopulations. Mean ± s.d; n=4 independent RNA isolations per condition.
(O-P) Galectin-3 expression in media and lysates from parental PC3, High-surface and Low-surface PODXL-sorted subpopulations. Mean ± S.D; n=3 independent lysate and media isolations per condition.
(Q) Phenotype of PC3 cysts depleted for endogenous *PODXL* and rescued with RNAi-resistant GFP-PODXL WT or non-N-glycosylated (5N>Q) mutant. Heatmap, relative levels of user-defined behaviours (Round, Spread, Tunnel-forming), Z-score normalised values. p-values were calculated by Cochran-Mantel-Haenszel test, and Bonferroni adjusted, to compare proportion of

each classification to control, greyscale as indicated. n=3 independent experiments, 4 technical replicates per condition, totalling 515-1,188 initial cysts per condition per experiment.

Scale bars, 300 μm (L,S). p-values (unpaired Student's 2-tailed t-test), ns, not significant,*P ≤ 0.05, **P ≤ 0.005, ***P ≤ 0.0005, ****P ≤ 0.0001

**Figure S8. Mutual antagonistic regulation of GAL3 and PODXL.**
(A-B) Surface PODXL levels by flow cytometry in Low-surface (A) or High-surface (B) endogenous PODXL expressing cells, stably expressing either Scramble, *PODXL* or *GAL3* shRNA. Geometrical mean, surface PODXL fluorescence intensity. Floating bar chart: dots, experiments; midline, mean; boundaries, min and max values; n=3 (A) or n=4 (B) independent experiments, respectively.
(C-E) Surface PODXL levels by flow cytometry in High-surface PODXL (C-D) or parental (E) PC3 cells without (control) or with prior 30min stimulation with 1.5 μM recombinant GAL3 (ReGAL3). (C) Representative plot for PC3-high. (D-E) Quantitation, floating-bars plot: dots, experiments; midline, mean; boundaries, min and max values; n=3 independent experiments per condition.
(F-G) Western blotting for extracellular GAL-3 in media or GAL3, PODXL and GAPDH as loading control in parental PC3 cells stably expressing scramble or *PODXL* shRNA. Quantification across replicates, mean ± SD, n=3 independent experimental isolations of cell media.(H-J) Recombinant-GAL3 internalisation in endogenous GAL3-depleted PC3 cells (shRNA), without (scramble) or with PODXL co-depletion (shRNA). (H) Schema. (I) Representative images of cells stained for GAL3 (black). Cell outline (Magenta dashed line); nucleus outline (blue line). (J) Quantitation of GAL3 internalisation at indicated times. Data, Log2-normalised to control condition at 0 min. Box-and-whiskers: 10-90 percentile; +, mean; dots, outliers; midline, median; boundaries, quartiles; n=3 independent replicates, 204-568 total cells imaged per condition.

Scale bars, 12.5 μm. p-values (unpaired Student's 2-tailed t-test), ns, not significant,*P ≤ 0.05, **P ≤ 0.005, ***P ≤ 0.0005, ****P ≤ 0.0001.

**Figure S9. Characterisation of ITGB1 surface levels upon PODXL or GAL3 alteration.**
(A) Western blotting for ITGB1 and GAPDH as loading control in parental PC3 cells or subpopulations sorted for High or Low endogenous surface PODXL levels. Top, representative image, bottom quantification across replicates, mean ± SD, n=3 independent experimental isolations of cell lysates.
(B-F) Representative plot of cell surface ITGB1 and active ITGB1 levels in parental PC3 cells or subpopulations sorted for High or Low endogenous surface PODXL levels by flow cytometry. Quantitation, fluorescence intensity geometrical mean normalised to mean. Floating-bars plot: dots, experiments; midline, mean; n=3 independent experiments.
(G-L) Representative plot of cell surface ITGB1, active ITGB1 and GFP levels in PC3 cells expressing GFP, GFP-PODXL WT or GFP-PODXL 4>R mutant by flow cytometry. I Quantitation, fluorescence

intensity geometrical mean normalised to mean. Floating-bars plot: dots, experiments; midline, mean; n=3 independent experiments.
(M-P) Representative plot of cell surface ITGB1 and active ITGB1 levels in PC3 cells stably expressing GAL3 shRNA or scramble control by flow cytometry. Quantitation, fluorescence intensity geometrical mean normalised to mean. Floating-bars plot: dots, experiments; midline, mean; n=3 independent experiments.

p-values (unpaired Student's 2-tailed t-test), ns, not significant,*P ≤ 0.05, **P ≤ 0.005, ***P ≤ 0.0005, ****P ≤ 0.0001.

**Figure S10. Cortical Podxl regulates association with the ECM.**

(A-D) Cell attachment to ECM in PC3 cells with (A-B) Low-surface or High-surface endogenous PODXL or (C-D) endogenous PODXL-depleted cells (shRNA, +) expressing GFP or GFP-PODXL WT or 4K>R. Cells were seeded on plates pre-coated with 20 µL of: 50% Matrigel or 10 µg/mL of Laminin, Fibronectin or Collagen-I. Non-adherent cells removed by washing (after 0.5, 1, 2 or 4 hours) and fixed. Cells, detected by F-actin labelling (black, inverted image) were counted. (A,C) Representative images. Scale bars 300 µm. (B,D) Graphs displaying number of attached cells along time within each of the matrixes/conditions, symbols, mean; error bars, SD; n=1 independent experiment with 4 technical replicates. 1500 cells seeded per well.
(E) Confocal images of endogenous PODXL-depleted PC3 cysts (shRNA, +) expressing GFP, GFP-PODXL WT or 4K>R, fixed 3 days after plating. Cysts were stained for Collagen-IV (magenta), F-actin (grey) and nuclei (blue). GFP expression (green). Scale bars, 50 µm.
(F) Cyst assay, PC3 cells stably expressing Scramble shRNAs without (IgG control) or with b1-integrin blockade (AIIB2, blocking antibody). Phase-contrast time-lapse images of 3D structures. Cysts outlines, yellow. Elapsed time (hours, h), bottom corner. Outlines from each hour between 37-48h colour-coded by time in rainbow look-up table overlaid on single image (Composite). White rectangle, region of interest.
(G) Heatmap, proportions of distinct phenotypes (Round, Spread, Tunnel-forming), Z-score normalised values. p-values were calculated by Cochran-Mantel-Haenszel test, and Bonferroni adjusted, to compare proportion of each classification to control, greyscale as indicated. n=3 independent experiments, 2 technical replicates per condition, totalling 154-281 initial cysts per condition per experiment.

**Figure S11. Additional PODXL and GAL3 characterisation in PC3 tumours.**

(A) Schema, intraprostatic transplantation. PC3 cells manipulated for PODXL expression or surface levels were transplanted into the prostate of 7-week old CD-1 nude male mice. Tumour growth and metastatic events were analysed at endpoint of 8 weeks.
(B) Serial sections of the primary tumour (prostate) and metastases (Lymph Node, Abdominal Wall, Epidydymal Fat) from mice intraprostatically transplanted with PC3 shRNA Scramble cells, stained for Hematoxylin & Eosin, or either PODXL or GAL3. Scale bar, 100um. Arrowhead, cortical PODXL. Mouse ID as indicated.

**Figure S12. Additional clinical associations in GAL3-PODXL expression-divided patient groups.**
(A-D) *PODXL* mRNA expression in (A-B) normal prostate, primary tumour and metastasis or (C-D) primary tumours in disease-free or recurred patients.
(E-F) Disease-free survival (% patients, over months) in patient cohorts of high (quartile 4, Q4) versus not high (quartiles 1-3, Q1-3) *PODXL* mRNA expression.
(G-J) *LGALS3* mRNA expression in (G-J) normal prostate, primary tumour and metastasis or (K, L) primary tumours in disease-free or recurred patients.
(M,N) Disease-free survival (% patients, over months) in patient cohorts of low (quartile 1, Q1) versus not low (quartiles 2-4, Q2-4) *LGALS3* mRNA expression.

Box-and-whiskers graphs (A, B, C, D, G, H, I, J, K, L): 10-90 percentile; +, mean; dots, outliers; midline, median; boundaries, quartiles; number of patients in figure. Sample numbers indicated in on graphs.

Statistical analyses used were Kruskal–Wallis Test (A, B, G, H, I, J), Welch's t-test (C, D, K,L), or Log-rank t-test (E, F, M, N). p values: ns, not significant,*P ≤ 0.05, **P ≤ 0.005, ***P ≤ 0.0005, ****P ≤ 0.0001. Datasets: Glinsky (*78*), Grasso (GSE35988), Lapointe (GSE3933), Taylor (GSE21032), TCGA (PRAD), Tomlins (GSE6099), Varambally (GSE3325).

**Figure S13. Uncropped Western blots from main figures.**
Uncropped membranes related to figures as annotated.

**Table S1. mRNA expression and copy number of *PODXL* in cancer lines from Cancer Cell Line Encyclopedia**

**Table S2. SgRNA and shRNA target sequences**

**Table S3. Antibodies**

**Movie S1. Wound healing invasion of PC3 shRNA Scramble vs PODXL KD cells.**
Wound-healing motility in 3D invasion of PC3 Scramble (left) or PODXL KD (right) cells assays examined by hourly time-lapse imaging over 72h.

**Movie S2. Wound healing invasion of MDA-MB-231 shRNA Scramble vs PODXL KD cells.**
Wound-healing motility in 3D invasion of MDA-MB-231 Scramble (left) or PODXL KD (right) cells examined by hourly time-lapse imaging over 48h.

**Movie S3. 3D cyst invasion of PC3 cells shRNA Scramble vs PODXL KD.**
3D culture of PC3 Scramble (left) or PODXL KD (right) cysts examined by hourly time-lapse imaging over 72h.

**Movie S4. 3D cyst invasion of MDA-MB-231 cells shRNA Scramble vs PODXL KD.**
3D culture of MDA-MB-231 Scramble (left) or PODXL KD (right) cysts examined by time-lapse imaging every half an hour over 192h.

**Movie S5. 3D cyst invasion of PC3 cells with different surface levels of PODXL.**
3D culture of PC3 Parental (left) or cells sorted for low (middle) or high (right) surface levels of PODXL, grown as cysts examined by hourly time-lapse imaging over 60h.

**Movie S6. Wound healing invasion of PC3 cells with different surface levels of PODXL.**
Wound healing of PC3 Parental (left) or cells sorted for low (middle) or high (right) surface levels of PODXL, examined by hourly time-lapse imaging over 120h.

**Movie S7. ECM-tunnel lined by collagen fibres.**
Animated 3D reconstruction of confocal images using IMARIS software showing cells within ECM-tunnel structures lined by collagen fibres. Collagen-IV (white), GFP-PODXL 4K>R (green), F-actin (red) and nuclei (blue).

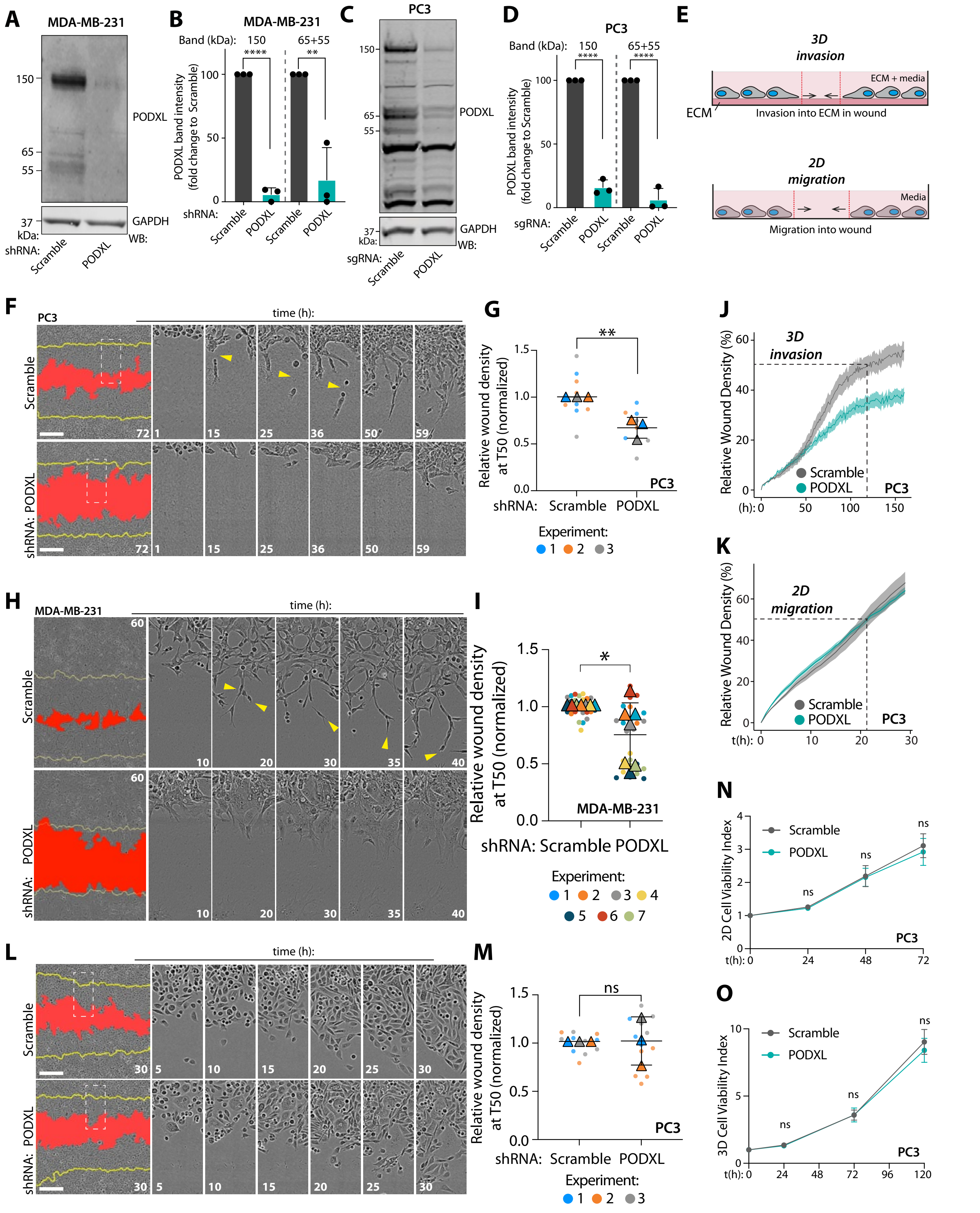


**Supplementary Figure 1**

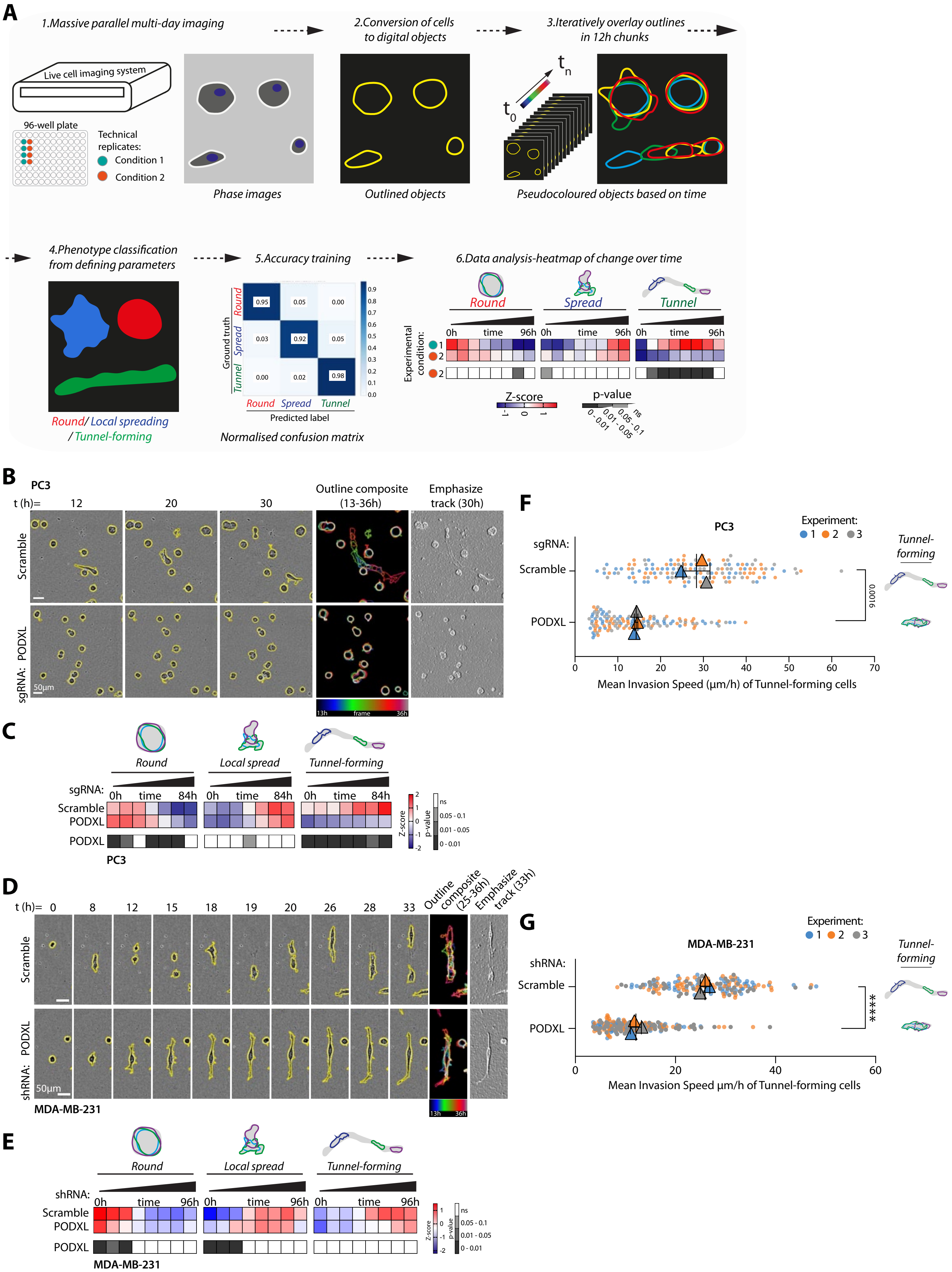


**Supplementary Figure 2**

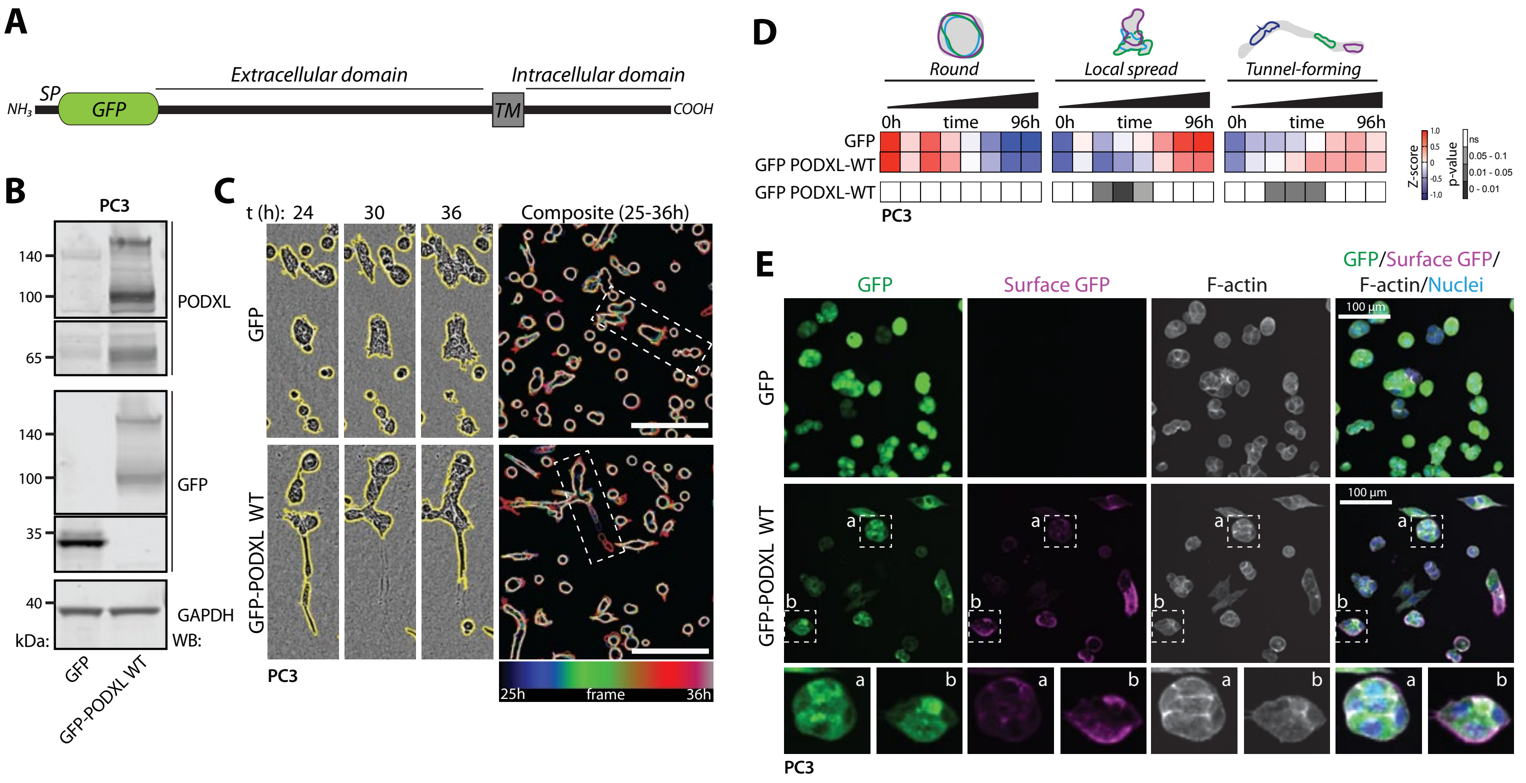


**Supplementary Figure 3**

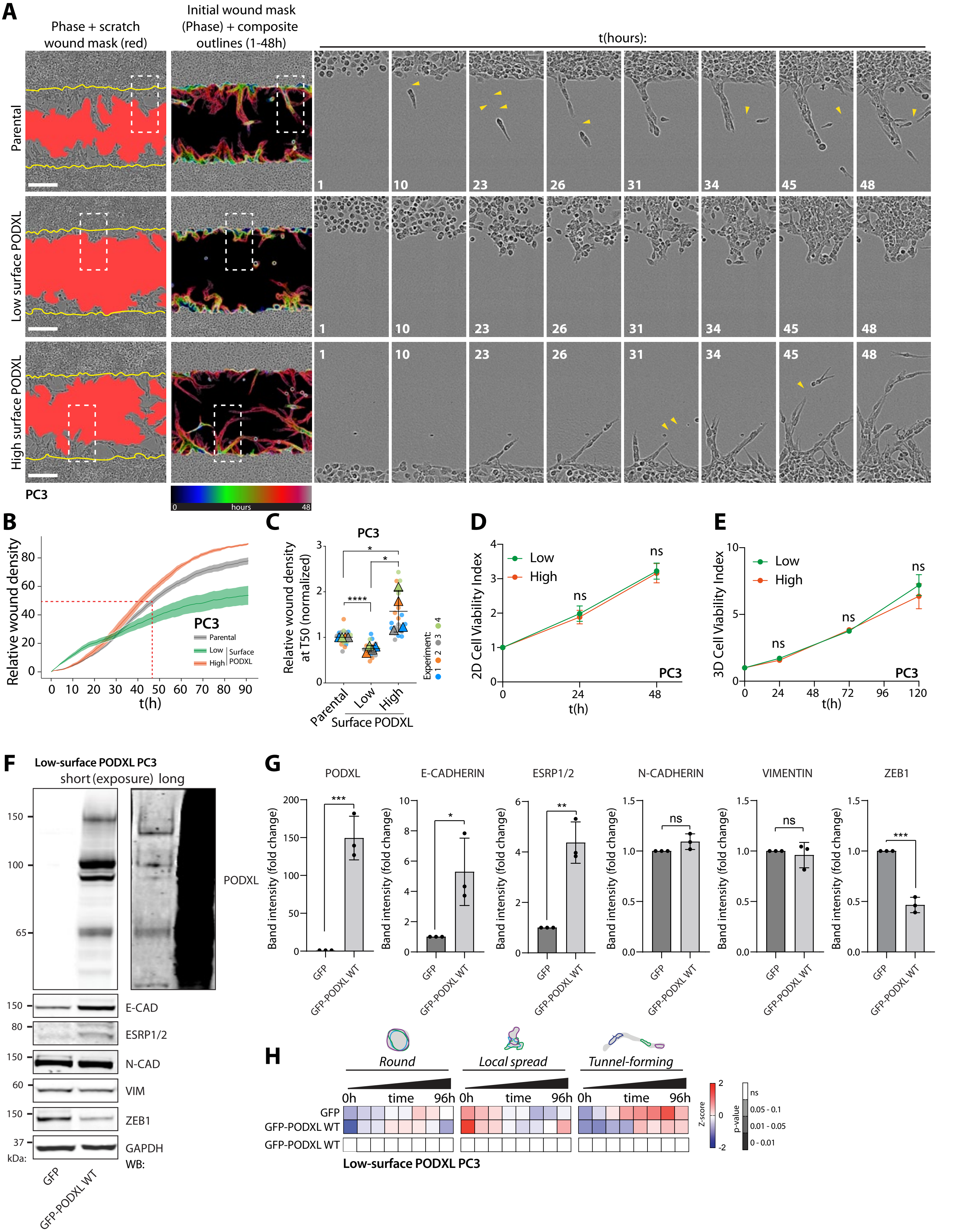


Supplementary Figure 4

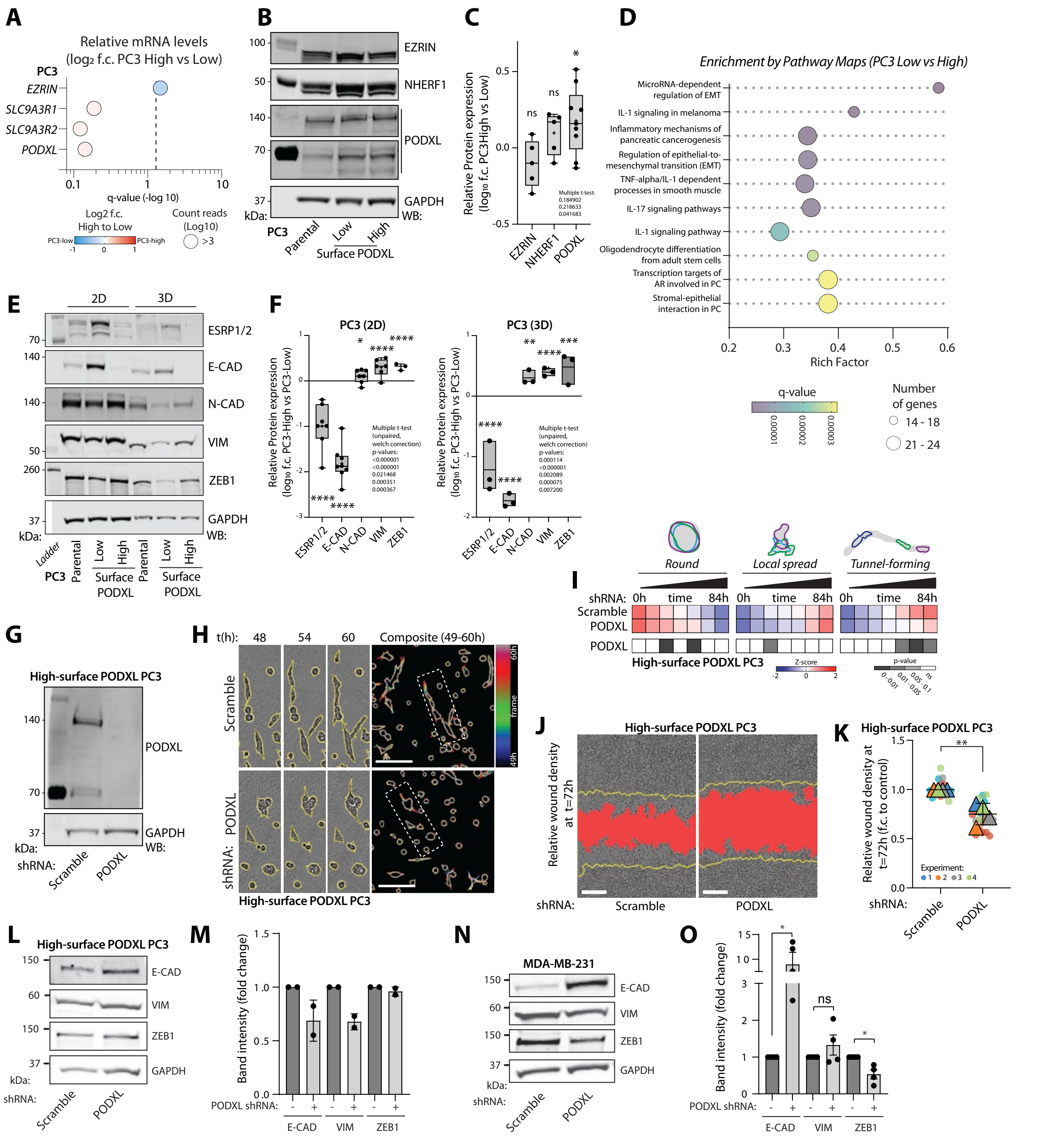


**Supplementary Figure 5**

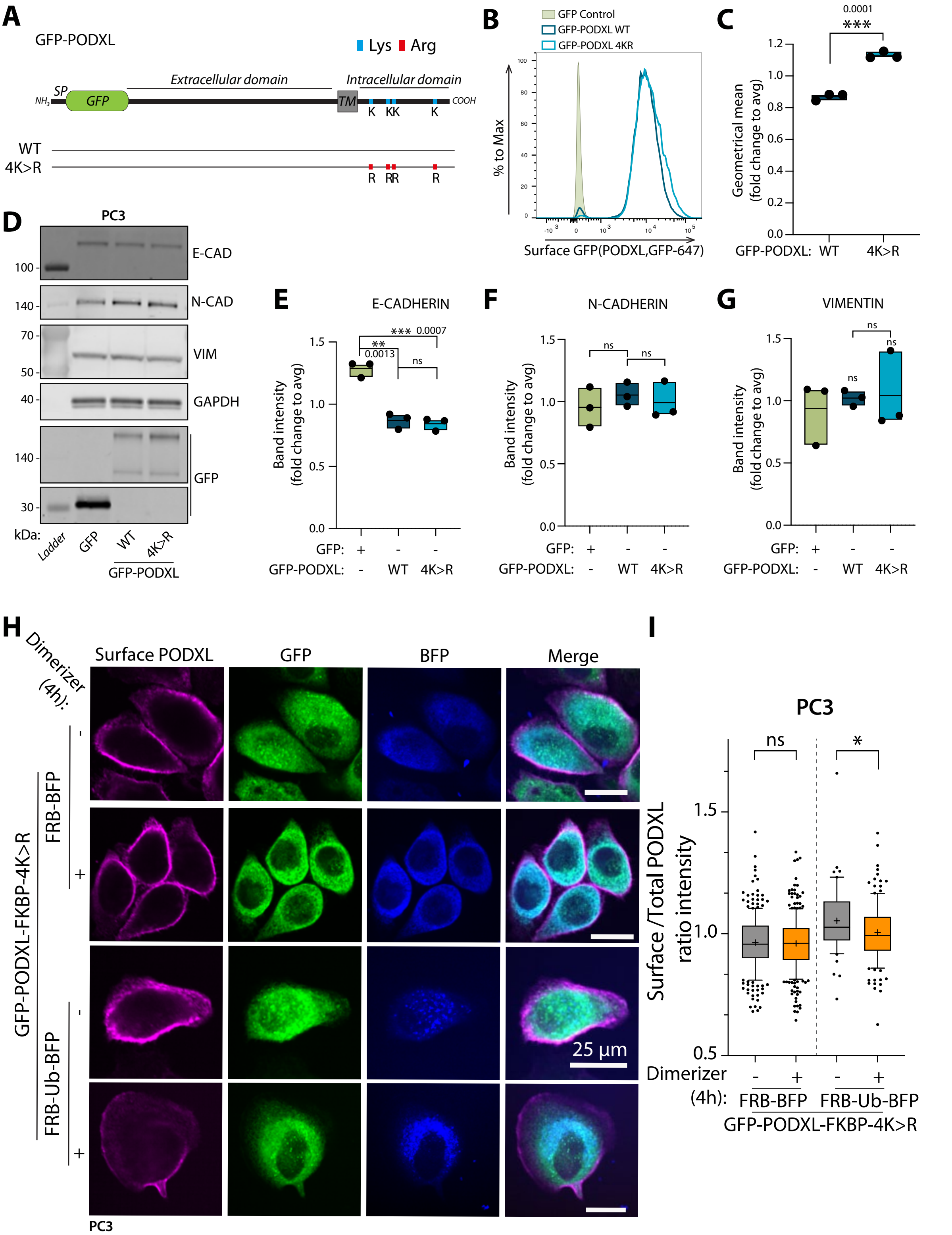


**Supplementary Figure 6**

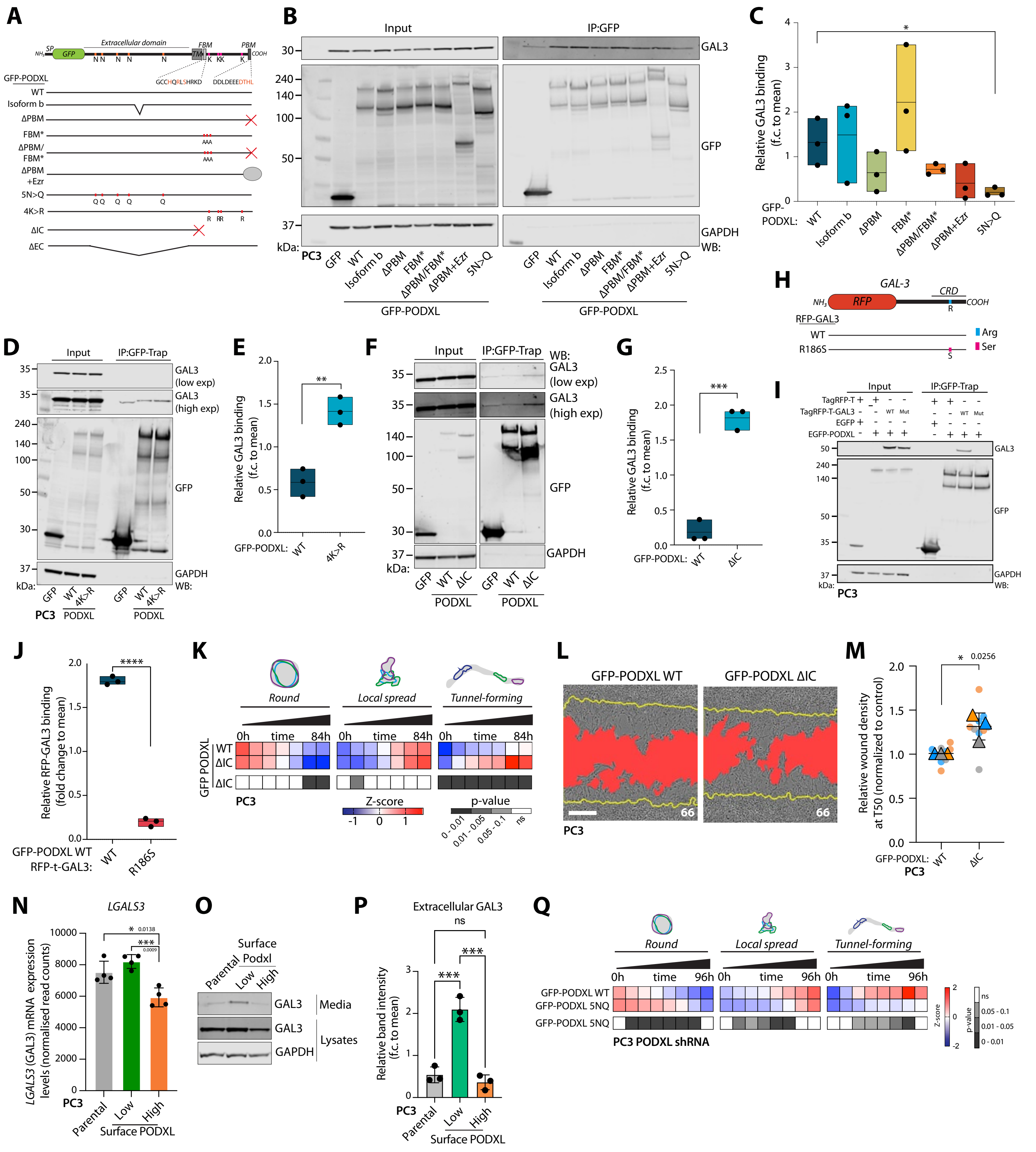


**Supplementary Figure 7**

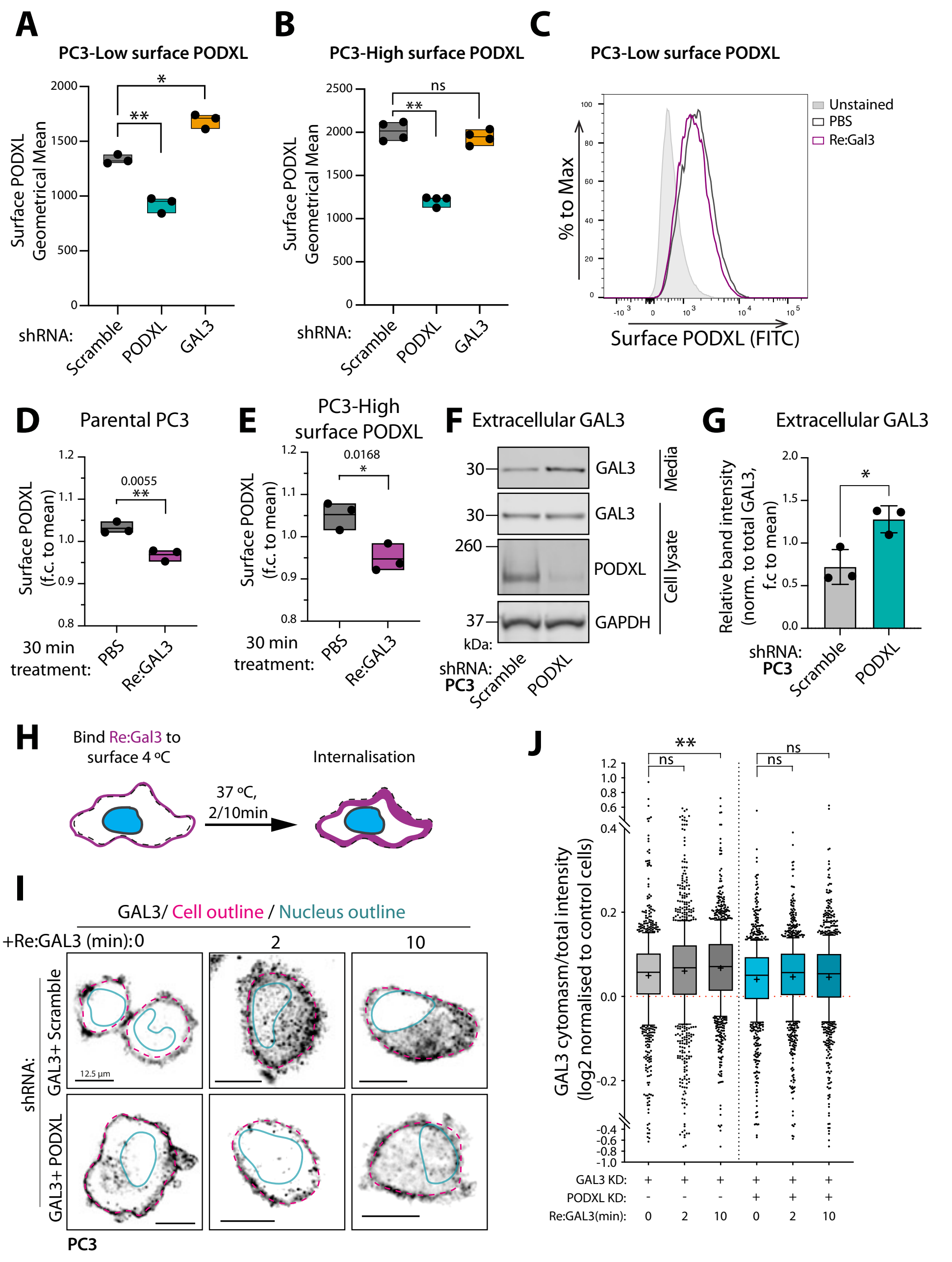


**Supplementary Figure 8**

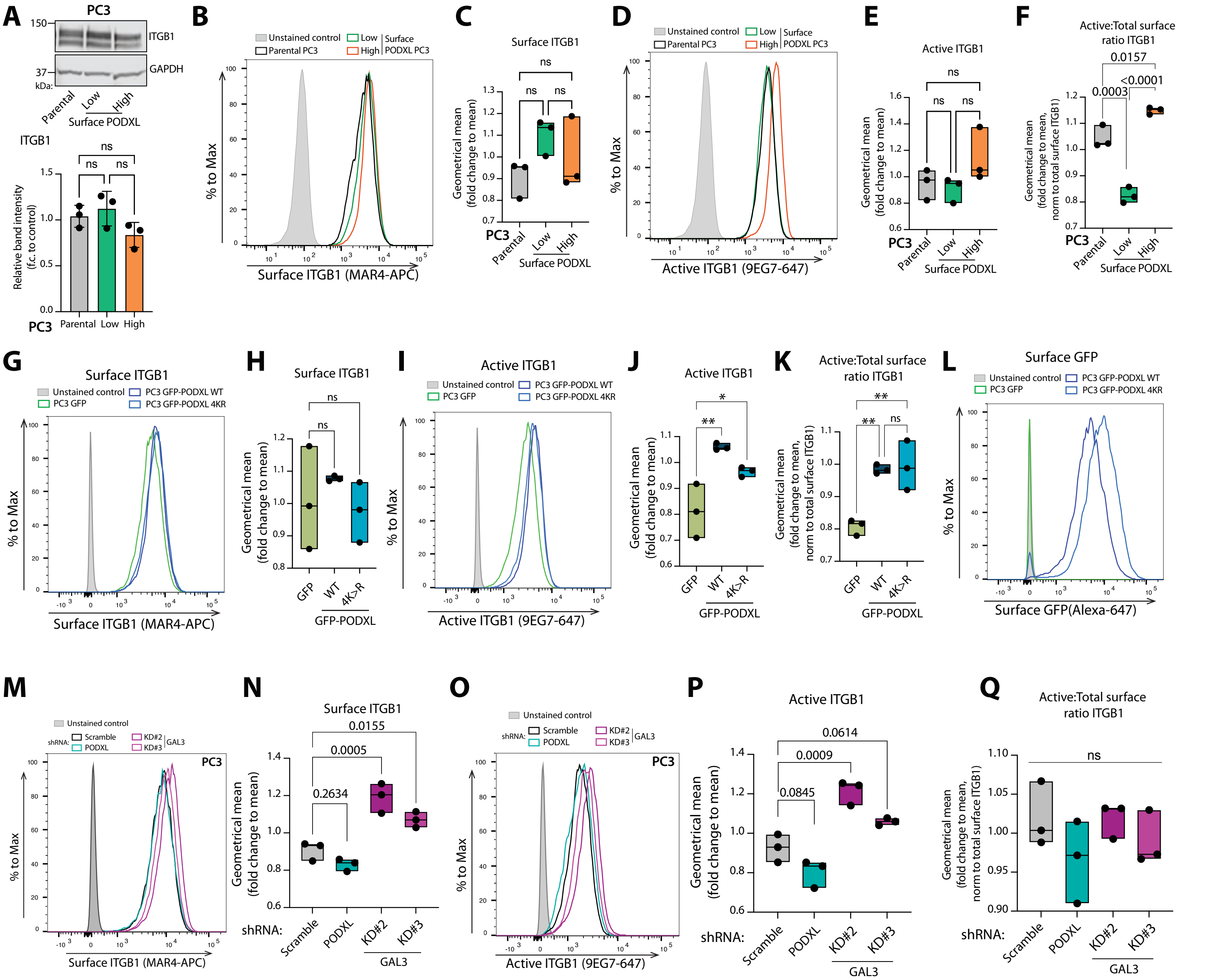


**Supplementary Figure 9**

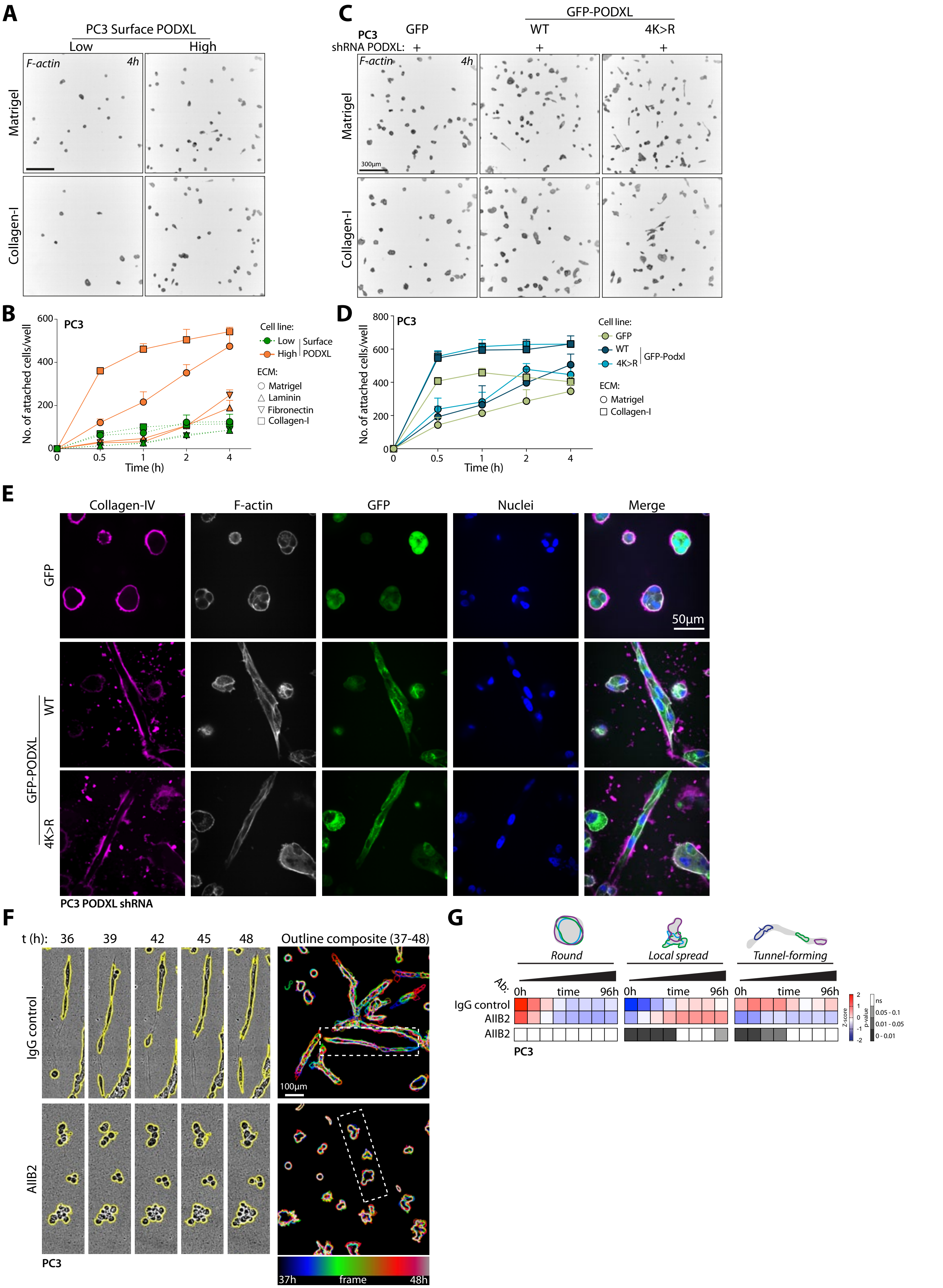


Supplementary Figure 10

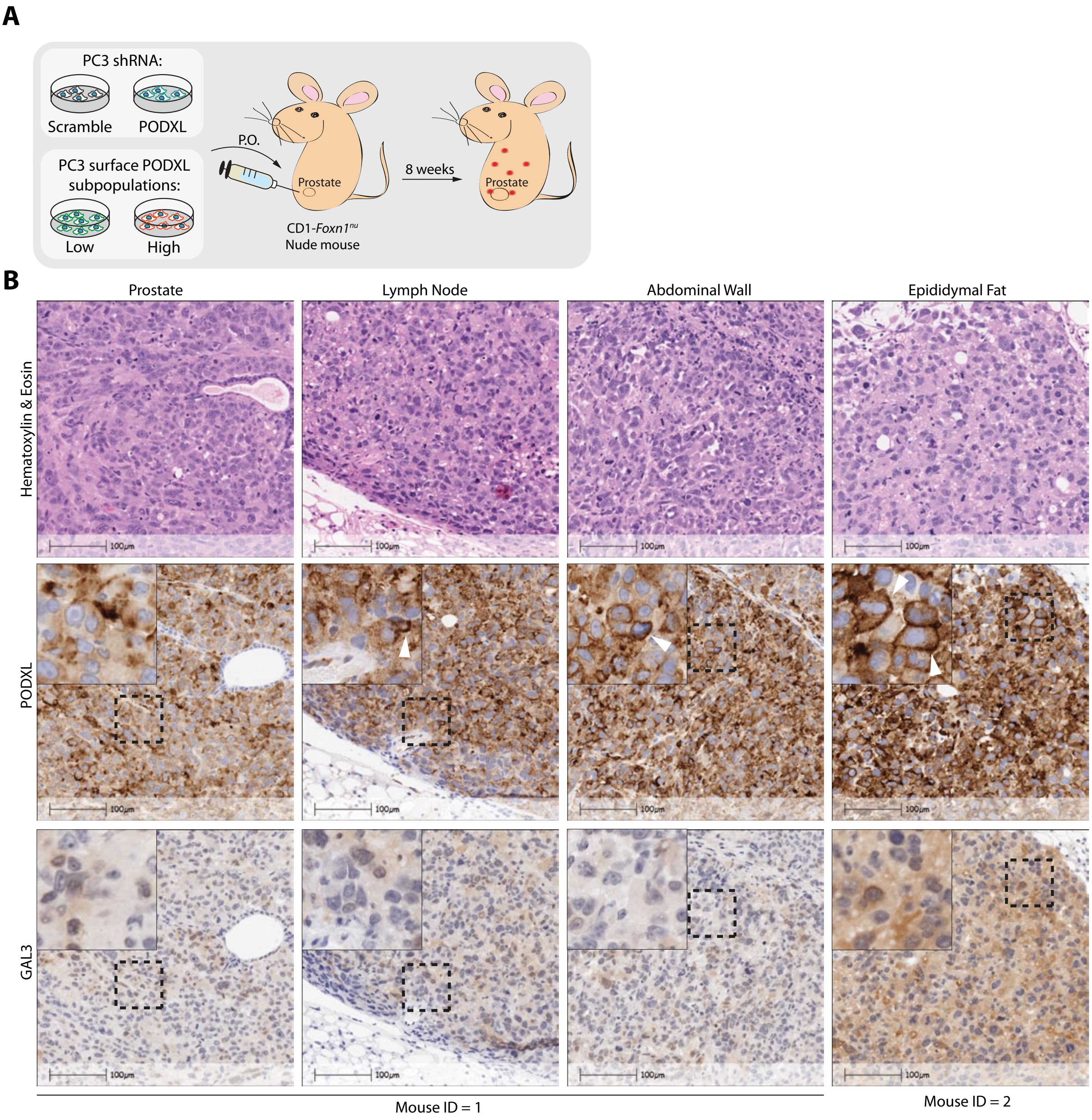


**Supplementary Figure 11**

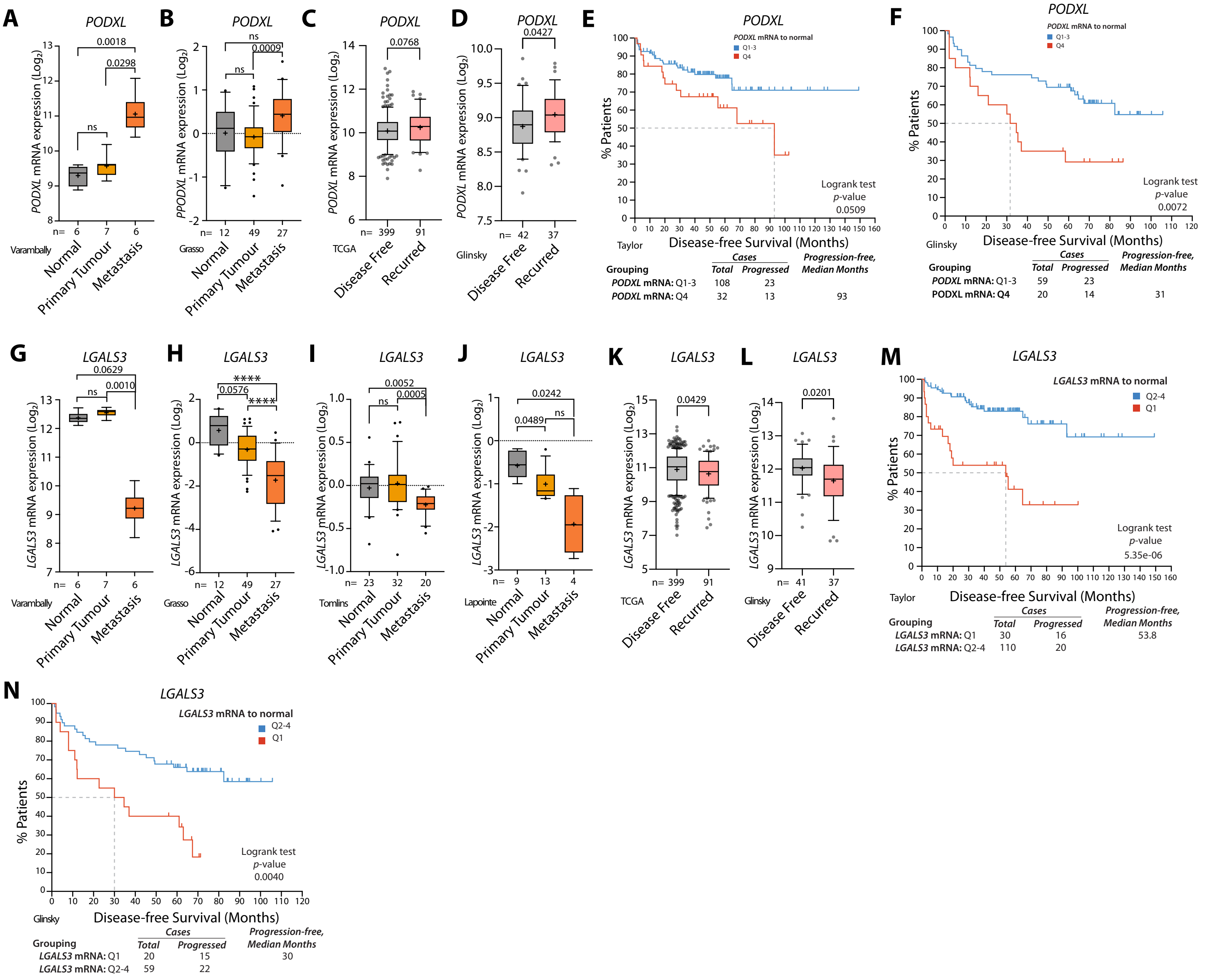


**Supplementary Figure 12**

Figure 1C:

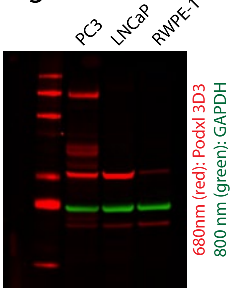


Figure 1F:

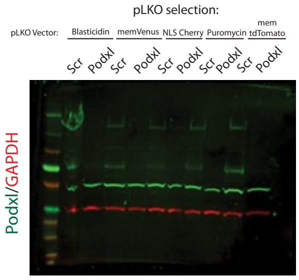


Figure 2G:

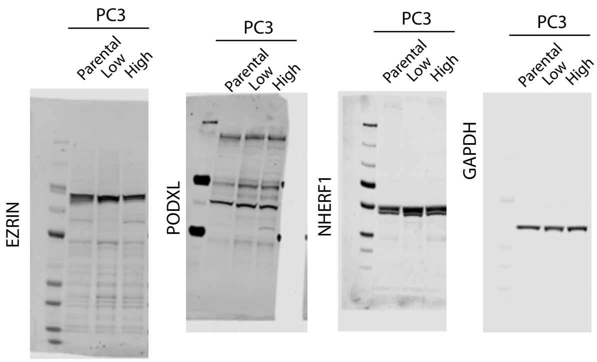


Figure 3A:

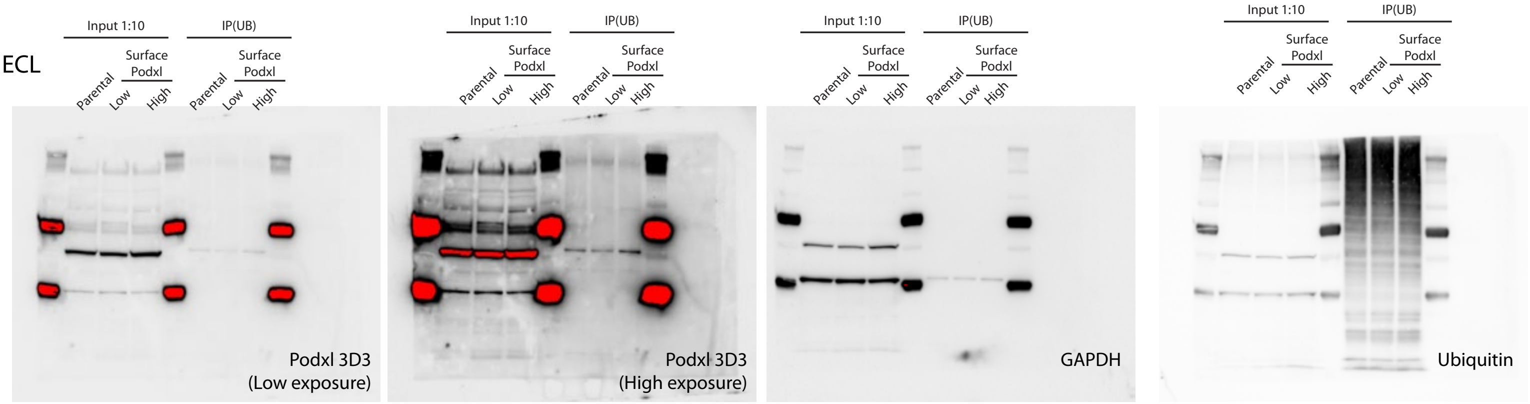


Figure 3C:

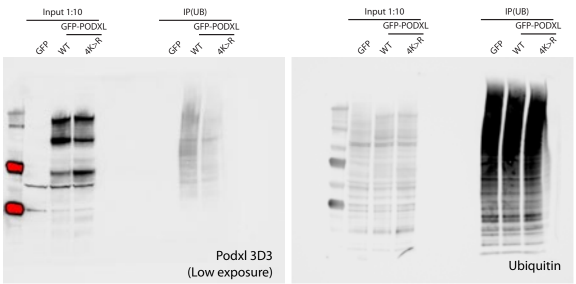


Figure 3J:

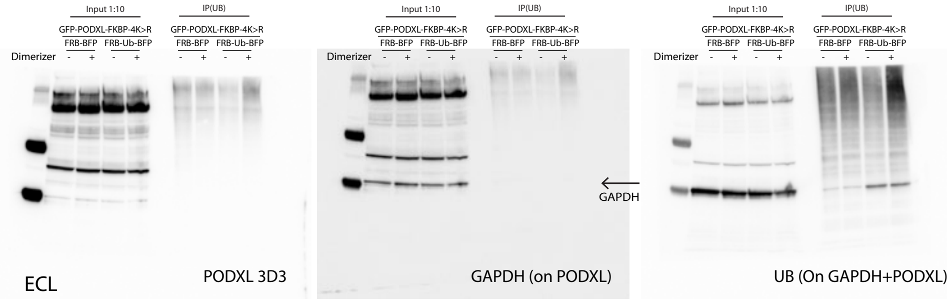


Figure 4E (and Supplementary 5B):

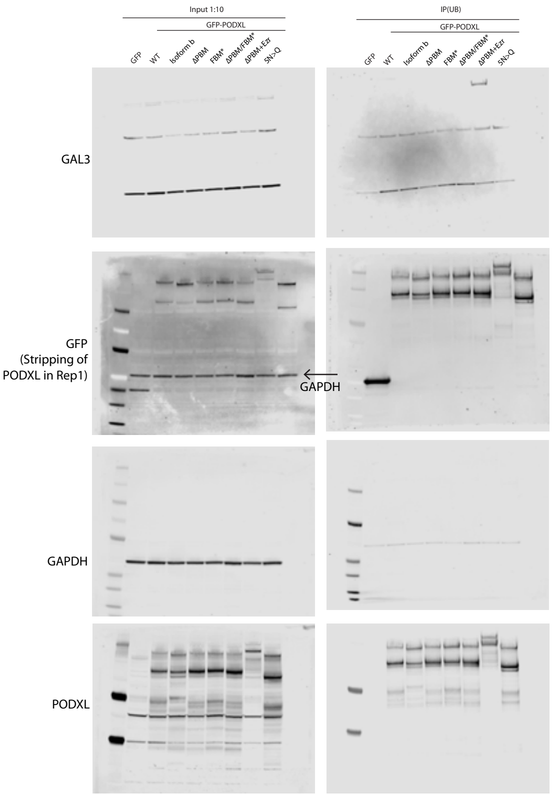


Figure 5A:

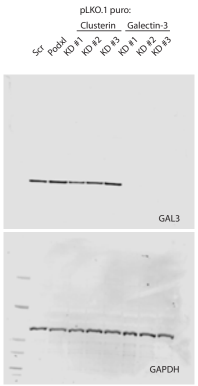


Figure 5G:

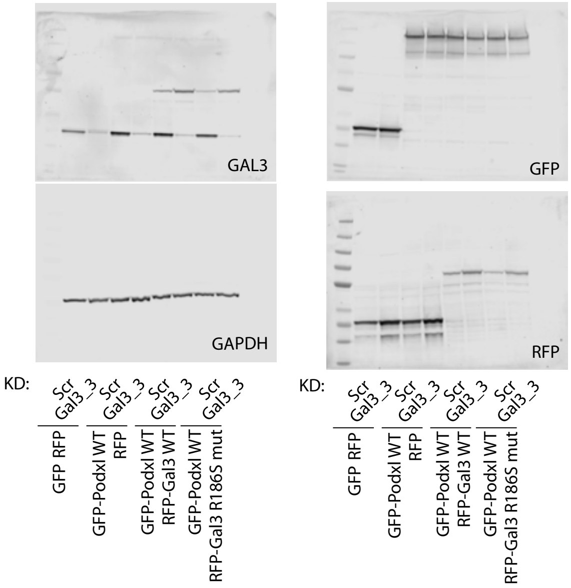

#### Figure 6I:

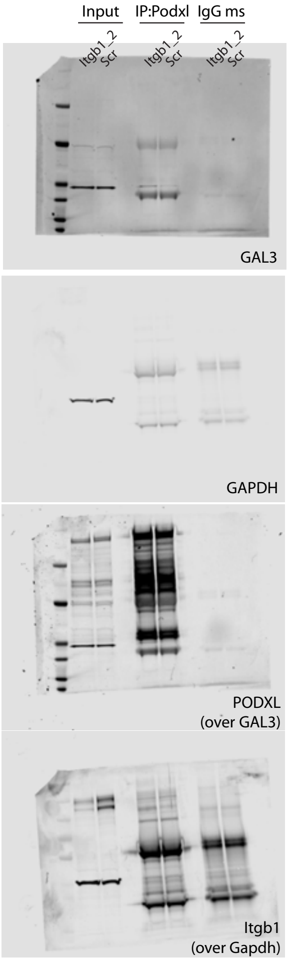


#### Figure 6M:

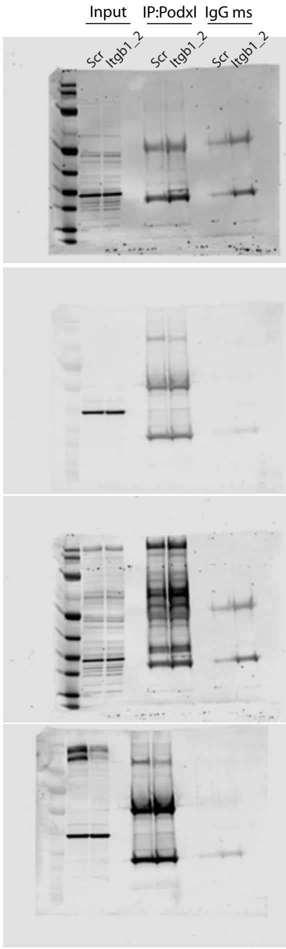


#### Figure S1A:

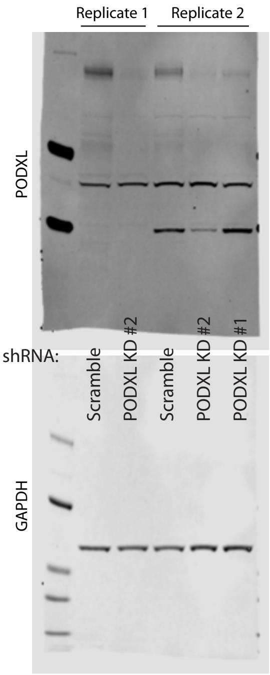


#### Figure S1C:

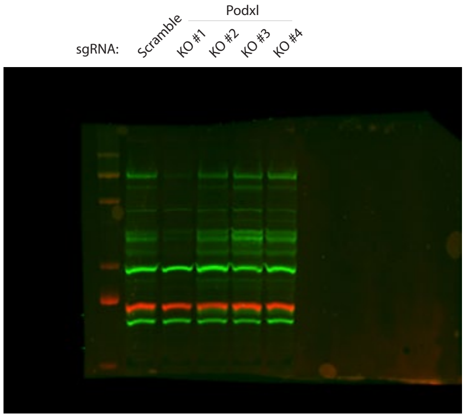


680nm (red): GAPDH
800 nm (green): Podxl 3D3

#### Fig. S2C:

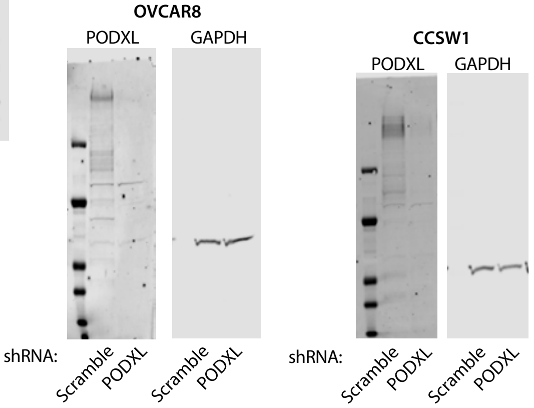


#### Figure S3B:

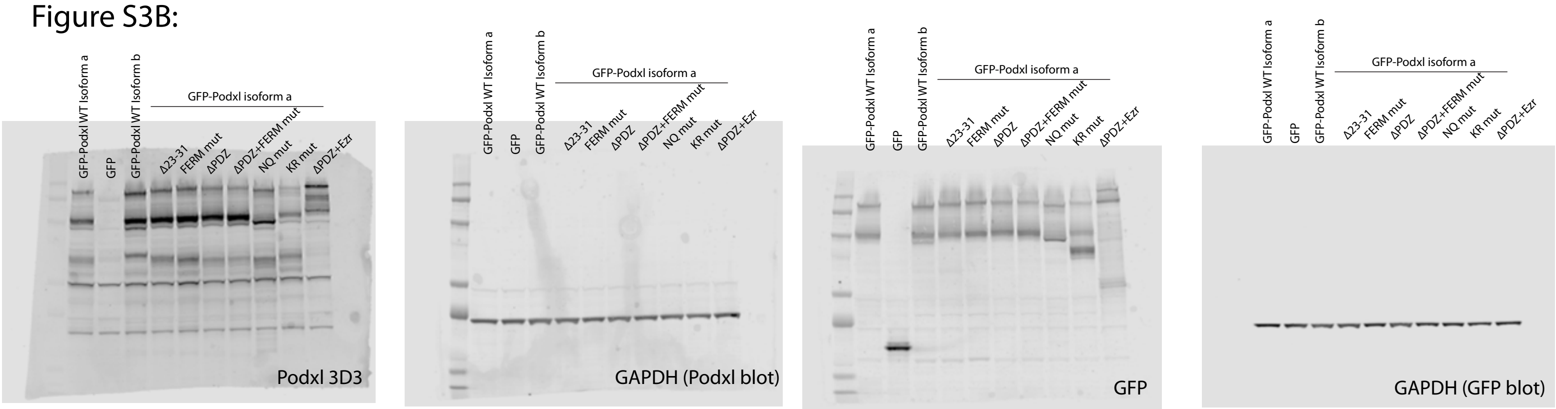


#### Figure S4F:

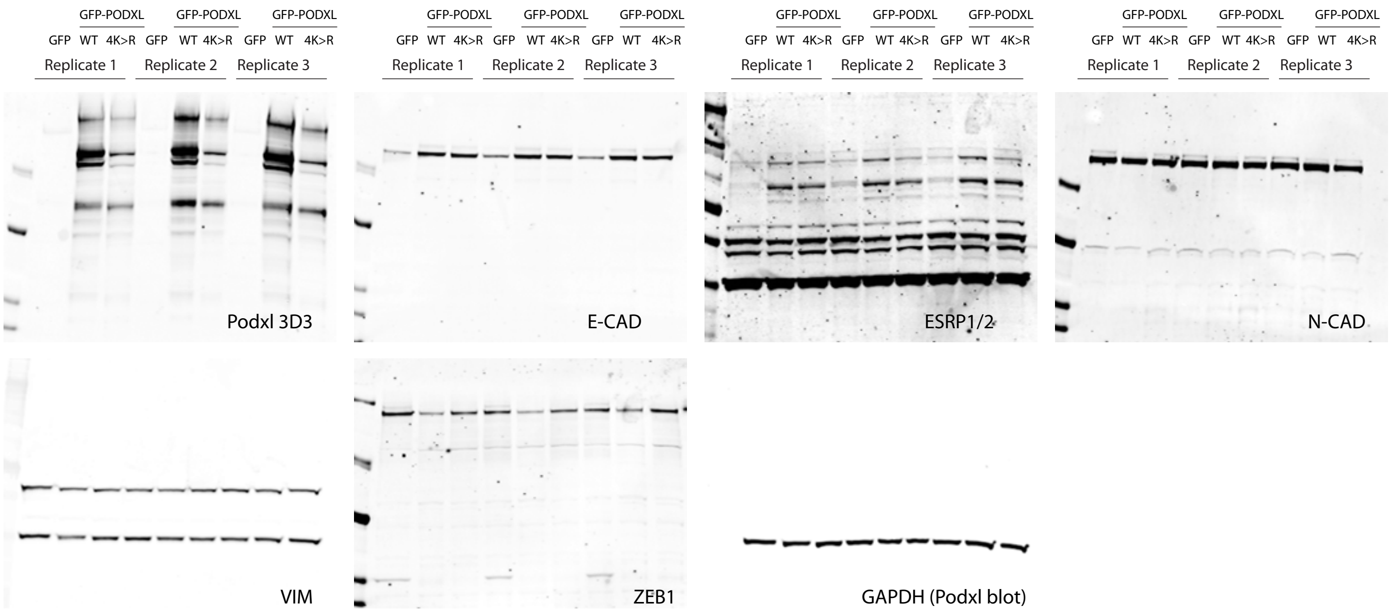

### Figure S5C:
### (In figure, rep 3)

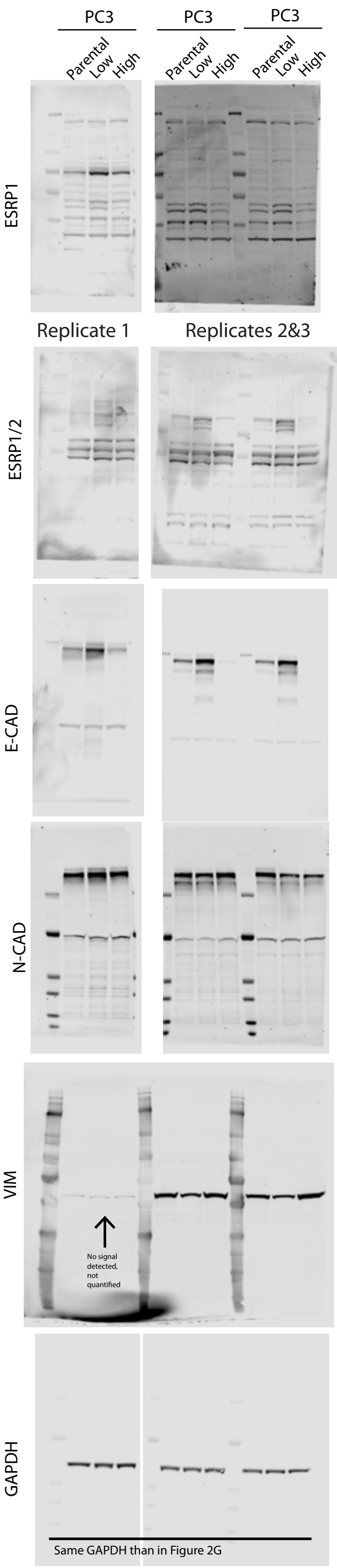


### Figure S5E:

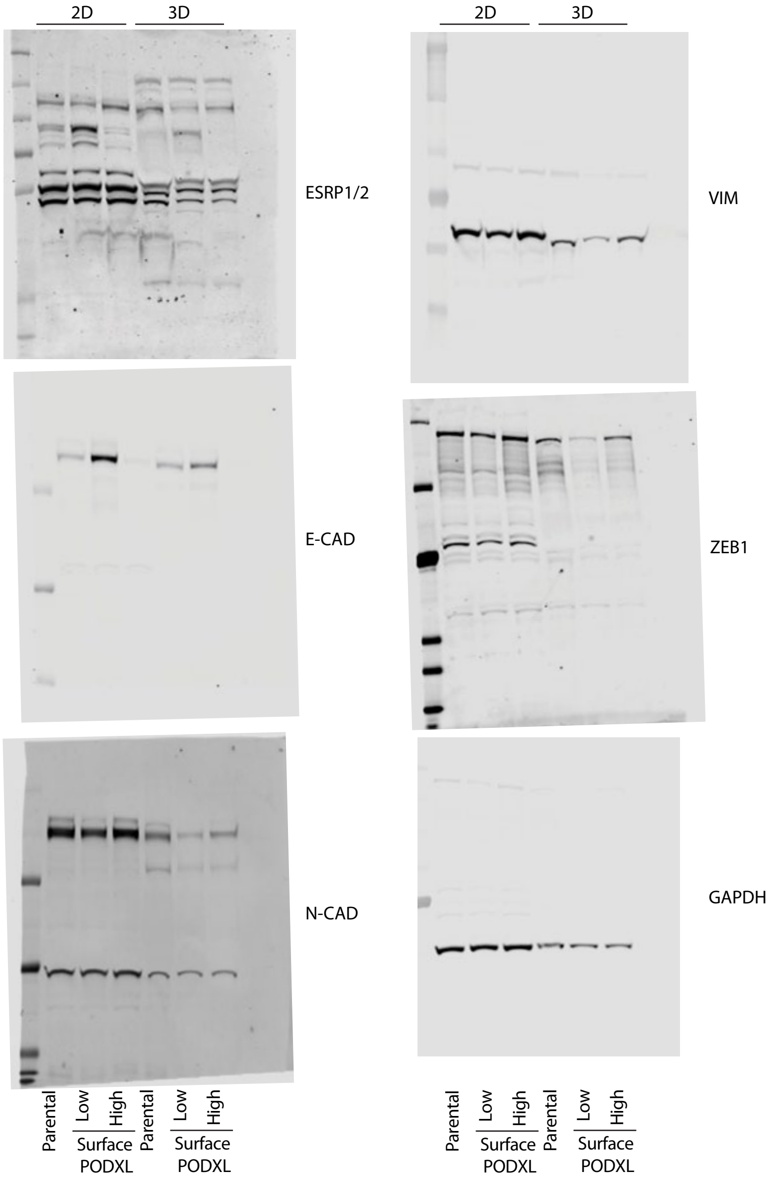


### Figure S5G:

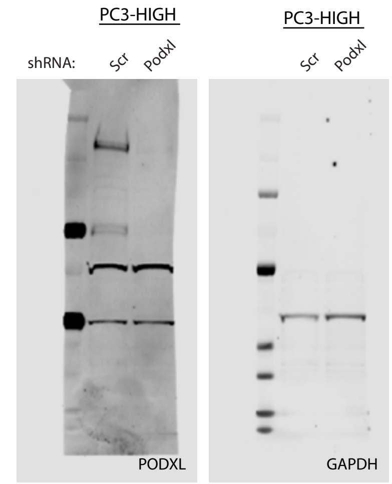

#### Figure S5L:

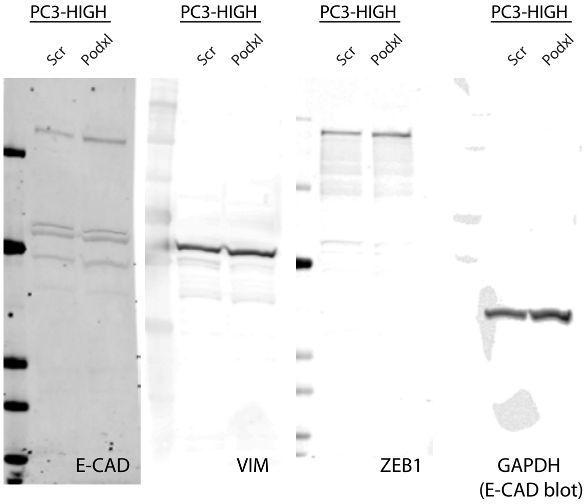


#### Figure S5N:

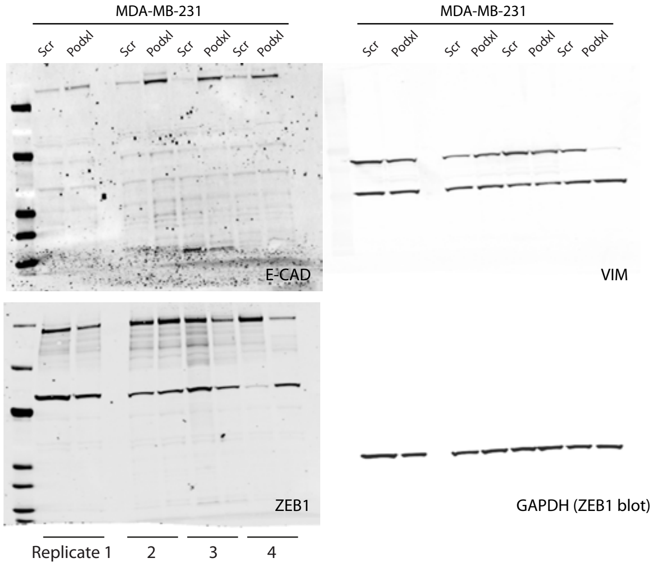


#### Figure S6D:
#### (In figure, rep 1)

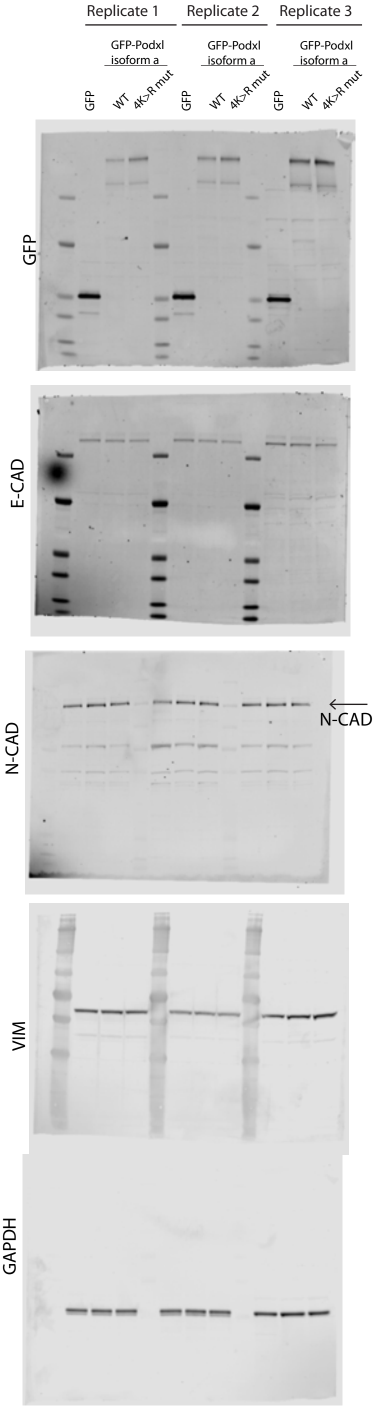


#### Figure S7B:

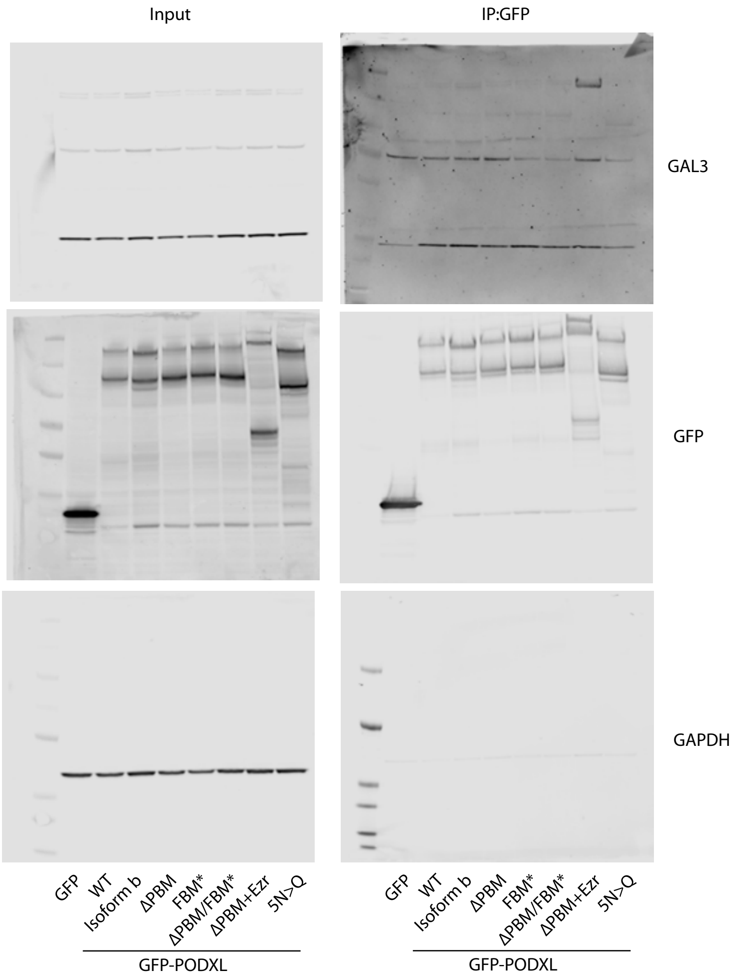


#### Figure S7D:

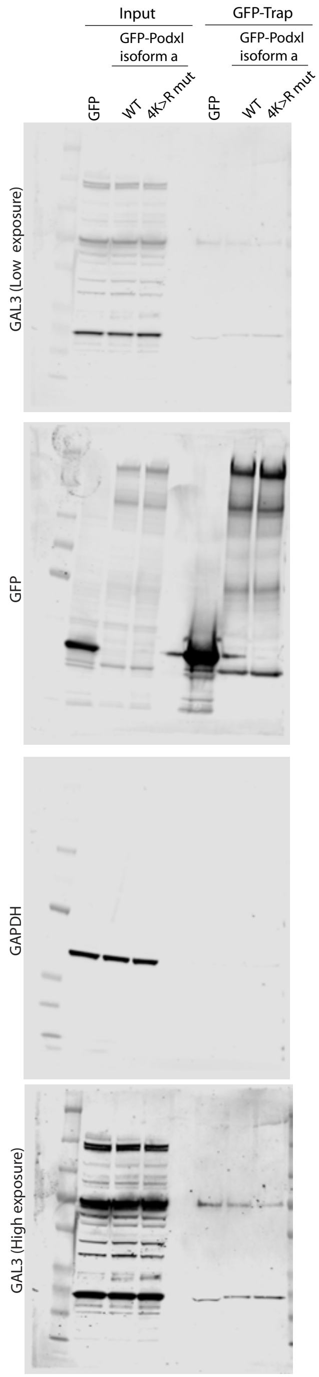

#### Figure S7F:

Replicate 1

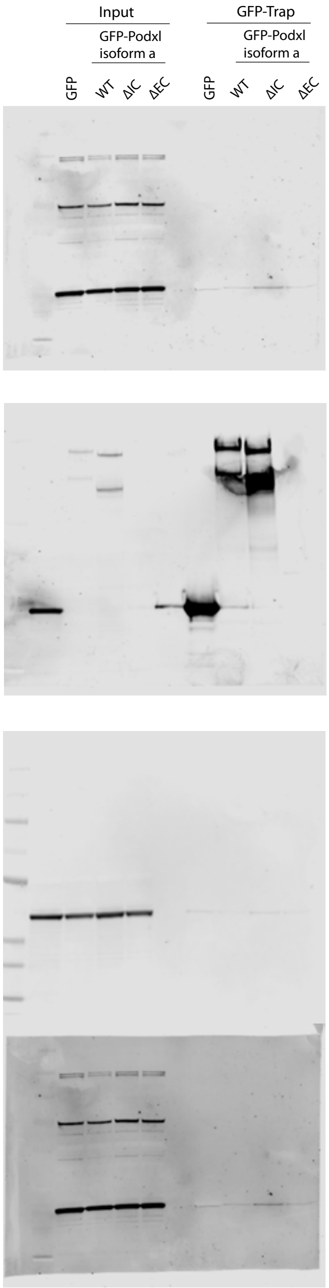


#### Supplementary Figure S7I:

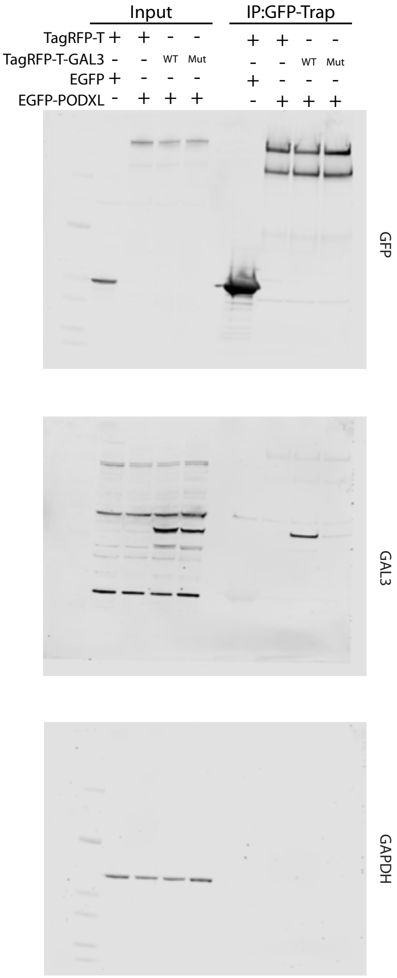


#### Supplementary Figure S7O:

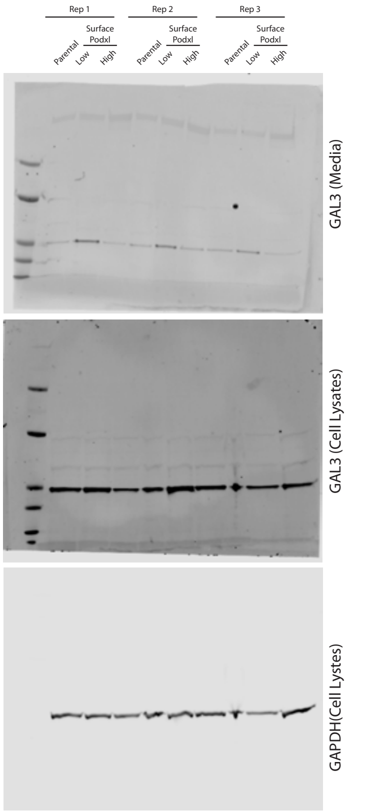


#### Supplementary Figure S8F:

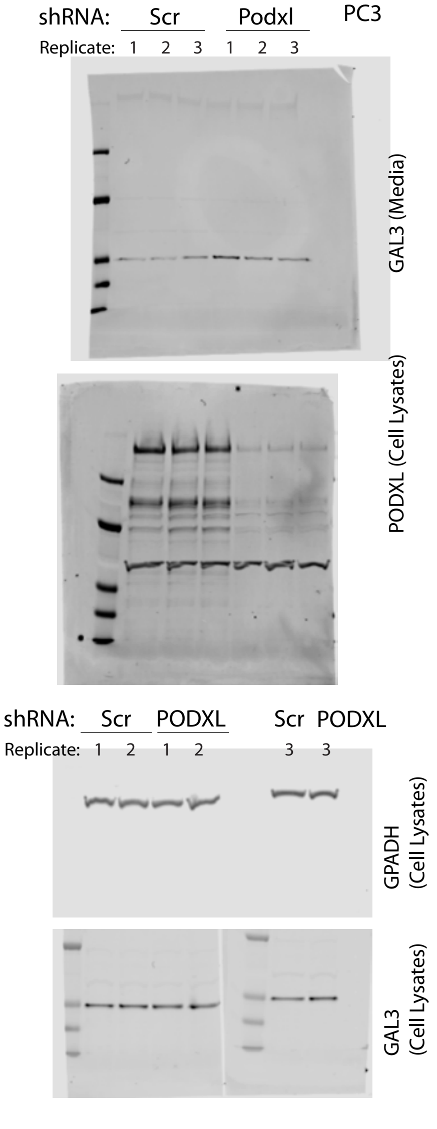


#### Supplementary Figure S9A:

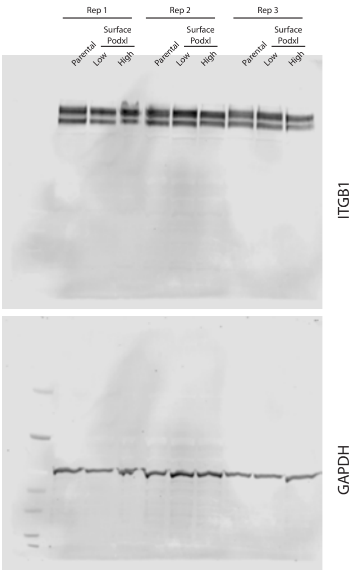
